# A super-pangenome for cultivated citrus reveals evolutive features during the allopatric phase of their reticulate evolution

**DOI:** 10.1101/2024.10.17.618847

**Authors:** Gaetan Droc, Delphine Giraud, Caroline Belser, Karine Labadie, Simone Duprat, Corinne Cruaud, Benjamin Istace, Fredson Dos Santos Menezes, Franck Curk, Gilles Costantino, Alexandre Soriano, Pierre Mournet, Alexis Dereeper, Maëva Miranda, Elodie Marchi, Sylvain Santoni, Raner Santana, Stéphanie Sidibe-Bocs, François Luro, Nathalie Choisne, Florian Maumus, Barbara Hufnagel, Fabienne Micheli, Patrick Wincker, Jean-Marc Aury, Arnaud Lemainque, Patrick Ollitrault

## Abstract

Most of the rich genetic diversity observed in cultivated citrus results from a reticulate evolution involving four ancestral taxa, *C. medica*, *C. reticulata*, *C. maxima* and *C. micrantha*, whose radiation occurred in allopatry. In such an evolutive context, genome diversity studies, GWAS analysis and transcriptomic studies will be significantly enhanced through pangenome approaches. We report the implementation of a super-pangenome for cultivated citrus, established with *de novo* assemblies in pseudochromosomes of *C. medica* (346 Mb), *C. reticulata* (332 Mb) and *C. micrantha* (333 Mb), released for the first time alongside a previously published chromosome-scale assembly of *C. maxima*. Gene annotation of these four genome assemblies revealed 28,090, 29,477, 29,258 and 30,101 genes, respectively, with a focus on pattern recognition receptors. Repetitive element annotation revealed that nearly half of each genome consisted of transposable elements or DNA-satellites. The 3 new genome assemblies display strong synteny and collinearity, while significant discrepancies are observed with the *C. maxima* assembly. Resequencing information (single nucleotide polymorphisms [SNPs], small indels and gene presence-absence variation [PAV]) from 55 accessions were used to explore the intra- and interspecific diversity of the four ancestral taxa and their relationships with the main horticultural groups resulting from reticulate evolution. Diagnostic SNPs of the ancestral taxa, all over the four genomes assemblies, revealed interspecific introgressions in several accessions representative of *C. reticulata*, *C. maxima* and *C. medica* as well as insights into the origin and phylogenomic structures of modern horticultural groups. PAV analysis revealed a gene whose absence or presence was specific to one of the ancestral taxa (dPAV). Interestingly, diagnostic PAV (dPAV) analysis uncovered a large chloroplastic introgression in chromosome 4 of *C. medica*, inherited in horticultural groups and recent hybrids having this species as the male parent. Implementing the super-pangenome combined the identification of orthologous genes across the four genome assemblies and the PAV information from the resequencing data. Significant inter- and intraspecific variations were highlighted, including 25,291 core, 2,431 soft-core and 5,171 dispensable genes. The analysis of the functional enrichment and species-specific adaptations in the citrus super-pangenome revealed distinct functional specializations. This highlights the evolutionary paths that have shaped each species, contributing to the diversity in the citrus super-pangenome while maintaining a shared foundation of essential biological processes. The development of this super-pangenome is a significant milestone in citrus genomics, providing a comprehensive resource that captures the extensive genetic diversity of modern cultivated citrus. Additionally, we established a dedicated Genome Hub, offering a platform for continuous genomic research and allowing for ongoing updates and future inclusion of additional ancestral species.

## INTRODUCTION

Most modern citrus varieties have arisen from reticulate evolution involving four ancestral taxa that evolved in allopatry (Swingle and Reece, 1967; Wu et al., 2018). The radiation period of Asian *Citrus* species was estimated to be in the late Miocene (6–8 million years ago), corresponding to a significant climate transition from wet to drier conditions (Wu et al., 2018). *Citrus medica* L. originally evolved in northeastern India and neighbouring regions of Burma and China. *Citrus reticulata* Blanco originated in China and diversified between China and Japan. The Malay Archipelago and Indonesia have been home to the evolution of *Citrus maxima* (Burm.) Merr. whereas *Citrus micrantha* Wester is native to the Philippines. Hybridizations between these ancestral taxa followed this initial allopatric differentiation, giving rise to the major modern horticultural groups (Figure 1; Curk et al., 2016; Nicolosi et al., 2000; Wu et al., 2018). Among them, mandarins, pummelos and citrons are derived mainly from *C. reticulata*, *C. maxima* and *C. medica*, respectively. However, WGS and GBS studies revealed evidence of *C. maxima* and *C. reticulata* introgressions in cultivated mandarins and some pummelos, respectively (Oueslati et al., 2017; Wu et al., 2018).

**Figure 1:**
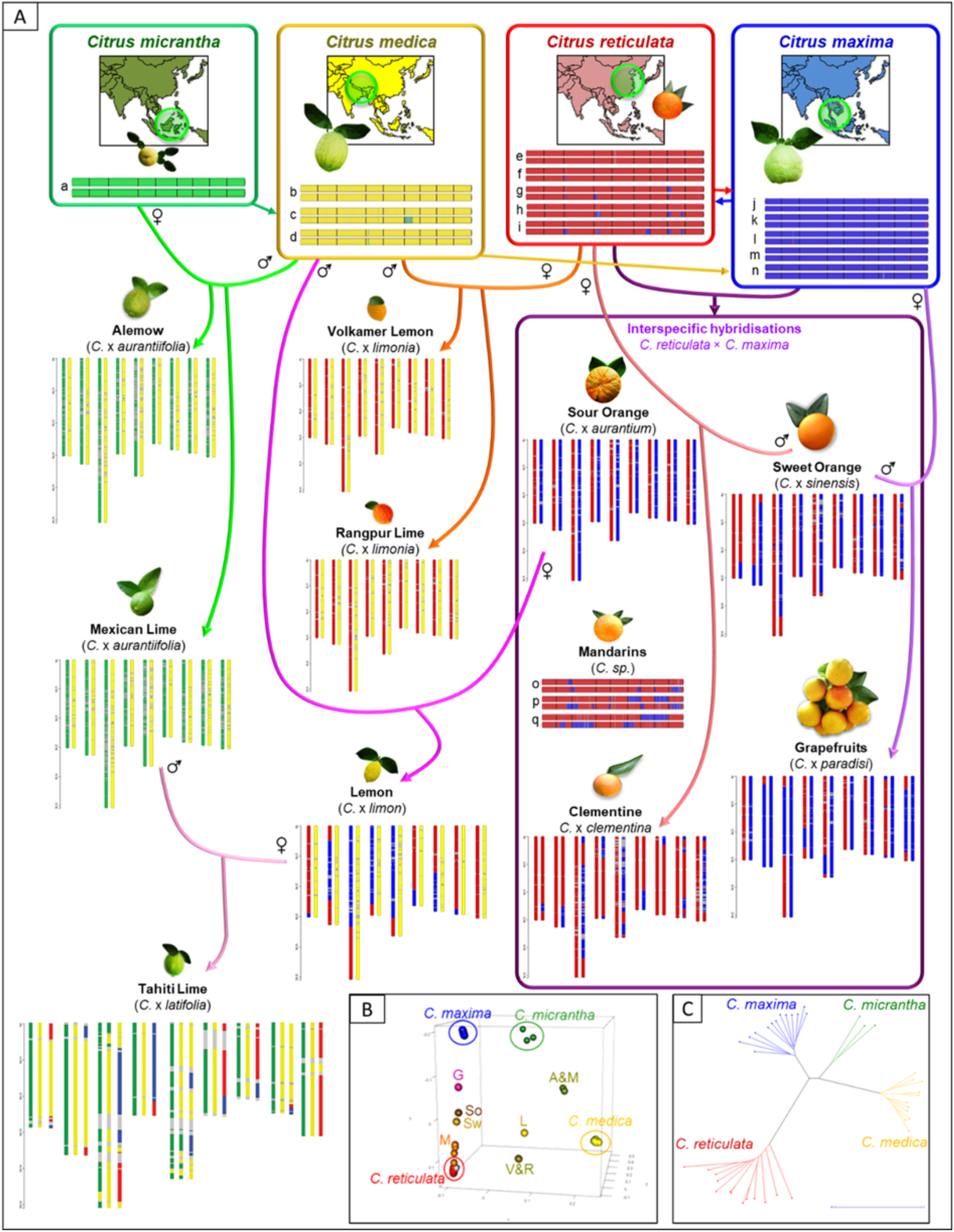
The reticulate evolution of the main horticultural groups of cultivated citrus and their interspecific mosaic structures. A: origin and phylogenomic karyotypes of *Citrus* species derived from the four considered ancestral taxa (a: pap-macrD, pap-melaD, pap- mic1D; b: co-ed, cit-cor1D, cit-diamD, cit-humpD, cit-macnD, cit-macvD, cit-wc1D, cit-wc2D, cit-wc3D; c: cit-chinD; d: cit-e861D; e: man- daoxD, man-HezWD, man-JiaWD, man-ManWD, man-scshD, man-sunkD, man-tachD; f: man-clo1D, man-shekD; g: man-nfmcD; h: man- DaoW1D, man-szinD : i: man-avanD; j: pum-ChonD, pum-florD, pum-hf22D, pum-HuazD, pum-indeD, pum-reinD, pum-tahiD; k: pum-HuanD; l: pum-cha2D, pum-chaH, pum-kaopD; m: pum-GuilD; n: pum-timoD; o: man-ponkD; p: man-kingD; q: man-owarD [correspondence between codes and variety names in Supporting data Table 1]) B: factorial analysis displaying the intermediate position of the horticultural groups resulting from reticulate evolution (G: grapefruits, So: Sour oranges; Sw: Sweet oranges; M: Cultivated mandarins; A&M: ‘Alemow’ and ‘Mexican’ limes; L: Mediterranean lemon); C: diversity of the representative accessions of the four ancestral species (neighbour-joining tree based on Manhattan distances). A, B and C analyses were conducted using CITRE as the reference genome.

Facultative apomixis, through polyembryony of nucellar origin, is present in several *Citrus* species (Wang et al., 2018, 2022b, 2017) and has constrained the number of interspecific recombination cycles. This resulted (Figure 1) in the fixation of genomic structures either in complete interspecific heterozygosity, such as for the sour orange (*C.* x *aurantium* L. = *C. maxima* x *C. reticulata*), ‘Mexican’ lime and ‘Alemow’ (*C.* x *aurantiifolia* [Christm.] Swing. = *C. micrantha* x *C. medica*), Rangpur lime and Volkamer lemon (*C.* x *limonia* Osb. = *C. reticulata* x *C. medica*), or in interspecific mosaics combining large genome fragments from different species (Wang et al., 2018; Wu et al., 2014). These mosaics may involve two species, such as sweet orange or pummelo (respectively *C.* x *sinensis* L. Osb. and *C.* x *paradisi* Macf. resulting from *C. reticulata* / *C. maxima* admixture), or even three ancestors, such as Mediterranean lemon (*C.* x *lemon* [l.] Burm.) resulting from admixture of *C. medica*, *C. reticulata* and *C. maxima*. There can even be four ancestors, such as for the ‘Tahiti’ lime tree (*C.* x *latifolia* Tan. = *C. reticulata*/*C. micrantha*/*C. medica*/*C. maxima* admixtures; Ahmed et al., 2019).

Phenome analysis has shown that a large part of the actual phenotypic diversity in cultivated citrus results from the differentiation between these four ancestral species during allopatric evolution. This is true for the content of some important secondary metabolites such as coumarins, furanocoumarins (Dugrand-Judek et al., 2015) and carotenoid (Fanciullino et al., 2006), as well as pomological and adaptation traits (Santini et al., 2012; Wu et al., 2018). Therefore, the ancestral species and admixed horticultural groups (i.e., oranges, grapefruits, lemons and limes) constitute fundamental genetic resources to breed new citrus varieties. Thus, it is essential to decipher the determinants of the traits supporting the cultivated ideotypes of modern *Citrus* and the variability between these four ancestral taxa.

In this context of reticulate evolution, genome diversity studies, GWAS analysis and transcriptomic studies will be significantly improved through pangenome approaches. Pangenomes limit the biases of analyses based on a single species reference genome (Markello et al., 2022; Stevenson et al., 2013), and pangenome graphs should be particularly useful for allele-specific expression studies in interspecific hybrids and admixed genotypes (Salavati et al., 2019; Sibbesen et al., 2022). Recent studies in plants were conducted to capture a wide range of genome diversity in species or genera (Guo, 2023; Li et al., 2023; Liu et al., 2020) and to decipher the genomic control of some traits, such as fruit quality (Ding et al., 2023; Gao et al., 2019; Sun et al., 2020; Yang et al., 2023; Yocca et al., 2023). Few pangenomic studies on citrus have been published (Gao et al., 2023; Huang et al., 2023; Liu et al., 2022; Zia et al., 2022), and none encompass the whole diversity of cultivated citrus because they miss *C. micrantha* and the integration of intraspecific variability.

Here, we report the implementation of a super-pangenome for cultivated citrus, established for the first time with *de novo* assemblies in pseudochromosomes of *C. medica*, *C. reticulata* and *C. micrantha*, complemented by a published *C. maxima* chromosome scale assembly (Wang et al., 2017). We annotated transposable elements (TEs) and protein coding genes. Special attention was devoted to pattern recognition receptor (PRR) genes, coding for immune system receptors. The definition of core and dispensable genomes integrates (i) the PAV information from resequencing data from 43 accessions representative of the four ancestral species and (ii) a comparative genomics approach based on homology clusters predicted from the four ancestral proteomes. Functional analysis from genes whose absence or presence is specific to one ancestral taxon reveals important evolutionary features during the allopatric phase of cultivated citrus’s reticulate evolution.

## RESULTS

### *De novo* reference genomes for three ancestral species of cultivated citrus

#### Assembly in pseudochromosomes

The estimation of genome size and heterozygosity with Genomescope2.0 from Illumina reads provided the following values (supporting information, Figure 1): 312 Mb (0.9% heterozygosity) for CITMI (*C. micrantha*), 415 Mb (0.242% heterozygosity) for CITME (*C. medica* cv ‘Corsican citron’) and 343 Mb (0.712% heterozygosity) for CITRE (*C. reticulata* cv ‘Cleopatra mandarin’). We produced 42.74 Gb, 32.34 Gb and 52.33 Gb of long-read sequences (Nanopore) for CITMI, CITME and CITRE, respectively (supporting information, Table 2), with N50 values of 18.0 kb, 26.3 kb and 19.7 kb, respectively. The metrics for optical mapping are available in supporting information, Table 3.

The combination of Nanopore long-range sequencing, Illumina PCR free sequencing and optical mapping resulted in 141, 314 and 310 scaffolds encompassing 346.4, 332.9 and 332.1 Mb for the CITME, CITMI and CITRE assemblies, respectively (supporting information Table 4). The Busco score for the three assemblies was high with values for a complete gene over 98.4%.

For the final assemblies in pseudochromosomes, the CITRE and CITME assemblies were anchored on the available genetic maps of Cleopatra mandarin and Corsican citron (Ollitrault et al., submitted). For Cleopatra, the genetic map was saturated and allowed for the establishment of the final CITRE assembly of 9 pseudochromosomes by anchoring 14 scaffolds that represent 89.2% of the whole CITRE assembly (supporting information, Table 5). The synteny between the CITRE assembly and Cleopatra genetic map was high (98.5%) as was collinearity (Spearman coefficient = 0.956 +/− 0.018 on average for the 9 chromosomes; supporting information Table 5, Figure 2a). The anchoring of the Cleopatra genetic map to the CITRE genome revealed a recombination landscape closely associated with gene sequence density along the genome (supporting information, Figure 2c). For Corsican citron, the genetic map presented large gaps due to homozygosity related to the inbreed origin of Corsican citron (Luro et al., 2012). However, sequence anchoring allowed for the integration of 13 scaffolds in 9 pseudochromosomes, encompassing 96.5% of the whole CITME assembly. Synteny (99.2%) and collinearity (Spearman coefficient = 0.935) between the assembly and the partial genetic map were high (supporting information, Table 6, Figure 2b). For CITMI, no genetic map was available, and the synteny and collinearity with the *C. clementina* V1.0 assembly (Wu et al., 2014) drove the final assembly in pseudochromosomes (supporting information, Figure 3). Twenty-one scaffolds were included in the 9-pseudochromosome assembly, representing 90.1% of the whole CITME assembly (supporting information, Table 7). High synteny and collinearity were observed (supporting information, Figure 4) between our 3 genome assemblies and the 2 sweet orange haplotypes published by Wu et al. (2022), *C. trifoliata* (Peng et al., 2020), *C. hindsii* and *C. australis* (Nakandala et al., 2023) genome assemblies. The 2 major discrepancies systematically observed between *C. clementina* V1.0 assembly Chr 3 and Chr 5 and our 3 new assemblies (Supporting information Figure 4a) were previously described in genetic mapping studies and attributed to local mistakes in *C. clementina* assembly (Ollitrault et al., 2012, submitted).

#### Chloroplast genome assemblies

The chloroplast genome assemblies of CITRE, CITME and CITMI encompass 160,601 bp, 160,016 bp and 159,936 bp, respectively. They contain, respectively, 97, 91 and 95 genes, and each assembly display 10 rRNAs and 37 tRNAs (supporting information, Table 8). Theses genomes exhibit the classical chloroplast structure, featuring 2 copies of an inverted-repeated region that separate large and small single-copy regions (Figure 2). The phylogenic position of the 3 new assembled genomes is consistent with previously published chloroplast genomes for *C. reticulata*, *C. medica* and *C. micrantha* (supporting information, Figure 5). The sequences and annotations of the 3 chloroplast genomes are available on our website, https://citrus-genome-hub.southgreen.fr.

**Figure 2:**
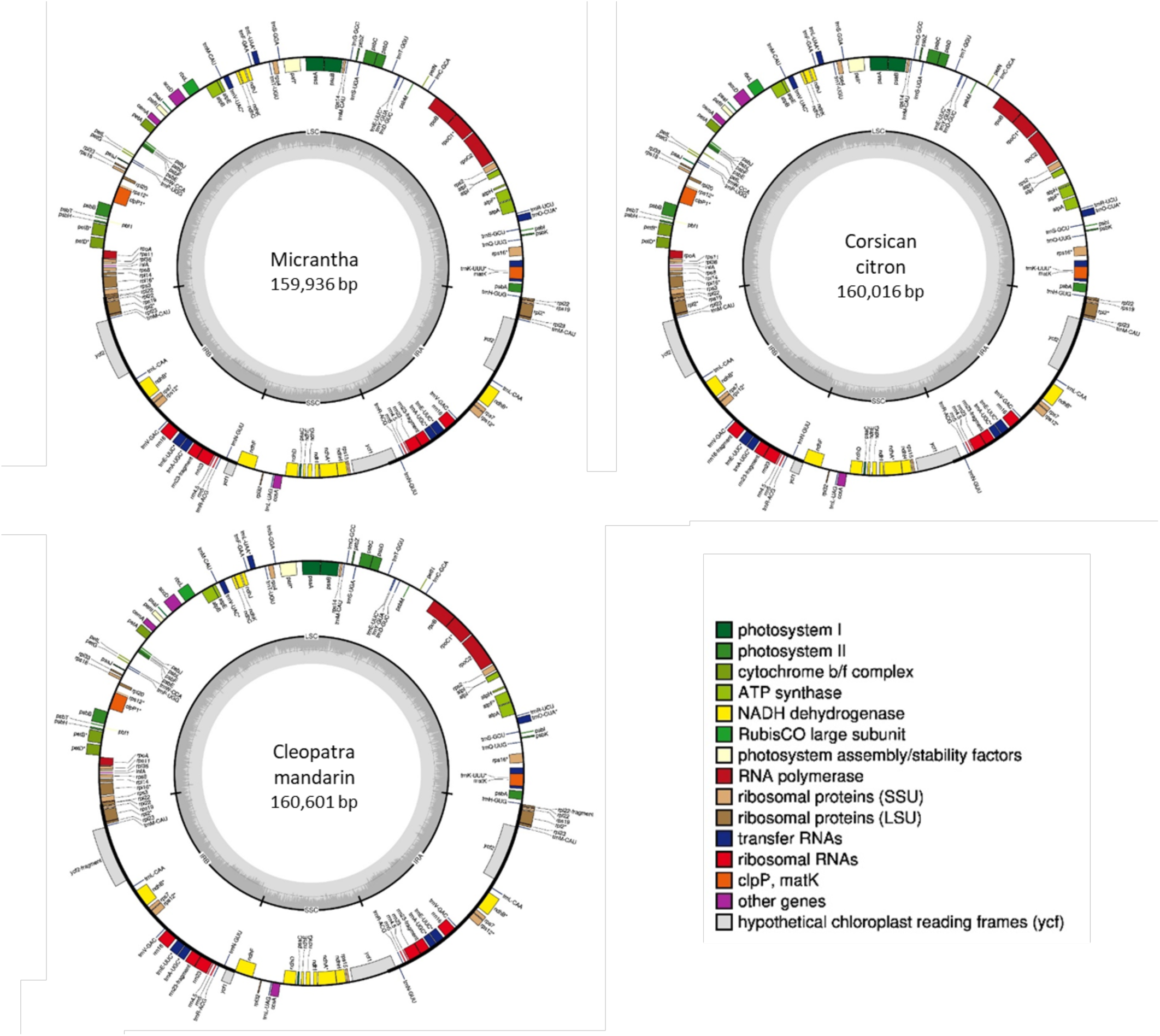
Annotation of the 3 chloroplast genome assemblies.

#### Annotation of the 3 nuclear genome assemblies and comparison with *C. maxima* genome

To accurately compare representative genome assemblies of the four ancestral species, we performed with the same pipe-line the annotation of the 3 new genome assemblies (CITME, CITMI and CITRE) and of the chromosome-scale *C. maxima* assembly published by Wang et al. (2017) and called CITMA in the present paper. Two main citrus chromosome numbering methods are currently used at the international level. They result from the first 2 pseudochromosome assemblies for citrus: the one of *C. sinensis* v1.0 (Xu et al., 2013) used for the *C. maxima* assembly (Wang et al., 2017) and the one of *C. clementina* v1.0 (Wu et al., 2014) that we used for CITME, CITMI and CITRE. In addition, the chromosome orientations appeared to be more diverse (Ollitrault et al., submitted). This inconsistency in chromosome numbering and orientation may result in confusion when one tries to integrate information based on different reference genomes. For synteny and collinearity analyses, we considered a numbering and orientation for *C. maxima* identical to our 3 new genome assemblies.

##### • Genes

RNA-seq was performed with young and mature leaves, entire flowers and roots of the 3 accessions used to establish the *de novo* assembly, with the Chandler pummelo as a representative of *C. maxima*. A total of 307,532 Mb of raw data were produced, corresponding to an average coverage of 561x for each combination accession/organ (supporting information, Table 9).

A total of 30,101, 28,090, 29,258 and 29,477 high-confidence protein coding genes were retained respectively for CITMA (345 Mb), CITME (346 Mb), CITMI (333 Mb) and CITRE (332 Mb) genome assemblies, respectively, by using a combination of homologous proteins, transcriptomic data and *ab initio* prediction. Gene completeness was evaluated with BUSCO v5.0.0 against the eudicots_odb10 lineage, yielding high BUSCO scores of 94.8%, 94.7%, 93.3% and 93.1% for CITME, CITMI, CITRE, and CITMA, respectively. The number of genes per chromosome along with gene completeness is detailed in supporting information Table 10. Functional annotations were performed based on InterProScan domain searches and similarity searches using the UniProt/SwissProt and UniProt/TrEMBL databases (BlastP). Gene ontology (GO) terms were derived from InterProScan results (Jones et al., 2014), and enzyme classification (EC) numbers were predicted using KofamScan (supporting information, Table 11). Gene density varies along the chromosomes with similar patterns for the homologous chromosomes of the 4 assemblies (Figure 3).

**Figure 3:**
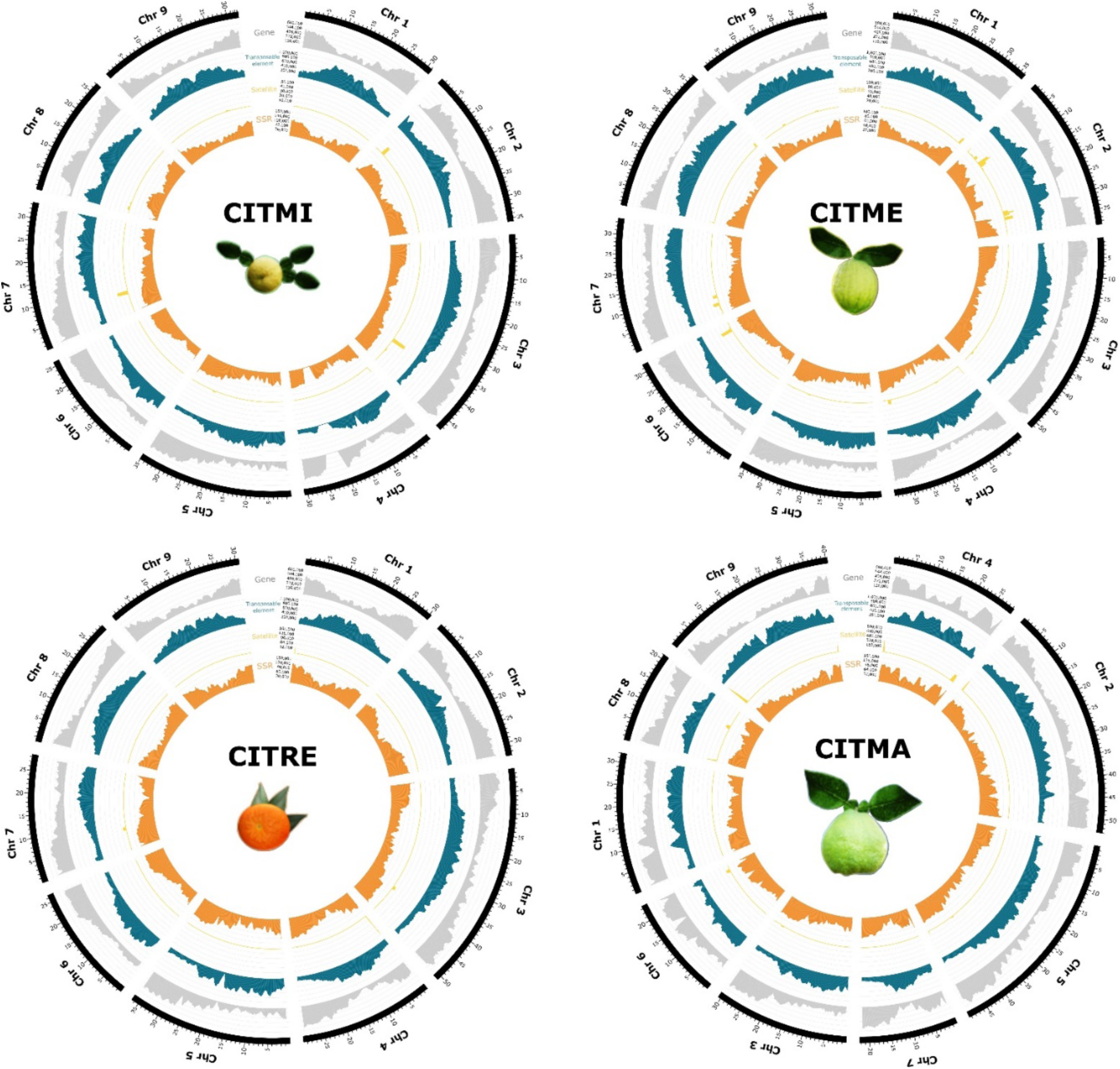
Density of genes (grey histogram), transposable elements (blue histogram), DNA satellites (yellow histogram) and SSRs (orange histogram) along the 9 assembled pseudo-chromosomes of the 4 *Citrus* ancestors. Density was calculated with a 1-Mb sliding window. CITMA chromosomes are numbered per, but ordering and orientationin the figure follows that of the 3 other genome assemblies.

##### • Transposable elements and DNA-satellite sequences

Transposable elements (TEs) constitute a significant portion of the analysed genomes, with 42.8% in CITMI, 43.6% in CITRE, 47.3% in CITME and 42.5% in CITMA (supporting information, Figure 6, Table 12). Most of them (74–77%) are LTR-retrotransposons, including *Copia Tork*, *Sire* and *Retrofit* elements (comprising 12.6% to 15.2% of genomes across species) as well as *Gypsy, Athila* and *Tat* elements (comprising 12.0% to 15.0% of genomes) (supporting information, Figure 7). The higher TE content in *C. medica* compared to the other 3 ancestors (164Mb vs. 142Mb–147Mb) was explained by an accumulation of *Gypsy, Athila* and *Copia Sire* elements in the former. Analyses of intraspecific variability also confirmed the significantly larger amount of TEs in *C. medica* (t-test with p-value < 0.001) (supporting information Figure 8). Additionally, we estimated that approximately 10% of genomes consisted of Class II elements, predominantly TIR elements such as *Mutator*, *hAT* and *CACTA*. Non-autonomous MITE elements constituted from 1.70% to 1.92% of genomes in *C. medica* and *C. reticulata*, respectively. Analysis of chromosome distribution indicated that TEs were dispersed throughout the chromosomes but were more concentrated in specific regions (Figure 3). Such concentrated regions are generally central, but in some chromosomes, such as Chr 6 for all 4 ancestors, TEs are notably accumulated in one chromosome side. In addition, we identified 2 distinct DNA-satellite sequences in the four species under study, accounting for 0.53% of the *C. micrantha* genome to 1.95% of the *C. reticulata* genome. Both DNA satellites, 181 bp and 141 bp in size, are tandemly repeated mainly in chromosome extremity. The 181-bp satellite varies in abundance across species, with 1.7 Mb in *C. micrantha*, 2.8 Mb in *C. maxima*, 5.3 Mb in *C. medica* and 6.4 Mb in *C. reticulata*.

### Nuclear genome diversity in cultivated Citrus according to individual reference genomes

#### Structural variation between chromosome scale genome assemblies of the 4 ancestral species

A strong synteny and collinearity is observed for the 3 new *de novo* assemblies, with the exception of an inversion in the central region of the CITME Chr 2 when compared to CITRE and CITMI (Figure 4). In addition to different numbering and orientations of the CITMA pseudo-chromosomes compared with the 3 other assemblies, the riparian plot reveals many structural differences between CITMA and the 3 other assemblies. It concerns non-syntenic blocs, inverted regions and blocs in different positions in one chromosome.

**Figure 4:**
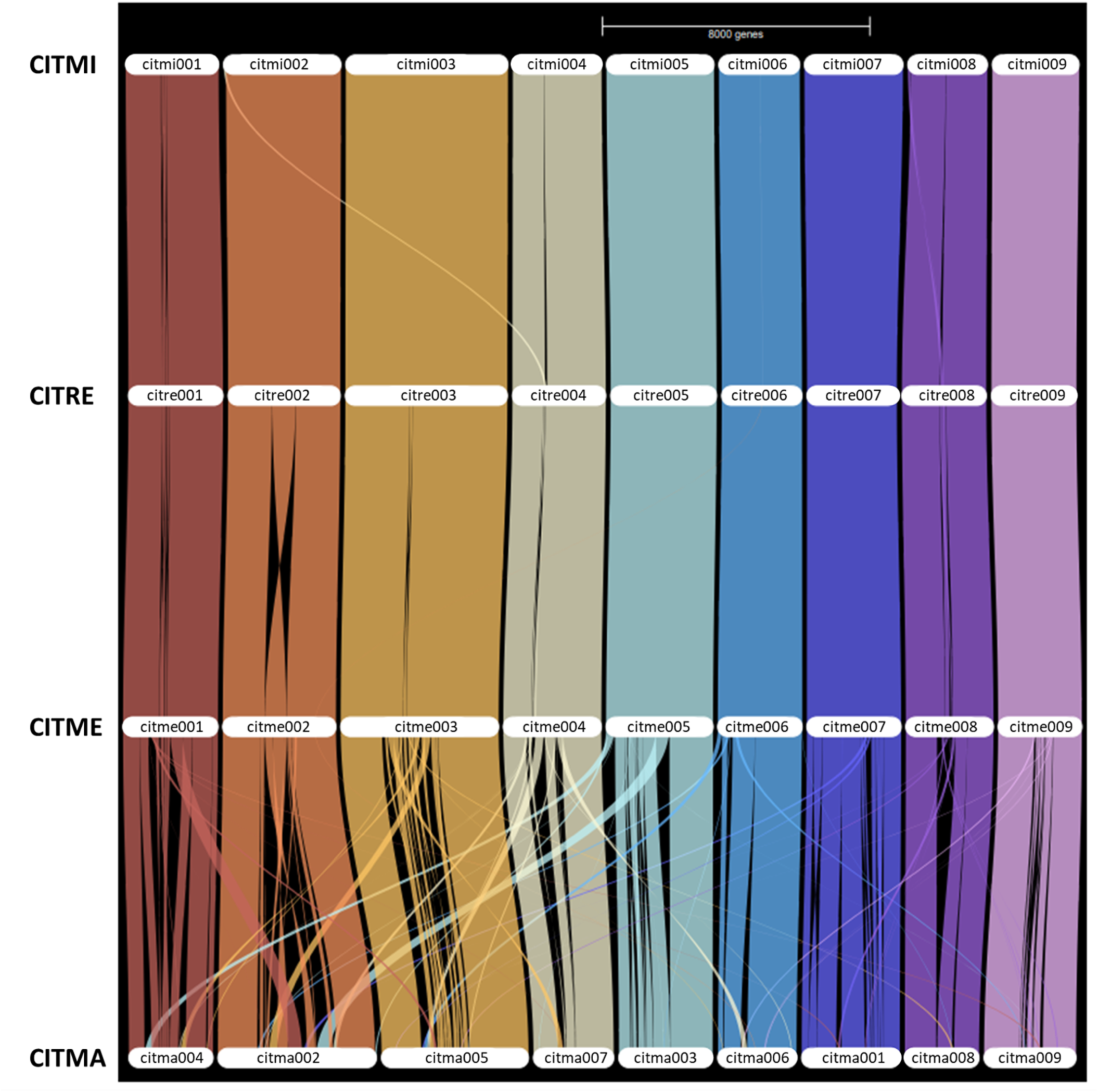
Riparian plots representing the synteny map of orthologous genes along the 9 citrus chromosomes.

#### PRR genes diversity and organization between the 4 ancestral genomes

The total number of PRRs) and R proteins (named as PRRs in the subsequent paragraphs) varied between 1,627 for CITME and 2,014 for CITRE (supporting information, Table 13). Across the 4 genome assemblies, a substantial portion (average of 64.3%) of the identified PRRs fell into the ‘unknown’ class as previously defined (Santana Silva and Micheli, 2020). Among the identified classes, CNL, NL and RLK had the greatest relative contributions across all 4 genome assemblies (supporting information, Figure 9).

Their combined proportions accounted for 51.9%, 57.4%, 66.4% and 67.5% of the identified PRRs for CITME, CITMI, CITRE and CITMA, respectively. The relative contribution of T, TN and TNL was lower in CITMA (6.2% in total) than in the 3 other assembled genomes (20.3%, 17.3% and 11.5% for the CITME, CITMI and CITRE genomes, respectively). The distribution of PRRs from the 12 identified classes is uneven among the chromosomes (supporting information, Table 14). For the 4 genome assemblies, their number is low in Chr 4, 6, 8 and 9 (from 1.9% in CITMI-Chr4 and CITME-Chr6 to 6.1% in CITMA-Chr9). Chr 3 concentrates the largest proportion of PRRs (over 25%) for CITME, CITMI and CITRE whereas the larger proportion for CITMA is found in Chr 2 (28.7%). For the 4 genome assemblies, a large proportion of PRRs (from 19.2% for CITME to 24.7% in CITMI) was also observed in Chr 7 and to a lesser extent in Chr 5 (ranging from 13.0% in CITMI to 21.7% in CITRE).

For these 2 chromosomes, the predominant class is CNL in the 4 genomes (Figure 5). RLK is the predominant class in Chr 2 of the 4 genomes whereas TNL are concentrated in Chr 3 for CITME, CITMI and CITRE. Cluster analysis, including the PRR genes of the unknown class, revealed that a very large proportion (ranging from 85.5% in CITRE to 87.2% in CITMA) of the PRR genes were organized in clusters (supporting information, Table 15). Depending of the genomes and chromosomes, the clusters range from 70% to 95.3% (CITMA-Chr 3 and CITMI-Chr 7, respectively). The number of clusters is similar for all 4 genomes, varying from 298 to 332 for CITRE and CITMA, respectively, with an average number of 4.61+/−0.17 genes per cluster. The rates of duplication events appear to vary among the 4 genomes (Figure 5; supporting information, Table 16). It is more frequent in CITMA, with 46 duplications, mainly involving the CNL (25), RLK (5) and NL (5) classes. The lowest rate is observed in CITRE, with only 14 duplications, primarily for RLKGNK2 (4) and RLK (3). Sixteen duplication events, primarily involving RLK (6) and RLKGNK2 (3), were observed in CITME whereas CITMI displayed 21 duplication events, with the largest amounts present in CNL (6) and RLK (5). Most duplications appear to be interchromosomic for the 4 genome assemblies (Figure 5).

**Figure 5:**
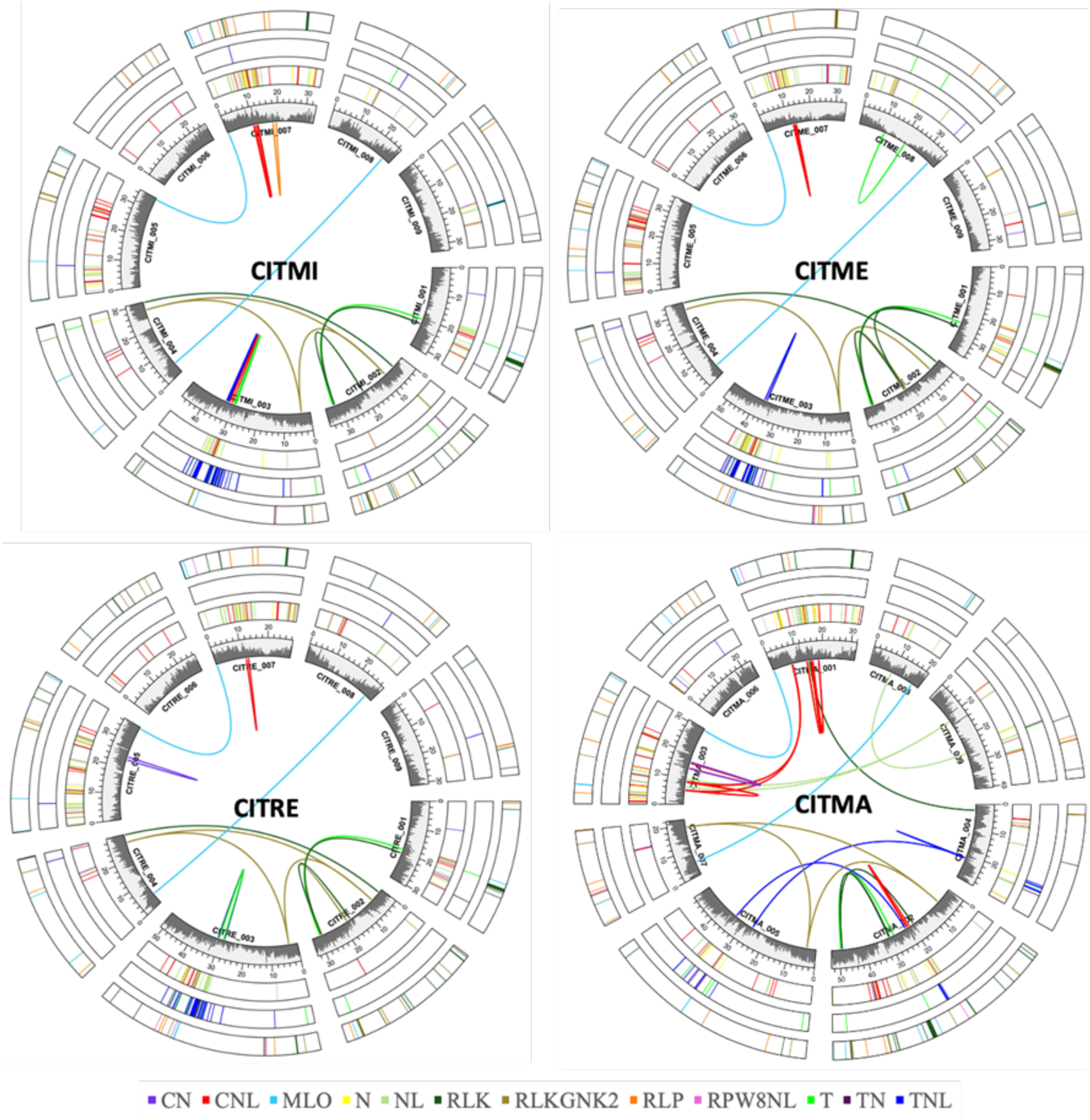
Distribution of PRRs and/or R genes in the 12 classes previously defined (Santana Silva and Micheli, 2020) along the 9 chromosomes of the CITMI, CITME, CITRE and CITMA genome assemblies. Grey histograms: gene density. CITMA chromosomes numbered per Wang et al. (2017), but ordering in the figure follows that of the 3 other genome assemblies.

#### Intra- and interspecific diversities based on resequencing data of 55 accessions of cultivated citrus

##### SNP and small indel variation

The intraspecific diversity was analysed for each genome assembly using the corresponding representative accessions (supporting information, Table 17, Figure 10). It reveals a lower polymorphism for *C. medica* than for the other species. The total number of polymorphisms (SNPs and indels) was 3,924,735 for *C. medica* whereas it varies between 6,905,527 (*C. maxima*) and 8,471,249 (*C. reticulata*) for the others species. The average heterozygosity of the *C. medica* accessions was also very low (0.95 heterozygous sites per kilobases) when compared with the other species (5.10, 4.17 and 3.70 for *C. micrantha*, *C. reticulata* and *C. maxima,* respectively). The proportion of indels over the whole variants is remarkably similar (10.78+/−0.32%) for all 4 species. Analysis of all representatives of the 4 species on each genome assembly yielded similar results regardless of the reference genome, with 19,633,997+/−501,376 SNPs and 3,006,432+/−122,037 and a proportion of indels (13.28+/−.22%) slightly increasing compared to intraspecific data. There was a limited increase in total diversity (24,169,356+/−760,674) when admixed accessions were included, with no significant change in the indel rate relative to whole polymorphism (13.46+/−0.22). The diversity of the representative accessions of the 4 ancestral species represents, respectively, 92.75%, 93.68%, 94.04% and 94.27% of the total diversity for the CITRE, CITME, CITMA and CITMI genome assemblies, with interspecific diversity constituting the main component (77.94%, 65.21%, 62.20% and 58.32% for the variant calling using CITME, CITMA, CITMI and CITRE, respectively; supporting information, Figure 10). Admixed accessions displayed a logical, significant increase in heterozygosity rate (on average 3.43 times higher) compared to accessions representative of the ancestral species. Phylogenetic trees of the representative accessions of the 4 ancestors and the factorial analysis of the whole set of accessions (Figure 1C and 1B, respectively, for example, for the CITRE reference and supporting information, Figures 11 and 12) confirm this high level of interspecific differentiation and the strong structuration between the ancestral species. Admixed accessions are located in the factorial analysis according to their ancestors’ contribution.

##### Introgressions and admixture in cultivated citrus

The first step was to identify SNPs that fully differentiated one species from the 3 others. These diagnostic SNPs (dSNPs) of the 4 ancestors were mined in the 4 genome assemblies. A total of 2,227,454, 2,298,439, 2,170,653 and 2,271,482 dSNPs were found for the CITMA, CITME, CITMI and CITRE genome assemblies, respectively (supporting information Table 18). The DSNP lists for the 4 genome assemblies are downloadable at https://citrus-genome-hub.southgreen.fr. DSNPs represent, respectively, 11.38%, 11.39%, 11.44% and 11.47% of the total number of SNPs found for the set of representative accessions of the 4 ancestral species, and they are dispersed all along the genome (supporting information, Figure 14). Across all 4 assemblies, the relative proportion of dSNPs for each ancestral species was similar, with a majority of *C. medica* dSNPs (47.43+/−0.88 %) and a minority for *C. maxima* (13.90+/−0.81 %) and *C. micrantha* ones (12.75+/−1.52 %), with *C. reticulata* displaying intermediate values (25.90+/−1.01%). These results are consistent with the relative differentiation between the 4 corresponding horticultural groups that the phylogenomic analysis revealed (Figure 1C). The admixture patterns established using the CITME, CITMI and CITRE assemblies are very similar for all re-sequenced accessions (supporting information, Figures 15, 16, 17 and 18). However, the results from CITMA (supporting information, Figure 17) differ slightly from those of the 3 other assemblies due to structural differences between CITMA and the others assemblies. Nonetheless, the same conclusions are obtained from the analysis on the 4 reference genomes. Figure 1A shows the interspecific mosaic structures revealed using CITRE as a reference. The mandarin horticultural group displays a predominance of the *C. reticulata* genome but also shows frequent introgressions from *C. maxima*. Seven of the analysed pummelos appear to be pure representatives of *C. maxima* whereas 6 display limited introgressions. For 3 varieties (‘Chandler’, ‘Kao pan’ and ‘Chandler haploid’), it consists of 1.2-Mb *C. reticulata* introgressions in Chr 2. *Citrus reticulata* introgressions are also observed in Chr 9 spanning 0.5 Mb and 1.5 Mb for ‘Guil’ and ‘Huan’ pummelos, respectively. The ‘Flores’ pummelo displays a 0.5-Mb *C. medica* introgression. Of the 11 citrons, 9 are pure representatives whereas the remaining 2 varieties present *C. micrantha* introgressions; large (19 Mb) and small (0.4 Mb) ones in Chr 6 and 4, respectively, for ‘Chinese’ citron; and a limited one (1.3 Mb) in Chr 4 for ‘Etrog’ citron s861. Of the horticultural groups resulting from admixture, sour orange displays a full *C. maxima/C. reticulata* heterozygosity all along the genome. Sweet orange and grapefruit present more complex admixture structures between the same ancestors, with heterozygous and homozygous regions. These 3 groups have a *C. maxima* chlorotype (supporting information, Figure 19). ‘Alemow’ and ‘Mexican’ limes display similar phylogenomic structures with complete *C. micrantha/C. medica* heterozygosity and *C. micrantha* chlorotype. The Volkamer lemon had a full *C. reticulata/C. medica* heterozygosity and a *C. reticulata* chlorotype. The Mediterranean lemon displays the more complex phylogenomic structure with a mosaic of *C. reticulata/C. medica* and *C. maxima/C. medica* heterozygous regions and a *C. maxima* chlorotype.

#### Gene PAV according to each reference genome

To analyse the intra- and interspecific diversity of PAVs, we adopted a methodology similar to the one used in the search for DSNPs. Regions of the representative accessions of each species found to be introgressed in previous admixture studies were excluded from the analysis. *Citrus medica* appeared to be the least polymorphic species, with only 1,953 polymorphic genes, whereas *C. reticulata*, *C. maxima* and *C. micrantha* have 4,577, 4,691 and 3,568 polymorphic genes, respectively (Supporting information Table 19).

At the interspecific level, including the admixed horticultural groups, the percentages of core genes found in the CITRE, CITMA and CITMI genomes are very similar (67.6, 67.3 and 67.4%, respectively) and slightly higher for CITME (71.7%). For each genome assembly, we identified genes that were specific to the corresponding species (i.e., found in all accessions of the species and absent of all accessions of the three other species) and genes specifically absent from one of the other ancestral species (absent in all accessions of one species and present in all accessions of the three other species). The number of these diagnostic genes (dPAV) varied from 1,029 for CITMI to 1,438 for CITME, with similar values for CITRE and CITMA (1,158 and 1,148, respectively). Several genes annotated in one of the four genome assemblies were missing in all accessions of another ancestor. *Citrus medica* is the species displaying the largest number of such missing genes in the three other species’ genome assemblies (respectively, 527, 522 and 435 in CITMA, CITMI and CITRE; Supporting information, Table 19). The average number of genes missing for *C. reticulata, C. maxima* and *C. micrantha* in the genome assemblies of the three other species is 184 (average for CITMA, CITME and CITMI annotated genes), 109 (CITME, CITMI, and CITRE) and 139 (CITMA, CITME, and CITRE), respectively. CITME also has the largest number of specific genes of *C. medica* (990) whereas 497, 295 and 210 are found in CITRE, CITMA, and CITMI, to be specific, of *C. reticulata*, *C. maxima* and *C. micrantha*, respectively (supporting information, Table 19). For the four reference genomes, the neighbour-joining (NJ) trees based on PAV displays a strong differentiation between species (supporting information, Figure 20) with increased branch lengths of the corresponding species, according to the contribution of specific genes to the interspecific differentiation.

Although the distribution of the number of diagnostic PAVs along the CITRE, CITMI and CITMA genomes is relatively homogenous, the CITME genome displays a significant peak of diagnostic PAV in the 17-18-Mb interval of Chr 4 (supporting information, Figure 21). This region includes 97 dPAVs specific to *C. medica* out of 185 genes. Among the admixture horticultural groups, these genes are found in the direct hybrids of *C. medica* as the male parent (such as ‘Eureka’ and ‘Volkamer’ lemons as well as ‘Rangpur’, ‘Mexican’ and ‘Alemow’ limes), but they are absent in the other admixed horticultural groups (supporting information, Figure 22a). These dPAV genes are annotated as chloroplastic genes. A dot plot analysis of the chloroplastic genomes with CITME revealed homology with the entire chloroplast genome in this CITME Chr4 region, with several duplications (Figure 23 a and e), whereas no homology was found between nuclear and chloroplast genomes for the three other genome assemblies (supporting information, Figure 23b, c and d). Phylogenetic analysis of various genes with chloroplastic homology places the nuclear genes near the chloroplastic *C. medica* ones (supporting information, Figure 24), suggesting an intraspecific introgression of the chloroplastic genome into Chr4 of *C. medica*. However, the nuclear orthologues are generally smaller than the original chloroplastic ones (with the exception of the NDHJ gene), displaying deletions (e.g., MatK gene in supporting information, Figure 25).

To confirm the considered genes’ nuclear location, we performed a new genotype calling with an additional natural or recent admixture of *C. medica* (first- and second-generation hybrids in which *C. medica* or its hybrids were the male parent). The dPAVs of the 17-18-Mb region of Chr 4 are found in all first-generation hybrids. For the second-generation hybrids, the dPAVs are found only in those that inherited the corresponding nuclear genomic region from *C. medica*, as evidenced by the interspecific mosaic analysis (supporting information, Figure 26). Thus, the considered dPAV genes are absent in bergamota, clementine x ‘Nestour’ lime and clementine x ‘Rangpur’ limes, which have no *C. medica* nuclear contribution in this region of Chr 4. Additionally, all first- and second-generation hybrids inherited the chloroplast from their female parent (*C. reticulata* for ‘Rough’ and ‘Volkamer’ lemons and ‘Rangpur’ lime, clementine x ‘Alemow’, clementine x ‘Nestour’ lime, clementine x ‘Volkamer’ lemon and clementine x ‘Rangpur’ lime; *C. micrantha* for ‘Alemow’, ‘Mexican’ and ‘Nestour’ limes; *C. maxima* for ‘Chandler’ pummelo x ‘Corsican’ citron, ‘Chandler’ pummelo x ‘Eureka’ lemon and bergamota; supporting information, Figure 22b). These results validated the nuclear location of the considerate dPAV genes. In the same variant calling, we also expanded the diversity to others *Citrus* ancestral species, such as *C. ichangensis*, *C. indica*, *C. japonica*, *C. trifoliata*, *C. glauca*, *C. australis, C. australasica* and *Atalantia buxifolia*. The dPAVs of the Chr4 17-18 Mb were absent in all these species except *C. indica* (supporting information, Figure 22a), which shared a phylogenetic branch with *C. medica* (supporting information, Figure 22b). Twenty-six of the 97 dPAVs were present in *C. indica*.

### A super-pangenome for cultivated citrus

The cultivated citrus super-pangenome was established by combining the identification of orthologous genes among the four genome assemblies and the PAV information for 55 accessions on the same four genome assemblies (Figure 6).

**Figure 6:**
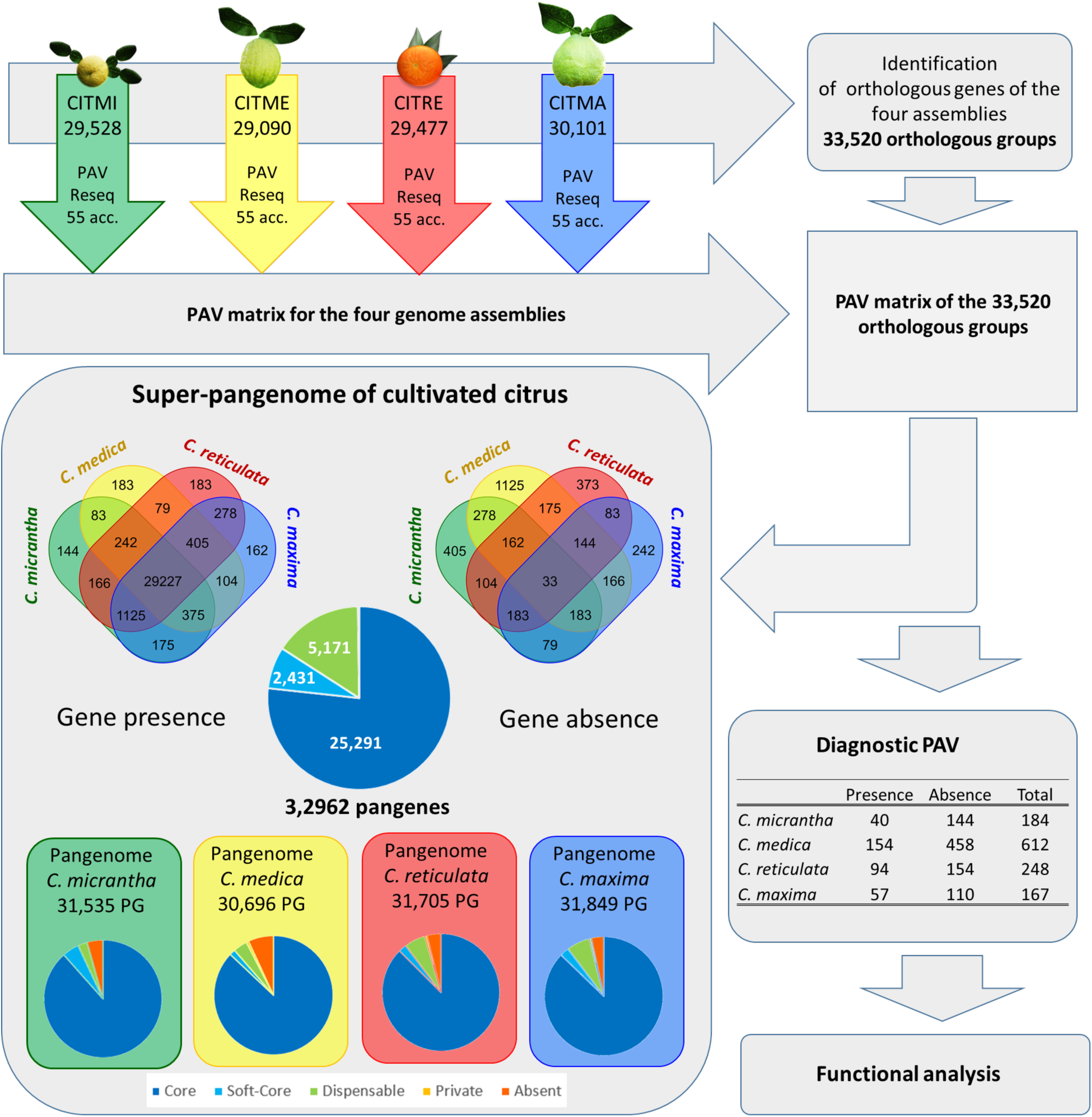
Implementation strategy and structure of the cultivated citrus super-pangenome; acc.: accession.

#### Search for orthologous genes among the four genome assemblies

The number of orthologous groups across the four assemblies is 33,520 (supporting information, Table 20). The number of orthologous groups per assembly ranges from 24,716 for CITMA to 25,322 for CITMI. The proportion of single-copy genes by orthologous group varies from 88.98% for CITRE to 92.10% for CITME (supporting information, Figure 27).

#### PAV Analysis and Core/Dispensable genome identification

The PAV for the pangenes was established by combining PAV information from individual assemblies and the established orthology links (Figure 6). A pangene was considered present in a specific accession if at least one of the corresponding orthologous genes was present in one of the four ancestral genome assemblies.

From the PAV analysis, 558 orthologous groups are found to be absent in all accessions. Therefore, the super-pangenome encompass 32,962 pangenes (considering each orthologous group a pangene). Of the pangenes, 33 are found only in some accessions of the admixed horticultural groups but not in the representative accessions of the four ancestral taxa. NJ tree analysis of the pangenome PAV data indicates a strong structuration among the four ancestral species (Figure 7a). Factorial analysis, based on the 55 accessions (Figure 7b), reveals the positions of admixed horticultural groups, which are fully consistent with their phylogenomic constitution.

**Figure 7:**
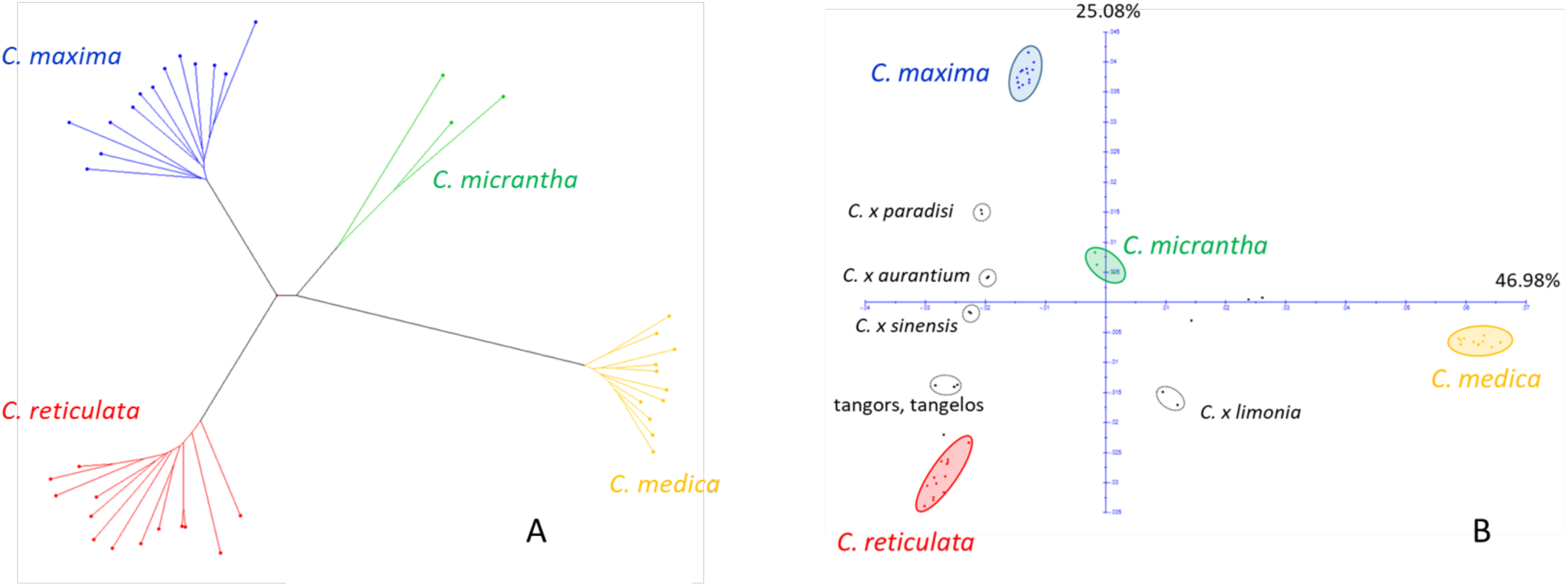
Organization of the PAV for 32,962 validated pangenes. A: Neighbour-joining tree of 43 accessions representative of the four ancestral species; B: Factorial analysis of the 55 re-sequenced accessions.

The number of pangenes varies between 30,696 for *C. medica* and 31,849 for *C. maxima* (supporting information, Table 21). The intraspecific core-gene frequency ranges from 90.2% for *C. maxima* to 93.5% for *C. medica* (Figure 6). At the interspecific level, the validated pangenes encompass 25,291 core pangenes, 2,431 soft-core genes, 5,171 dispensable pangenes and 69 private genes (supporting information, Table 21 and Table 22). A total of 29,227 pangenes are present in at least one accession of each of the four ancestral species (Figure 6).

A total of 1,211 pangenes were found to be diagnostic for one of the four ancestral species (Figure 6; supporting information, Table 23). Among them, the diagnostic pangenes for absences (866) are more frequent than those for presences (345). *Citrus medica* has the largest number of dPAVs, with 612 diagnostic pangenes.

### Functional Enrichment and Species-Specific Adaptations in the Citrus Pangenome

We conducted a gene ontologyGO enrichment analysis to elucidate the functional landscape of the core genome in the citrus super-pangenome, comprising the four ancestral species. The core genome, representing the set of genes shared across all species, revealed significant enrichment in several critical biological processes, including transcription regulation by RNA polymerase II, ubiquitin-dependent protein catabolic processes, stress response mechanisms and the vegetative-to-reproductive-phase transition (Figure 8; supporting information, Figure 28). These functions underscore the core genome’s essential roles in maintaining cellular homeostasis, regulating gene expression and ensuring successful development and reproduction across these citrus species.

**Figure 8:**
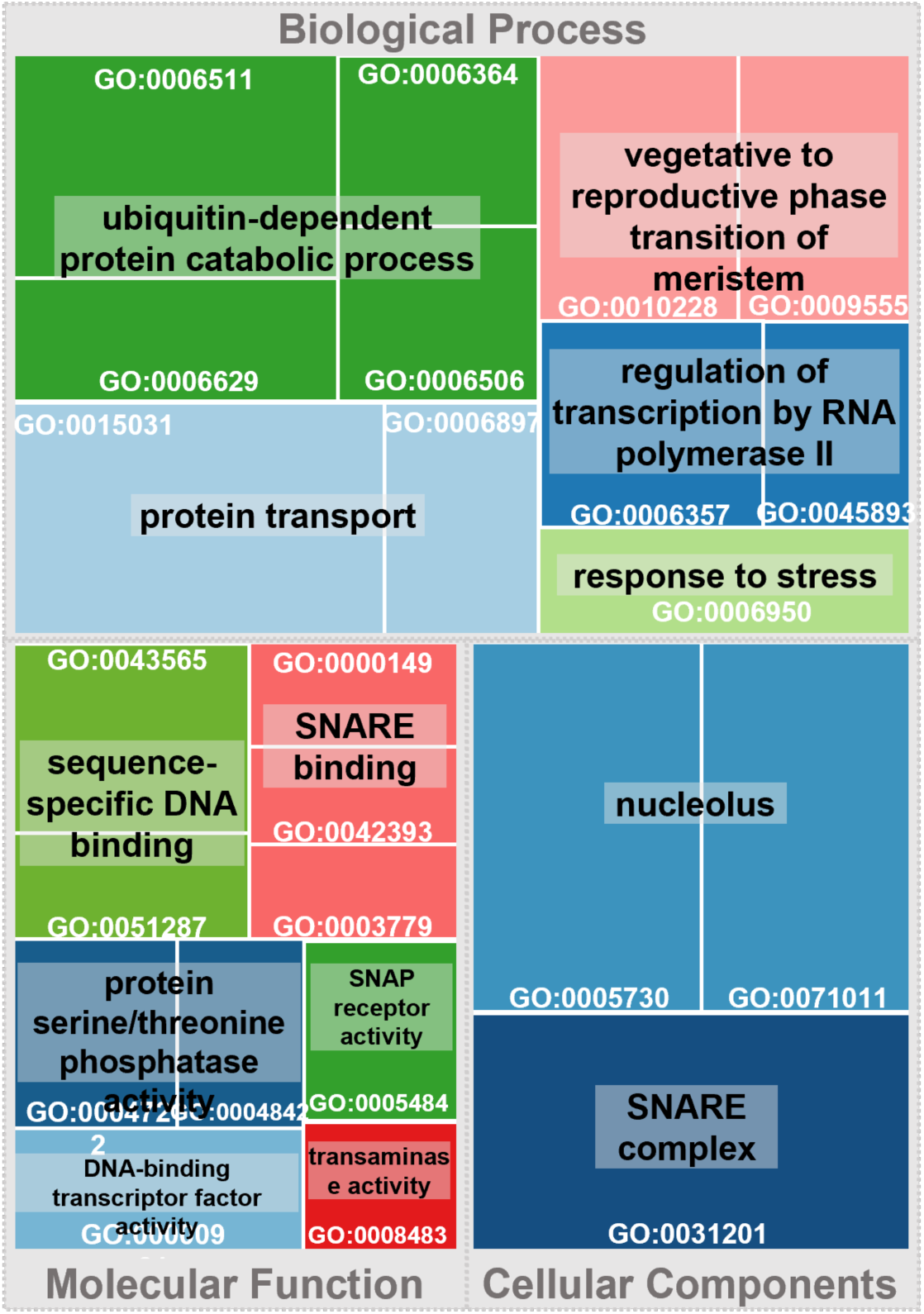
Significant GO terms of the core genome. This tree map represents the enriched Gene Ontology (GO) terms for the core genome genes shared across all species in the citrus pangenome. Each rectangle represents a GO term, with the rectangles’ size proportional to the level of statistical significance of the enrichment (p-value). The colour groups related GO terms into broader categories, illustrating the primary biological processes, molecular functions and cellular components that are conserved in the core genome.

Beyond the core genome, we analysed each species’ unique genes to identify specific functional enrichments that contribute to their distinct adaptive traits. In *C. reticulata*, diagnostic and unique genes related to the auxin-mediated signalling pathway, RNA repair and protein localization are enriched, with potential impact on growth regulation, maintaining RNA integrity and precise protein trafficking (supporting information, Figure 29). These functions are crucial for responding to environmental stimuli, ensuring accurate protein synthesis under stress and maintaining cellular organization. *Citrus maxima* exhibited enrichment in mRNA catabolic processes, autophagy and energy metabolism, reflecting its robust mechanisms for cellular quality control and efficient energy management. These processes are essential for degrading faulty mRNAs, recycling cellular components and sustaining energy production, which are vital under stress (supporting information, Figure 29). In contrast, *C. medica* absent genes showed enrichment in flavonoid and diterpenoid biosynthesis (supporting information, Figure 29), suggesting limitations in secondary metabolite production and stress response, which could affect its ability to defend against pathogens and manage oxidative stress. Meanwhile, *C. micrantha* displayed enrichment in antiviral responses, salicylic acid signalling and RNA processing (supporting information, Figure 29), highlighting its specialized defence mechanisms against viral pathogens and its sophisticated gene expression management. These functions likely provide a significant advantage in environments where pathogen pressure is high. However, *C. micrantha* also lacked genes involved in processes, such as transcription-dependent tethering of RNA polymerase II gene DNA at the nuclear periphery, RNA export from the nucleus, water transport, phylloquinone biosynthesis and sulfation (supporting information, Figure 29). These gene absences suggest potential limitations, which could impact *C. micrantha* varieties’ adaptability and ecological fitness in certain environments.

## DISCUSSION

### Chromosome-scale genome assemblies for C. micrantha, C. medica and pure C. reticulata are provided for the first time

We produced for the first time chromosome-scale genome assembly for *C. medica* and *C. micrantha*. A scaffold scale assembly for *C. medica* was previously published by (Wang et al., 2017) and no genomic resources except resequencing data were available for *C. micrantha.* The first chromosome-scale assemblies for mandarins have recently been released (Isobe et al., 2023). They concern the Satsuma mandarin and its two supposed parents, the ‘Kunembo’ and Kishu mandarins. The Satsuma and ‘Kunembo’ display very large introgressions of *C. maxima* (Oueslati et al., 2017; Wu et al., 2018) and therefore cannot be considered a pure representative of the ancestral *C. reticulata*. The ‘Kishu’ is a Japanese mandarin variety producing small fruits classified as *C. kinokuni* by Tanaka (1961), who considered it a different species from *C. reticulata*; however, its phylogenomic position remain to be clarified. In this work, we presented a chromosome-scale genome assembly for *C. reticulata* established with the Cleopatra mandarin that displays a pure *C. reticulata* phylogenomic ancestry all along its genome, without any interspecific introgression. The three new assemblies did not display any introgression; and, therefore, CITRE, CITME, and CITMI are the first chromosome-scale assemblies of pure *C. reticulata*, *C. medica* and *C. micrantha* genomes, respectively.

These three new assemblies were annotated alongside a previously published chromosome-scale assembly of *C. maxima*. (Wang et al., 2017). 30,101, 29,477, 29,258 and 28,090 high-confidence protein coding genes were identified respectively for the CITMA, CITRE, CITMI and CITME genome assemblies. We focused on resistance genes. PRRs and/or R genes are widely distributed across plant genomes, playing crucial roles in recognizing pathogen-associated molecular patterns (PAMPs) or effectors. This recognition triggers varying responses: PAMP-triggered immunity (PTI) or effector-triggered immunity (ETI) with different intensities and specificities (Jones and Dangl, 2006; Katagiri and Tsuda, 2010). In this study, these genes were identified in four ancestral citrus species: *C. medica* (1,627 genes), *C. micrantha* (1,899), *C. maxima* (1,851) and *C. reticulata* (2,014). Similar numbers were found in other species, such as in *C. sinensis* (2,345), *C. clementina* (1,617) *A. thaliana* (1,756), and others (Santana Silva and Micheli, 2020). The main receptors activating PTI are receptor-like kinases (RLKs), characterized by LRR, transmembrane and kinase domains. Each species showed the following number of RLK genes—*C. medica* (88), *C. micrantha* (97), *C. maxima* (95) and *C. reticulata* (102)—comparable to other citrus and plant species (Santana Silva and Micheli, 2020). For ETI, NBS-LRR genes, specifically CNL and TNL classes, are crucial. The numbers of CNL genes were 139, 182, 227 and 250 for *C. medica*, *C. micrantha*, *C. maxima* and *C. reticulata*, respectively whereas the numbers of TNL genes were 55, 73, 19 and 56 for *C. medica*, *C. micrantha*, *C. maxima* and *C. reticulata*, respectively, with comparisons to other species showing similar distributions (Santana Silva and Micheli, 2020; Wang et al., 2015). These genes tend to cluster on specific chromosomes across citrus species, such as Chr 3, 5 and 7 in *C. medica*, *C. micrantha* and *C. reticulata* and Chr 2, 5 and 7 in *C. maxima*. High concentrations were also observed on specific chromosomes for RLKs (*C. medica* and *C. micrantha*), CNLs (*C. medica* and *C. reticulata*) and TNLs (all species). Additionally, gene duplication events were identified that contributed to genetic diversity and adaptive responses in citrus species. These events play a crucial role in evolutionary processes, potentially enhancing resistance and adaptation in response to environmental stresses (Panchy et al., 2016; Qian and Zhang, 2014).

The annotation and the quantification of repeated sequences in the 4 *Citrus* ancestors revealed that despite their small sizes (from 332Mb to 347Mb by haploid genome), almost half of each genome is composed of transposable elements or DNA satellites (47.8% *C. micrantha*, 50.3% *C. reticulata*, 52.7% *C. medica*, 48.2% *C. maxima*). The larger number of *Gypsy Athila* and *Copia Sire* elements in *C. medica* compared with that in the other three species indicates a specific dynamic of TEs during *Citrus* radiation, particularly since the divergence of the four ancestors. The observed dynamic was congruent with previous studies that have shown the timing of retrotransposon activity and accumulation in *Citrus* species (Borredá et al., 2019; Liu et al., 2019). The number of TEs estimated in each reference genome appears to be representative of all ancestor varieties. However, minor variations identified between varieties reflect the more recent dynamics of TEs, which could explain the diversity of observed phenotypes in each *Citrus* species. In addition, we found that the TE density varied along the chromosomes, with more TEs on one side of Chr 5, 6 and 7 in CITMI, CITRE *and* CITME. Similar distribution patterns were observed for the corresponding *C. maxima* chromosomes of the (Wang et al., 2017) assembly (Chr 3, 6 and 1 respectively). This was already observed in Chr 1, 3, 6 of *C.* x *sinensis* (Xu et al., 2013), using the same chromosome numbering as for *C. maxima* and in Chr 5, 6 et 7 of *C. clementina* (Wu et al., 2014) and with the same chromosome numbering as for CITME, CITMI and CITRE assemblies. Aleza et al. (2015) located the centromere position in the nine clementine chromosomes by half tetrad analysis and determined the acrocentric and submetacentric configuration of Chr 5, 6 and 7. The analysis of the recombination landscape of *C. reticulata* combined with gene density along the genome yielded the same conclusions. Moreover, a comparative genetic mapping study (Ollitrault et al., submitted) revealed a strong similarity between the recombination landscape along the genome of several *Citrus* species and admixed horticultural groups, suggesting a conservation of the centromere location across species. Therefore, TEs were highly concentrated in pericentromeric regions in all chromosomes, but the position of centromeres varies across chromosomes, explaining our observations for TE density. Regarding the DNA satellite, the 181-bp satellite has previously been identified in *C.* x *limon*, *C.* x *sinensis*, *C.* x *aurantium*, *C. ichangensis* and *C. trifoliata* (Fann et al., 2001; Kang et al., 2008). Furthermore, fluorescence in situ hybridization analyses conducted by Deng et al. (2019) confirmed its localization in telomeric and subtelomeric regions across several *Citrus* species, such as *C.* x *clementina*, *C.* x *aurantium*, *C.* x *paradisi* and *C.* x *tachibana*. This satellite sequence appears to be conserved during the diversification of the *Citrus* genus.

### Synteny and collinearity of the ancestral taxa of cultivated Citrus

The synteny and collinearity are high among the three new genome assemblies. The main structural variation concerns an inversion in the centromeric region of the Chr 2 of CITME when compared with CITMI and CITRE. In addition, dot-plot analysis reveals an important synteny and collinearity of these three genomes, with five citrus genome assemblies recently published (Nakandala et al., 2023; Peng et al., 2020; Wang et al., 2022b; Wu et al., 2022). Conversely, the *C. maxima* assembly Wang et al. (2017) published displays numerous differences from the three new assemblies and other published citrus genomes. A recent comparative genetic study at the interspecific level (Ollitrault et al., submitted) revealed a large structural conservation of the genome of the various *Citrus* species, including *C. maxima*. The same authors also suggested probable local assembly mistakes in the *C. maxima* genome published by Wang et al. (2017) that displayed numerous differences from their *C. maxima* genetic map. The genome structural conservation across cultivated *Citrus* species is a favourable situation for further interspecific sexual breeding and for translational genomics.

### Introgressions and admixtures in cultivated Citrus

Regardless of which reference genome is used as a template, the general conclusions from the interspecific mosaic analyses were congruent. For mandarins, they confirm the previous observation of frequent introgressions of *C. maxima* genome in a *C. reticulata* background (Oueslati et al., 2017; Wu et al., 2018) and the probable role of *C. maxima* introgression in the domestication of mandarins (Wu et al., 2018). The small *C. reticulata* introgressions observed in Chr 2 of ‘Chandler’ and ‘Kao pan’ pummelos by Wu et al. (2018) are confirmed, suggesting a kinship between these two varieties. We also identified small C. *reticulata* introgressions in Chr 9 for ‘Guil’ and ‘Huan’ pummelos whereas a 0-5 Mb *C. medica* introgression was identified in the ‘Flores’ pummelo. Although previous studies (Ahmed et al., 2019; Wu et al., 2018) did not show interspecific introgression in *C. medica* accessions, here we revealed *C. micrantha* introgression in two citron varieties (‘Chinese’ and ‘Etrog e861’). The mosaic structure of the horticultural groups resulting from admixture as well as the assignment of cytoplasmic genomes corroborate previous results obtained using *C. clementina* V1.0 as a template genome (Wu et al., 2018) for sweet and sour oranges, grapefruits, Mediterranean lemons and Mexican limes. In addition, we confirmed the origin of the ‘Volkamer’ lemon and ‘Alemow’ lime from direct interspecific hybridizations between *C. reticulata* and *C. medica* on one side and *C. micrantha* and *C. medica* on the other side, as GBS data suggested (Ahmed et al., 2019). The two lemons and the two limes inherited, respectively, the *C. reticulata* and *C. micrantha* chloroplast, as previously proposed from cytoplasmic markers analysis (Curk et al., 2016). The contribution of *C. medica* as the male parent for these four interspecific hybrids aligned with the cleistogamy of this species. Introgression could have played an important role in citrus domestication. It would be interesting to conduct in-depth studies on the impact of ancestral species-specific variants (dSNPs, Indels, dPAV) involved in the introgressions in the cultivated citrus.

### Integration of plastid genome sequences in chromosome 4 of *C. medica* ancestor and its conservation in hybrid/admixed progenies

Our results revealed a complete chloroplast sequence insertion with several duplicated regions in the 17-18-Mb region of the Chr 4 of CITME genome assembly whereas no evidence of large chloroplastic sequence insertion was found in the genome assemblies of the three other ancestors of cultivated citrus. Moreover, this region’s chloroplastic genes were found in all *C. medica* accessions, in first-generation hybrids having *C. medica* as the male parent and in second-generation hybrids having inherited *C. medica* genome in the considered region, validating the nuclear location of these genes of chloroplastic origin. Part of these genes was also found in *C. indica*, an Indian species sharing a monophyletic origin with *C. medica*. Therefore, the considered nuclear insertion may have occurred after the first radiation of the Asian *Citrus* species (6-8 million years ago (Wu et al., 2018)) and before the secondary radiation between *C. indica* and *C. maxima* (supporting information, Figure 22b). Nuclear plastid DNA sequences (NUPTs) are frequently found in plant genomes, and it is accepted that the transfer of organellar DNA into the nuclear genome occurs continuously and contributes to genome structure and evolution (Leister and Kleine, 2011; Zhang et al., 2020). Plastid genes generally show very low or null expression in the nucleus because they often lack the regulatory element for gene expression in the nucleus or do not encode for completely open reading frames (Zhang et al., 2020; Zhao et al., 2019). Furthermore, epigenetic regulation has also been associated with the transcriptional repression of integrated chloroplastic genes (Yoshida et al., 2019). In the absence of functional implication, the evolution of NUPTs may be neutral and even selected against (Marczuk-Rojas et al., 2024). This is consistent with the fact that all plastid genes inserted in the Chr 4 of the CITME genome, except NDHJ, display large deletions compared with the original chloroplast gene sequences. Moreover, the expression of the integrated plastid genes appears insignificant based on our RNASeq data of the ‘Corsica’ citron (data not shown). The 17-18-Mb region of CITME Chr4 – in which the large plastid insertion is located – was included in a genomic area with very low gene sequence frequency and a high repeat-element level that probably corresponds to the centromeric region of Chr 4. Such insertion of plastid sequence in the centromere’s vicinity has been described in Arabidopsis and rice genomes (Michalovova et al., 2013). In a pioneer study of whole-chloroplast genome diversity based on WGS data (paired-end reads) of 34 citrus accessions mapped against the *C. sinensis* chloroplast genome (Bausher et al., 2006), Carbonell-Caballero et al. (2015) found frequent heteroplasmy in the citrus genus after interspecific hybridization. Their conclusions were based on the observation of SNPs with strong preeminent alleles and low frequency alleles, corresponding to those of the female and the male parents, respectively. The main gene pool sustaining their hypothesis was *C. medica* and F1 interspecific hybrids with *C. medica* as the male parent. According to our results, it is more likely that the observations of Carbonell-Caballero et al. (2015) for this gene pool resulted from the nuclear integration of the large *C. medica* plastid sequence in the *C. medica* nuclear genome. Our hypothesis is in agreement with the conclusion of Wang et al. (Wang et al., 2022a) for strict maternal inheritance of chloroplast genome in citrus.

### A super-pangenome for cultivated citrus

The implemented super-pangenome of cultivated citrus integrates the identification of orthologous genes of the four genome de novo assemblies and PAVs from resequencing data of 55 accessions. This analysis revealed 32,962 pangenes, 25,291 classified as core 2,431 as soft-Core, 5,171 as dispensable and 69 as private, reflecting citrus’s intricate genomic complexity and evolutionary history. The strong genetic structuration observed among the ancestral species and the alignment of horticultural groups with their phylogenomic backgrounds underscored this pangenome’s value in our understanding of the genetic diversity underlying modern cultivated citrus. 1,211 pangenes were found to be diagnostic for one of the four ancestral species, and specific absences (866) were more frequent than specific presences (345). Of the four ancestral taxa, *Citrus medica* displays the largest number of dPAV (612 pangenes). The comparison of the unique gene enrichments in each species with the core genome revealed distinct functional specializations that complement the conserved processes shared across the pangenome. Whereas the core genome is enriched on broad, essential functions, such as transcription regulation, stress response and protein turnover (Tranchant-Dubreuil et al., 2019), individual species display unique adaptations that enhance their survival and reproduction in specific environments. *Citrus reticulata*’s enrichment of auxin signalling and RNA repair likely provides refined growth control and stress resilience (Aslam et al., 2020; Jing et al., 2023) whereas *C. maxima*’s emphasis on autophagy and energy metabolism suggests robust mechanisms for maintaining cellular homeostasis (Magen et al., 2022). The absence of certain biosynthetic pathways in *C. medica* contrasts with the core genome, possibly reflecting a trade-off or specialization in other areas. Meanwhile, *C. micrantha* enrichment in antiviral response and RNA processing indicates a specialized defence mechanism (Yang and Li, 2018), aligning with the core genome’s general stress response but with a more targeted focus. These differences highlight the evolutionary paths that have shaped each species, contributing to the diversity in the citrus super-pangenome while maintaining a shared foundation of essential biological processes.

### Genome-hub for the cultivated Citrus super-pangenome

The Citrus Genome Hub (CGH, https://citrus-genome-hub.southgreen.fr) was developed to support post-genomics efforts by centralizing citrus genomic information. It offers a set of user user-friendly interfaces for the analysis and visualizations of genomic data, providing a comprehensive platform for researchers to explore and interpret citrus genomes efficiently.

Users can search for genes in two ways, by querying using various filters, such as name, putative function, chromosomal position and functional annotation (putative function, InterPro domain, gene ontology…) and by sequence similarity using Blast. In both methods, results are displayed in a dynamic table that summarizes the relevant information about the search. Each entry includes a link to the gene report and the genome browser (Jbrowse), providing easy access to detailed gene information and visualizations.

Comparative analysis at the genome scale is supported by Synvisio, an interactive multiscale synteny visualization tool. This feature allows users to explore syntenic relationships between genomes at various scales, facilitating a detailed understanding of genomic structural variation and conservation across species.

A download section provides users with direct access to the data used by the various tools that compose the hub.

## CONCLUSION

To implement a super-pangenome for cultivated *Citrus,* we performed *de novo* chromosome scale assemblies for *C. medica* (346 Mb), *C. micrantha* (333 Mb) and *C. reticulata* (332 Mb) for the first time. These assemblies were annotated for genes and repetitive elements, alongside a previously published assembly of *C. maxima*. We identified 28,090, 29,258, 29,477 and 30,101 genes in the respective genomes, with 1,627, 1,899, 2,014 and 1,851 of those genes categorized as PRRs and/or R genes. Despite their relatively small sizes, nearly half of each genome consists of transposable elements or DNA-satellites. Strong synteny and collinearity were observed among the three new genome assemblies whereas significant discrepancies were observed with the *C. maxima* one.

Intra- and interspecific diversity were explored by resequencing data from 55 citrus accessions. The phylogenomic structure along the genome of these accessions were revealed, highlighting interspecific introgressions in several representative accessions of *C. reticulata*, *C. maxima* and *C. medica* as well as the origin of horticultural groups shaped by reticulate evolution. A super-pangenome for cultivated *Citrus* was established by combining orthologous gene identification across the four assemblies and gene PAV based on resequencing data. This analysis revealed significant inter- and intraspecific variations, including 25,291 core genes, 2,431 soft-core genes and 5,171 dispensable genes. PAV analysis also revealed genes with presence or absence specific to particular ancestral taxa. In particular, these dPAVs revealed a large chloroplastic introgression on chromosome 4 of *C. medica*, which is inherited by horticultural groups and recent hybrids having this species as the male parent. Comparing the unique gene enrichments or those lost in each species with the core genome revealed distinct functional specializations that complement the conserved processes shared across the pangenome. Whereas the core genome is characterized by broad, essential functions, such as transcription regulation, stress response and protein turnover, individual species exhibit unique adaptations enhancing their survival and reproduction in specific environments.

We established a dedicated Genome Hub to provide a platform for ongoing genomic research, allowing for continuous updates and the future inclusion of additional ancestral species. This super-pangenome offers a comprehensive resource that captures the vast genetic diversity in modern cultivated citrus. Together with specialized tools, it will enhance the efficiency of association genetics and transcriptomics studies, expanding our understanding of the molecular basis of phenotypic diversity in cultivated Citrus. Ultimately, this resource will contribute to breeding programs focused on sustainable citriculture.

## EXPERIMENTAL PROCEDURES

### Genomic sequences of citrus accessions

#### Germplasm accession used for sequencing and optical mapping for de novo assembly of three accessions

The genome assemblies of *C. reticulata*, *C. medica* and *C. micrantha* were implemented respectively from the ‘Cleopatra’ mandarin (ICVN-0110274), ‘Corsican’ citron (SRA-613) and ‘Biasong’ (SRA-1115) localized on the Corsican field collection hosted by the Citrus Biological Resource Center (Citrus BCR) (INRAE-CIRAD, San Giuliano, Corsica, France).

#### Resequencing of citrus accessions

For variant calling on the four genome assemblies, a set of 40 citrus varieties was re-sequenced (Supporting Information Table 1). This set included 30 representatives of the four ancestral species and 10 representatives of admixed horticultural groups. Additionally, for the analysis of chloroplastic sequences, we re-sequenced 6 accessions representing 5 additional pure species, 3 admixed varieties and 6 controlled hybrids. All these accessions were also sources from the Corsican field collection hosted by the Citrus BRC (INRAE-CIRAD) and Cirad-INRAE breeding program in San Giuliano, Corsica, France. Total DNA was extracted using the Plant DNAeasy kit (Qiagen), following the manufacturer’s instructions. Genomic libraries were prepared according to standardized Illumina protocols for plant DNA and pair-end sequences (2×150) using an Illumina HiSeq 4000 platform at Genoscope, with an average coverage of 30X per accession. Demultiplexing, quality filtering and adapter removal were carried out according to the standardized protocol at Genoscope.

### *De novo* nuclear genome assembly

The de novo genome assemblies were obtained by combining nanopore long-read sequencing, Illumina PCR-Free sequencing and bionano optical mapping.

#### PromethION library preparation and sequencing

The libraries were prepared according to the ‘1D Genomic DNA by ligation (Kit 9 chemistry)’ protocol, provided by Oxford Nanopore. Genomic DNA was first repaired and end-prepped with the NEBNext FFPE Repair Mix (New England Biolabs, Ipswich, MA, USA) and the NEBNext® Ultra™ II End Repair/dA-Tailing Module (NEB). DNA was then purified with AMPure XP beads (Beckmann Coulter, Brea, CA, USA), and sequencing adapters provided by Oxford Nanopore Technologies were ligated using Concentrated T4 DNA Ligase 2M U/ml (NEB). After purification with AMPure XP beads (Beckman Coulter) using dilution buffer (ONT) and wash buffer (ONT), the library was mixed with the sequencing buffer (ONT) and the Library Loading Bead (ONT) and loaded on the PromethION R9.4.1 Flow Cells. The nanopore long reads were not cleaned, and raw reads were used for genome assembly.

#### Illumina PCR-Free library preparation and sequencing

PCR free libraries were prepared using the Kapa Hyper Prep Kit (Roche, Basel, Switzerland). Briefly, genomic DNA (1-1.5μg) was sonicated using a Covaris E220 sonicator (Covaris, Woburn, MA, USA). Fragments were end-repaired and 3ʹ-adenylated, and Illumina adapters (Bioo Scientific, Austin, TX, USA) were ligated following the manufacturer’s instructions. Ligation products were purified twice with AMPure XP beads (Beckman Coulter Genomics, Danvers, MA, USA) and quantified by qPCR (MxPro, Agilent Technologies, Santa Clara, CA, USA) using the KAPA Library Quantification Kit for Illumina Libraries (Roche). The library profile was assessed using an Agilent High Sensitivity DNA kit on the Agilent 2100 Bioanalyser. Libraries were paired-end sequenced on an Illumina HiSeq2500 instrument (Illumina, San Diego, CA, USA) using 250 base-length read chemistry. After the Illumina sequencing, an in-house quality control process was applied to the reads that passed the Illumina quality filters. The first step discards low-quality nucleotides (Q < 20) from both ends of the reads. Next, Illumina sequencing adapters and primer sequences were removed from the reads. Then, reads shorter than 30 nucleotides after trimming were discarded. These trimming and removal steps were completed using in-house-designed software based on the FastX package (Engelen and Aury; https://www.genoscope.cns.fr/fastxtend). The last step is to identify and discard read pairs that are mapped to the phage phiX genome, using SOAP aligner (Li et al., 2008), and the Enterobacteria phage PhiX174 reference sequence (GenBank: NC_001422.1). This processing, described in Alberti et al. (2020), resulted in high-quality data.

#### Optical mapping

The direct label and stain labelling (using the DLE-1 enzyme) and the nick label repair and stain (NLRS) labelling (using the BspQI enzyme) protocols and Bionano Genomics protocols were followed with 750ng and 600ng of DNA, respectively. The chip loadings were performed as Bionano Genomics recommended.

#### Long-read-based genome assembly

First, we estimated each organism’s genome size and heterozygosity rate using Genomescope2.0 (Ranallo-Benavidez et al., 2020) with the Illumina reads as inputs. Using these values, we launched the Necat assembler (Chen et al., 2021)(git commit 0c5fc85) with the appropriate genome size and with all Nanopore reads as inputs. Each assembly was then polished two times with HAPO-G (Aury and Istace, 2021) and Illumina reads.

#### Chromosome sequence reconstruction

Hybrid scaffolding: the BspQI and DLE-1 preparations were run on a single flow cell each. The generated molecules were assembled to produce optical maps using software provided by Bionano Genomics with the following two options: ‘add pre-assembly’ and non haplotype without extend and split’ (bionano solve and tools Version: 3.3_10252018). The two optical maps and the ONT contigs were subjected to the 2-enzyme hybrid scaffolding pipeline to generate the hybrid scaffolds. BisCoT (Istace et al., 2020) was used to correct artifactual duplications (negative gaps) introduced during the scaffolding process.

Pseudochromosome construction: for the ‘Cleopatra’ mandarin and ‘Corsican’ citron, the final genome assembly in pseudochromosomes was guided by anchoring their scaffolds to the genetic maps of the same varieties (Ollitrault et al., submitted), using the locOnRef tool from the Scaffhunter toolbox (Martin et al., 2016). For *C. micrantha,* for which no genetic map is available, the final assembly was guided by the synteny and collinearity with the *C. clementina* V1.0 pseudochromosome assembly (https://phytozome-next.jgi.doe.gov/info/Cclementina_v1_0). Chromosome numbering and orientation of the three *de novo* genomes are identical to those of *C. clementina* V1.0 (https://phytozome-next.jgi.doe.gov/info/Cclementina_v1_0) and *P. trifoliata* V1.3.1 (https://phytozome-next.jgi.doe.gov/info/Ptrifoliata_v1_3_1).

### Chloroplast genome assembly

Raw WGS-paired Illumina HiSeq 4000 sequences of ‘Corsican’ citron, *C. micrantha* and ‘Cleopatra’ mandarins were treated with fastQC v0.11.7, and *de novo* assembly of CITME, CITMI and CITRE chloroplastic genomes was performed with novoplasty v4.3.1 (Dierckxsens et al., 2017), using the default parameters and the Pineapple sweet orange chloroplastic genome NC_008334.1 as an assembly guide. The assembled circular sequence was annotated with GeSeq v2.03 (Dec 18, 2020) using default parameters. A phylogeny was realized with these three new assembled chloroplastic genomes and was completed with 23 already published assembled genomes of Rutaceae, 21 *Aurantioideae* species (Shi et al., 2023a, 2023b) and 2 outgroups species (*Zanthoxylum madagascariense* NC_046744 and *Zanthoxylum asiaticum* NC_056094). All sequences were aligned with MAFFT v7.045 (Katoh and Standley, 2013), and maximum likelihood analysis was conducted with phyML 3.0 (Guindon et al., 2010) after selection of the best evolutionary model by SMS (Lefort et al., 2017).

### Annotation of the three *de novo* genomes and the Chinese *C. maxima* assemblies

To compare our three *de novo* assemblies and the *C. maxima* (HWB.v1.0) available at http://citrus.hzau.edu.cn/download.php, we annotated the four genome assemblies following the procedure described below.

#### Annotation of genes

##### RNA-Seq experiment and transcriptome assembly

Three replicates of young leaves, mature leaves and flowers from the Cleopatra mandarin (ICVN-0110274), Corsican citron (SRA-613), Chandler pummelo (SRA-608) and *C. micrantha* (SRA-1115) were harvested on adult trees of the Corsican field collection hosted by the Citrus Biological Resource Center (Citrus BCR (INRAE-CIRAD, San Giuliano, Corsica, France). Three root replicates from the same genotypes were taken from cuttings in greenhouses. RNA was extracted using the NucleoMag® kit (Macherey-Nagel GmbH & Co.KU, Düren, Germany) with the Kingfisher automated system (KingFisher Flex Purification System, ThermoFisher Scientific, Waltham, MA) according to the manufacturer’s protocol. RNA integrity and quality were checked on TapeStation, screentape D5000. RNAseq libraries were prepared to obtain labelled and matched sequences using UDI indexes, following the protocol and recommendations from the Illumina Truseq kit. The libraries’ integrity and quality were checked on TapeStation, screentape D5000, and assayed by qPCR. An equimolar pool of the libraries was then built up using the TruSeq Stranded mRNA Sample Preparation kit from Illumina. Pair-end sequencing (2 x 150 nt) was performed on an Illumina HiSeq 4000 platform at Genewiz. Clean RNA-seq reads, which had been filtered by Trimmomatic (v0.39), were mapped onto their respective assemblies using Hisat2 (v2.1.0) with default parameters. Each of these files was then coordinate-sorted and converted to BAM format. The resulting BAM file was used for genome-guided transcriptome assembly using Trinity.

##### Prediction of protein-coding gene models

Automatic gene prediction was performed using the EuGene Eukaryotic Pipeline (EGNEP v.1.5 with EuGene v.4.2a), an integrative gene finder software that is able to combine several sources of information to predict genes (Sallet et al., 2019).

A EuGene internal prediction model can be built and trained using proteomic and transcriptomic data. Therefore, a set of protein coding gene models of *C. maxima* and *C. medica* proteins (Wang et al., 2017) as well as the UniProt/SwissProt Viriplantae database were used for the proteomic evidence, with the prior exclusion of proteins that were similar to those present in RepBase (v. 20.05). The CIRAD obtained an institutional license from https://phoenixbioinfo.org/repbase (ref:_00Do0J6b5._5001J10Do2W:ref; 07/23/2021-07/22/2024, access extended to 08/11/2024) to use the Repbase database (https://www.girinst.org/repbase). Two datasets of transcribed sequences were aligned on the genome and used by EuGene as transcription evidence: (i) the respective genome-guided transcriptome assembly produced by Trinity and (ii) a collection of more than 560,000 ESTs retrieved from the NCBI database using the query “Citrus”[Organism] AND plants[filter] AND biomol_mrna[PROP] AND is_est[filter]. Finally, *ab initio* was predicted using Augustus v3.3.2. To enhance prediction accuracy, a custom statistical model for splice-site detection was applied, leveraging information from both transcriptomic assemblies, using the egn_build_wam.pl companion script. Additionally, tRNAscan-SE was used to predict tRNA genes. Other non-coding RNAs, such as rRNA, miRNA and snRNA, were annotated using INFERNAL v1.1.2 through a search against the Rfam database v9.1

##### Functional annotation of protein-coding genes

To identify gene functions, we first used Diamond (v2.0.11) to annotate protein-coding genes’ functions based on items in the Swiss-Prot and TrEMBL databases (v2022-06-07; https://www.uniprot.org). Additionally, structural domains and GO annotations of predicted protein-coding genes were generated using InterProScan (v5. 48-83.0). KEGG orthology (KO) terms were assigned through searches against the hidden Markov model (KOfam) database using KofamScan v1.3.036 with default parameters.

##### Annotation of PRR

Domain annotation of the PRR proteins from the four genomes was performed using InterProScan 5.48-83.0 (Jones et al., 2014) with the default parameters. Subsequently, the RRGPredictor tool was employed to identify and classify the proteins into 13 classes (T, TN, TNL, RPW8NL, RLP, RLK, RLKGNK2, N, NL, CN, CNL, MLO and UNKNOWN) according to the methodology Santana Silva and Micheli (2020) described. The UNKNOWN class was used only to estimate the global PRR quantity and was excluded from other analyses. The genes on the chromosomes were graphically visualized using the TBtools tool (Chen et al., 2020). Gene duplication analysis for each PRR class and each genome was conducted using the MCScanX tool (Wang et al., 2012) with the default parameters. The search for PRR gene clusters was carried out by subdividing this analysis into two steps. First, clusters with three or more genes were identified following Holub’s (2001) methodology for genes included in a region smaller than or equal to 200 kb. Then, for the remaining genes, the clusters of two genes were identified following the methodology of Jupe et al. (2012).

#### Annotation of repeated sequences in genomes

The number of TEs was estimated using the *Citrus* repeat databank, developed by Giraud et al. (submitted). This bank was built from the genome assemblies and contained all the diversity of repeats found in the four *ancestral* species of cultivated citrus, including diversity of TEs, endogenous viral elements and DNA satellites. The TEannot pipeline from the REPET package was used for annotating the studied genomes (Flutre et al., 2011; Hoede et al., 2014; Quesneville et al., 2005). The pipeline aligns *Citrus* consensus sequences against the four reference genome sequences, using such tools as NCBI-Blast, RepeatMasker, CENSOR, BLASTER, SSR annotation with TRF, Mreps and RMSSR. Matches with a size larger than 100 bp and with at least 80% of homology with TEs (following the Wicker rule) were considered in the final annotation (Wicker et al., 2007). Additionally, intraspecific variability analyses were conducted to confirm that the TE amount annotated in assembled genomes was representative of the diversity of TEs founds in all the varieties of the four *Citrus* species ancestors. For this purpose, raw Illumina sequencing libraries from 10 varieties for each ancestor were aligned against the *Citrus* repeat databank using Bowtie2 (Langmead and Salzberg, 2012) (settings parameters: very sensitive local alignment, considered not-paired reads). The ten varieties selected by ancestor (only four varieties for *C. micrantha*) were chosen because the phylogenomic analysis showed they were pure representatives of the ancestors (supporting information, Table 1). The Illumina libraries used to assemble the reference genomes were included in the analyses. Finally, to enhance gene annotations, predicted genes showing sequence homology with TEs were removed. Genes were considered repeated sequences when more than 50% of their sequences were aligned with TEs (comparison of gene and repeat positions) and when they contained a coding protein domain associated with TEs.

#### Syntenic gene alignments

We estimated orthology and synteny between protein-coding genes for the 4 taxa using the R package GENESPACE v1.1.8 (Lovell et al., 2022). This package leverages MCScanX (Wang et al., 2012) to infer syntenic gene blocks and incorporates OrthoFinder v2.5.4 (Emms and Kelly, 2019) and DIAMOND v2.1.4.158 (Buchfink et al., 2021) to identify orthogroups in these syntenic blocks. This approach allowed us to analyse conserved gene relationships and structural genomic features across the taxa.

### Variant calling from resequencing data

The re-sequencing data from 40 new accessions generated for this study were combined with raw data from NCBI for 15 additional accessions (supporting information, Table 1). Of these, 11 were previously published by Wang et al. (2018, 2017) and 4 by Wu et al. (2018, 2014). A dedicated pipeline was developed using Snakemake to handle sequencing data (https://gitlab.cirad.fr/agap/workflows/GATK/-/tree/Pangenome_citrus?ref_type=tags). This pipeline automates various steps of data handling, ensuring reproducibility and scalability for large-scale genomic analyses. The raw data were cleaned using fastp (v 0.20.1). Variant calling was performed with GATK4 (https://github.com/gatk-workflows), using a standardized procedure and parameters, including the following steps: BWA alignment, SortSam, Mark Duplicate, HAploTypeCaller0, Select-VAriant, Variant-Filtration, Exclude-Filtered, BaseRecalibrator, ApplyBQSR, HAplotypeCAller1, GenomicsDB and GenotypeGVCFs. Four reference genome assemblies were used for variant calling: the three new ones released in this paper and the previously published HWB.v1.0 assembly for *C. maxima* (Wang et al., 2017), called CITMA in this work.

### Admixture analyses

The detailed method for interspecific mosaic analysis is provided in supporting information, Text 1 and Figure 13.

#### Identification of ancestral species’ diagnostic SNPs

We applied the two-step method Ahmed et al. (2019) proposed to identify diallelic dSNPs for each of the four ancestral species. Among the accessions classified as *C. reticulata,* we discarded clementine, ‘Satsuma’ and ‘King’ mandarins, found to be highly introgressed by *C. maxima*. The identification of dSNPs was therefore based on 11 citrons, 13 mandarins, 13 pummelos and 3 papedas (supporting information, Table 1). The first step of the dSNP search was intended to identify introgressions on the representative cultivars. It relied on the heterozygosity level and each accession’s similarity to the centroid of each ancestral taxon along the genome. SNPs with a differentiation parameter (GST) value >= 0.5 were selected to contrast areas with and without interspecific introgressions. Subsequently, the introgressed areas of each accession were removed and a second run was performed to select dSNPs with GST = 1 for the differentiation of one of the ancestral species from the three others.

#### Admixture along the four ancestral genomes

TraceAncestor software (Ahmed et al., 2019) was adapted for WGS data from diploid genotypes. Genotyping data were used instead of read counts to conduct maximum-likelihood tests of each ancestor contribution (i.e., allelic doses = 0, 0.5, 1 dose) in genomic windows of 20 dSNPs. Each ancestral taxon’s doses were then transferred to 100-kb consecutive windows all along the genome. In cases in which the sum of ancestral contributions exceeded 2, the corresponding windows were considered undetermined for the two haplotypes. The identification of dSNPs and mosaic structure analyses were performed using the SniPlay web tool (https://sniplay.southgreen.fr/cgi-bin/home.cgi). The DSNP databases for the four genome assemblies are available at CGH (https://citrus-genome-hub.southgreen.fr).

### Gene presence/absence variation from resequencing data

Gene PAV analysis was conducted by aligning DNA sequencing reads from each accession to the four reference genomes using Bowtie2 v2.4.2 in the ‘very-sensitive-local’ mode.

Gene PAVs were determined using SGSGeneLoss v0.1 (Golicz et al., 2015) according to the cumulative reads coverage of exons within each protein-coding gene. A gene was classified as ‘present’ if at least 90% of its exons were covered by a minimum of five reads (parameters: minCov = 5, lostCutoff = 0.9); otherwise, it was considered ‘absent’.

### Gene-based pangenome construction

#### Identification of orthologous genes among the four genome assemblies and pangene identification

To enable a comparison of CITME, CITMI, CITRE and CITMA genome assemblies, we used PanTools v3.4.0 (Jonkheer et al., 2022) to calculate homologous relationships in a predicted panproteome of these 4 species. Initially, we determined the optimal setting for relaxation using the optimal grouping. Based on the results, we selected a value of 2 for the relaxation parameter for homology grouping. Subsequently, gene classification of the homology groups was also done with PanTools.

#### Identification of core and dispensable genes among the pangenes from the PAV analysis of resequencing data from 55 cultivated citrus accessions

The pangenome was defined by combining the PAV information from resequencing data mapped on the four ancestor assemblies and the identification of the orthologous genes of the four genome assemblies (Figure 6). A pangene was considered present in a given accession if at least one of the corresponding orthologous gene identified in the four assemblies was found present for this accession. The number of shared groups was visualized using the tool available on the following page: https://bioinformatics.psb.ugent.be/cgi-bin/liste/Venn/calculate_venn.htpl. This approach allowed us to identify common homologous gene groups among the four *Citrus* species, providing insights into their evolutionary relationships and shared genetic features.

### GO term annotation and enrichment analysis

For each pangene, GO information from all source genes was merged to obtain nonredundant per pangene GO information. This was used to perform GO enrichment using a hypergeometric test with the clusterProfiler Bioconductor package (version 4.12.0) through the DIANE package (Version 1.1) [https://doi.org/10.1186/s12864-021-07659-2] with default parameters (qvalue = 0.1, p = 0.05).

The significantly enriched GO terms from the DIANE analysis were then summarized and visualized using REVIGO (http://revigo.irb.hr/). REVIGO was utilized to reduce the redundancy of GO terms by clustering semantically similar terms. The figures were generated with R and customized using ggplot2 (v3.5.1) and treemap (v2.4.4) when necessary.

## Acknowledgments

This study was conducted with financial support from CIRAD, the Genoscope, the *Commissariat à l’Énergie Atomique et aux Énergies Alternatives* and *France Génomique* (ANR-10-INBS-09–08). This work received support from MESO@LR-Platform at the University of Montpellier and technical support from the bioinformatics group of the UMR AGAP Institute, member of the French Institute of Bioinformatics (IFB) - South Green Bioinformatics Platform.

## Data availability

Raw data from long-read sequencing and genome assemblies of *C. micrantha*, *C. medica* and *C. reticulata* were deposited at EBI under the following accession numbers:

CITMI (*C. micrantha*): Raw data: PRJEB78372 - Assemblage: PRJEB78515 - Umbrella: PRJEB78526 CITME (*C. medica*): Raw data: PRJEB78445 - Assemblage: PRJEB78498 - Umbrella: PRJEB78525 CITRE (C. reticulata): Raw data: PRJEB78477 - Assemblage: PRJEB78537 - Umbrella: PRJEB78524

Resequencing raw reads derived from 58 citrus accessions were deposited in the EBI database under AC experiment accessions ERX12760151, ERX12780957, ERX12791201 and ERX12817346 to ERX12817400 (supporting information, Table 1).

The raw RNA-seq data for *C. medica*, *C. reticulata*, *C. maxima* and *C. micrantha* were deposited in the European Nucleotide Archive under accession code PRJEB77075.

The pseudochromosome and chloroplast genome assemblies of *C. medica, C. reticulata* and *C. micrantha* and their annotations are available on our website, https://citrus-genome-hub.southgreen.fr.

## Contributions

PO conceived and coordinated the experiments and analysis; MM,EM, SS, CC and KL performed DNA/RNA extractions; CC, KL and AL coordinated DNA sequencing; CB, SD, CC, KL, BI and JMA managed the sequencing data and produced the nuclear assemblies; GC produced the chloroplast assemblies, annotation and phylogenic studies; PM, MM and GC performed RNA sequencing and analysis; DG, NC, FM and SSB annotated repeated sequences; GD performed the gene annotation and structural variation studies; FDSM, RS and FM annotated PRR genes; FC, PO, AD and FL conducted the analysis of intra and interspecific diversity from resequencing data; GD, AS and PO conducted the gene PAV analysis; GD, AD, AS, BH and PO implemented the super-pangenome; AS and BH conducted the analysis of functional enrichment; PO, DG, GD, FDSM, FM, BH, GC, FC and AS wrote the manuscript and prepared the tables and figures; AL, PW and JMA coordinated the project at Genoscope; and PO and GD coordinated the project at Cirad.

