## Supplemental Information for "A super-pangenome for cultivated citrus reveals evolutive features during the allopatric phase of their reticulate evolution"

### Supporting Information

#### 1) Data accession

**Table 1:** List of plant material used for re-sequencing data studies

| Common Name | Horticultural group | Latin name | Pan_Genome Code | Pan-genome | CITME-Int | TE | Ref sequence | Publication |
|---|---|---|---|---|---|---|---|---|
| Buddha's hand | Citron | C. medica | cit_BdHD | 1 | | 1 | SRR6188453 | Wu_et_al_2018 |
| Chinese | Citron | C. medica | cit_chinD | 1 | 1 | | ERX12817362 | This publication |
| Corsican | Citron | C. medica | cit_cor1D | 1 | 1 | 1 | ERX12780957 | This publication |
| Diamante | Citron | C. medica | cit_diamD | 1 | 1 | 1 | ERX12817361 | This publication |
| Etrog-861 | Citron | C. medica | cit_e861D | 1 | 1 | | ERX12817360 | This publication |
| Humpang | Citron | C. medica | cit_humpD | 1 | 1 | 1 | ERX12817388 | This publication |
| Mac Nao San | Citron | C. medica | cit_macnD | 1 | 1 | 1 | ERX12817363 | This publication |
| Mac Veu Mountain | Citron | C. medica | cit_macvD | 1 | | 1 | SRR6188459 | Wu_et_al_2018 |
| wild 1 | Citron | C. medica | cit_wc1D | 1 | 1 | 1 | SRR3938056 | Wang_et_al_2017 |
| wild 2 | Citron | C. medica | cit_wc2D | 1 | 1 | 1 | SRR3944139 | Wang_et_al_2017 |
| wild 3 | Citron | C. medica | cit_wc3D | 1 | 1 | 1 | SRR3944160 | Wang_et_al_2017 |
| Nules | Clementine | C. reticulata x C. x sinensis | cle_nuleD | 1 | 1 | | SRR1022654 | Wu_et_al_2014 |
| Duncan | Grapefruit | C. x paradisi | grf_duncD | 1 | | | ERX12817386 | This publication |
| Star ruby | Grapefruit | C. x paradisi | grf_starD | 1 | | | ERX12817383 | This publication |
| Eureka | Lemon | C. x limon | lem_eureD | 1 | 1 | | ERX12817382 | This publication |
| Volkamer | Lemon | C. x limonia | lem_volkD | 1 | 1 | | ERX12817390 | This publication |
| Alemow | Lime | C. x macrophylla | lim_alemD | 1 | 1 | | ERX12817373 | This publication |
| Mexican | Lime | C. x aurantiifolia | lim_mexiD | 1 | 1 | | ERX12817381 | This publication |
| Rangpur | Lime | C. limonia | lim_RangD | 1 | 1 | | SRR6188467 | Wu_et_al_2018 |
| Avana Tard. di Ciaculli | Mandarin | C. reticulata | man_avanD | 1 | | | ERX12817385 | This publication |
| Cleopatra | Mandarin | C. reticulata | man_clo1D | 1 | 1 | 1 | ERX12791201 | This publication |
| Daoxian wild No1 | Mandarin | C. reticulata | man_DaoW1D | 1 | | | SRR5796819 | Wang_et_al_2018 |
| Citrus daoxianensis | Mandarin | C. reticulata | man_daoxD | 1 | | | ERX12817369 | This publication |
| Hezhou wild | Mandarin | C. reticulata | man_HezWD | 1 | | 1 | SRR3747527 | Wang_et_al_2018 |
| Jiangyong wild | Mandarin | C. reticulata | man_JiaWD | 1 | | 1 | SRR5796862 | Wang_et_al_2018 |
| King of Siam | Mandarin | C. reticulata | man_kingD | 1 | | | ERX12817370 | This publication |
| MS2 | Mandarin | C. reticulata | man_ManWD | 1 | 1 | 1 | SRR5796635 | Wang_et_al_2018 |
| Nan feng mi chu | Mandarin | C. reticulata | man_nfmcD | 1 | | | ERX12817353 | This publication |
| Owari | Mandarin | C. reticulata | man_owarD | 1 | | | ERX12817346 | This publication |
| Ponkan | Mandarin | C. reticulata | man_ponkD | 1 | | | ERX12817347 | This publication |
| Sun chu sha | Mandarin | C. reticulata | man_scshD | 1 | | 1 | ERX12817352 | This publication |
| Shekwasha | Mandarin | C. reticulata | man_shekD | 1 | | | ERX12817371 | This publication |
| Sunki | Mandarin | C. reticulata | man_sunkD | 1 | | 1 | ERX12817350 | This publication |
| Szinkom | Mandarin | C. reticulata | man_szinD | 1 | | | ERX12817348 | This publication |
| Tachibana | Mandarin | C. reticulata | man_tachD | 1 | | 1 | ERX12817351 | This publication |
| Citrus macroptera | Papeda | C. macroptera | pap_macrD | 1 | 1 | 1 | ERX12817393 | This publication |
| Melanesian | Papeda | C. macroptera | pap_melaD | 1 | | 1 | ERX12817365 | This publication |
| Biasong | Papeda | C. micrantha | pap_mic1D | 1 | 1 | 1 | ERX12760151 | This publication |
| Chandler | Pummelo | C. maxima | pum_cha2D | 1 | 1 | | ERX12817358 | This publication |
| Haploid Pummelo | Pummelo | C. maxima | pum_chaH | 1 | | | ERX12817392 | This publication |
| Chongqing No016 | Pummelo | C. maxima | pum_ChonD | 1 | | 1 | SRR3822303 | Wang_et_al_2017 |
| Flores | Pummelo | C. maxima | pum_florD | 1 | | 1 | ERX12817349 | This publication |
| Guilin No1 | Pummelo | C. maxima | pum_GuilD | 1 | | | SRR3822291 | Wang_et_al_2017 |
| F22 | Pummelo | C. maxima | pum_hf22D | 1 | | 1 | ERX12817359 | This publication |
| Huanonghongyou | Pummelo | C. maxima | pum_HuanD | 1 | | | SRR3823230 | Wang_et_al_2017 |
| Huazhoujuhong | Pummelo | C. maxima | Pum_HuazD | 1 | | 1 | SRR3823225 | Wang_et_al_2017 |
| Indian | Pummelo | C. maxima | pum_indeD | 1 | 1 | 1 | ERX12817354 | This publication |
| Kao pan | Pummelo | C. maxima | pum_kaopD | 1 | | | ERX12817384 | This publication |
| Reinking | Pummelo | C. maxima | pum_reinD | 1 | | 1 | ERX12817356 | This publication |
| Tahiti | Pummelo | C. maxima | pum_tahiD | 1 | | 1 | ERX12817357 | This publication |
| Timor | Pummelo | C. maxima | pum_timoD | 1 | | | ERX12817355 | This publication |
| Bouquetier de Nice | Sour orange | C. x aurantium | soo_boniD | 1 | | | ERX12817366 | This publication |
| Maroc | Sour orange | C. x aurantium | soo_marD | 1 | | | ERX12817391 | This publication |
| Moro | Sweet orange | C. x sinensis | swo_moroD | 1 | | | ERX12817372 | This publication |
| Pineaple | Sweet orange | C. x sinensis | swo_pineD | 1 | | | ERX12817380 | This publication |
| Aus. desert lime | Aus. desert lime | C. glauca | adl_corsD | | 1 | | ERX12817379 | This publication |
| Aus. finger lime | Aus. finger lime | C. australasica | afl_cav1D | | 1 | | ERX12817389 | This publication |
| Aus. round lime | Aus. round lime | C. australis | arl_taloD | | 1 | | SRR6188449 | Wu_et_al_2018 |
| Bergamota | Bergamota | C. x bergamia | ber_castD | | 1 | | ERX12817374 | This publication |
| Nules x Alemow | Hybrid | C. reticulata x C. x aurantiifolia | hCl_alemD | | 1 | | ERX12817397 | This publication |
| Nules x Nestour | Hybrid | C. reticulata x C. x aurantiifolia | hCl_nestD | | 1 | | ERX12817400 | This publication |
| Nules x Rangpur | Hybrid | C. reticulata x C. x limonia | hCl_rangD | | 1 | | ERX12817395 | This publication |
| Nules x Volkamer | Hybrid | C. reticulata x C. x limonia | hCl_volkD | | 1 | | ERX12817399 | This publication |
| Chandler x Corsican | Hybrid | C. maxima x C. medica | hPc_corsD | | 1 | | ERX12817375 | This publication |
| Chandler x Eureka | Hybrid | C. maxima x C. x limon | hPc_eu92D | | 1 | | ERX12817394 | This publication |
| Citrus indica | Indica | C. indica | ind_wangD | | 1 | | SRR3948493 | Wang_et_al_2017 |
| Hong Kong | Kumquat | C. hindsii | kum_hindD | | 1 | | ERX12817396 | This publication |
| Marumi | Kumquat | C. japonica | kum_maruD | | 1 | | ERX12817387 | This publication |
| Rough | Lemon | C. x limonia | lem_roleD | | 1 | | ERX12817367 | This publication |
| Nestour | Lime | C. x aurantiifolia | lim_nestD | | 1 | | ERX12817368 | This publication |
| Eustis quat | Limequat | C. japonica x C. x. aurantiifolia | liq_eustD | | 1 | | ERX12817378 | This publication |
| Ichang | Papeda | C. ichangensis | pap_ichaD | | 1 | | ERX12817364 | This publication |
| Wild Ichang | Papeda | C. ichangensis | pap_wic1D | | 1 | | SRR3928244 | Wang_et_al_2017 |
| Pomeroy | Trifoliate orange | C. trifoliata | pon_pomeD | | 1 | | ERX12817376 | This publication |

| | | | | | | |
|---|---|---|---|---|---|---|
| Rubidoux | Trifoliate orange | C. trifoliata | pon_rubiD | 1 | ERX12817398 | This publication |
| Box orange | Box orange | Atalantia buxifolia | sev_buxiD | 1 | ERX12817377 | This publication |

Pan-genome: accessions used (1) for the pangenome implementation; CITME-Int: accessions used (1) for the study of chloroplast introgression in Chr4 of CITME; TE: accessions used (1) for transposable element intra-specific diversity study. Reference: (Wang et al., 2017, 2018; Wu et al., 2018)

#### 2) Nuclear genome assemblies

**Table 2:** Metrics of Oxford Nanopore long reads

| Species | *C. micrantha* | *C. medica* | *C. reticulata* |
|---|---|---|---|
| Assembly | CITMI | CITME | CITRE |
| Number of reads | 4,905,706 | 2,465,218 | 5,509,665 |
| Cumulative size | 42,738,216,849 | 32,344,708,635 | 52,334,600,298 |
| N50 | 18,003 | 26,334 | 19,676 |
| Coverage | 136x | 77x | 152x |

**Table 3:** Metrics of optical Bionano mapping data (molecules > 150Kb)

| Species | Assembly | | Cumulative size | nb of genome maps | N50 (Mb) |
|---|---|---|---|---|---|
| *C. micrantha* | CITMI | DLE-1 | 326.91 Gb | | 234.72 |
| | | DLE-1 optical map | 541.87 Mb | 79 | 12.03 |
| | | (after haplotype filtering) | (308.50 Mb) | (24) | 20.89 |
| | | BspQI | 302.55 Gb | | 250.81 |
| | | BspQI optical map | 562.85 Mb | 250 | 3.38 |
| | | (after haplotype filtering) | (386.72 Mb) | (171) | 3.66 |
| *C. medica* | CITME | DLE-1 | 282.57 Gb | | 266.74 |
| | | DLE-1 optical map | 385.36 Mb | 33 | 35.83 |
| | | BspQI | 425.78 Gb | | 265.01 |
| | | BspQI optical map | 410.95 Mb | 169 | 3.802 |
| *C. reticulata* | CITRE | DLE-1 | 284.26 Gb | | 255.25 |
| | | DLE-1 optical map | 482.62 Mb | 81 | 26.98 |
| | | (after haplotype filtering) | (263.89 Gb) | (13) | 30.2 |
| | | BspQI | 169.36 Gb | | 263.11 |
| | | BspQI optical map | 462.8 Mb | 226 | 2.67 |
| | | (after haplotype filtering) | (330.65 Mb) | (151) | 3.37 |

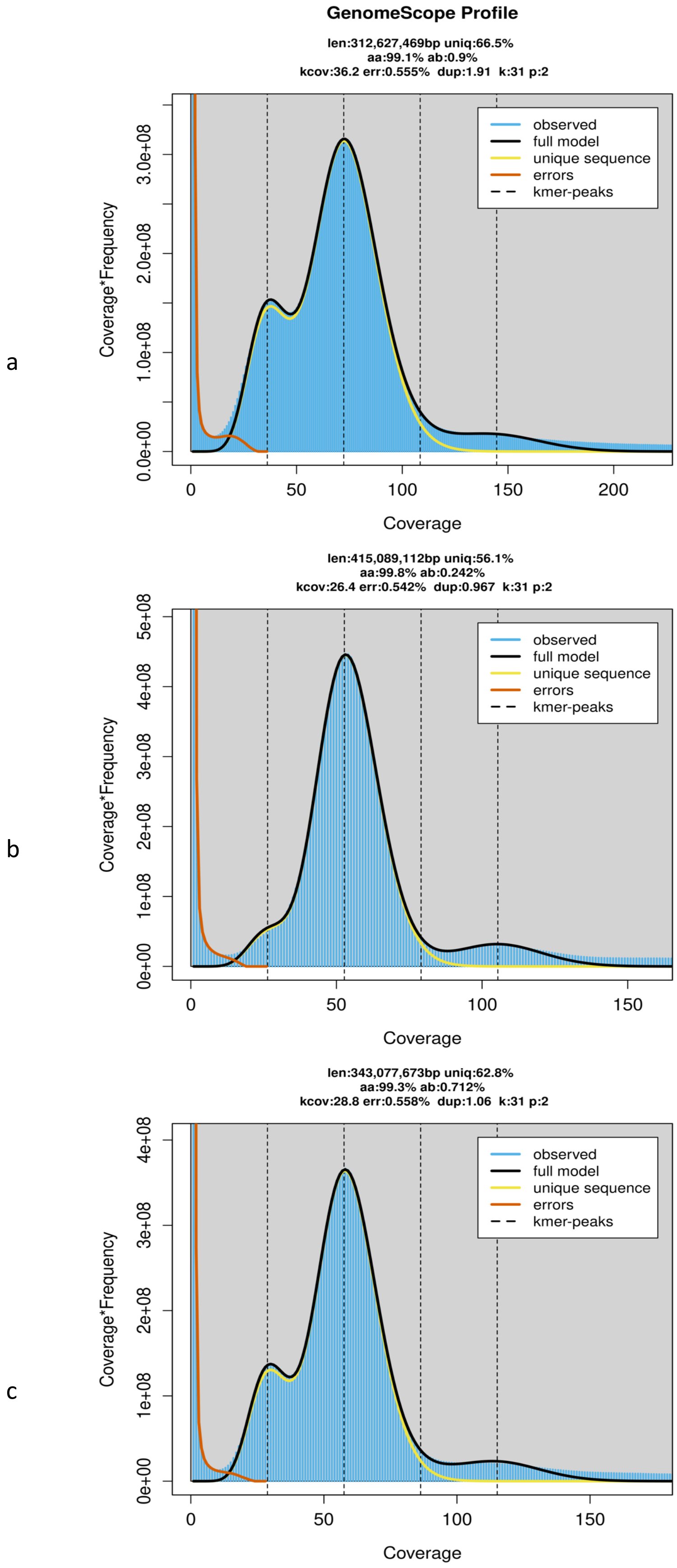

**Figure 1**: Genome size and heterozygosity rates estimations.
a: *C. micrantha*, b: *C. medica*, c: *C. reticulata*.

**Table 4:** Statistic of the scaffold assembly combining Nanopore long-range sequencing, Illumina PCR Free sequencing and Optical mapping

| Species | | *C. micrantha* | *C. medica* | *C. reticulata* |
|---|---|---|---|---|
| Assembly | | CITMI | CITME | CITRE |
| Number of scaffolds | | 314 | 141 | 310 |
| N50 scaffolds | | 23,232,283 | 33,362,141 | 29,540,655 |
| N90 scaffolds | | 2,809,014 | 28,156,396 | 1,429,702 |
| Min. scaffold size | | 10,640 | 11,623 | 10,290 |
| Max. scaffold size | | 37,639,730 | 51,661,475 | 50,999,016 |
| Cumul. scaffolds size | | 332,897,117 | 346,387,558 | 332,147,486 |
| Number of Ns (%) | | 12,108,336 (3.64%) | 3,711,038 (1.07%) | 3,960,896 (1.19%) |
| Number of contigs | | 447 | 199 | 448 |
| N50 contigs | | 4,200,014 | 10,930,299 | 5,154,950 |
| N90 contigs | | 393,380 | 2,826,067 | 367,416 |
| Busco (N=1614) | Complete | 1589 (98.5%) | 1594 (98.8%) | 1587 (98.4%) |
| | Dup. | 40 | 21 | 46 |
| | Frag. | 18 | 12 | 17 |
| | Missing | 7 | 8 | 10 |
| Merqury score | | 28.3194 | 29.1606 | 29.4018 |

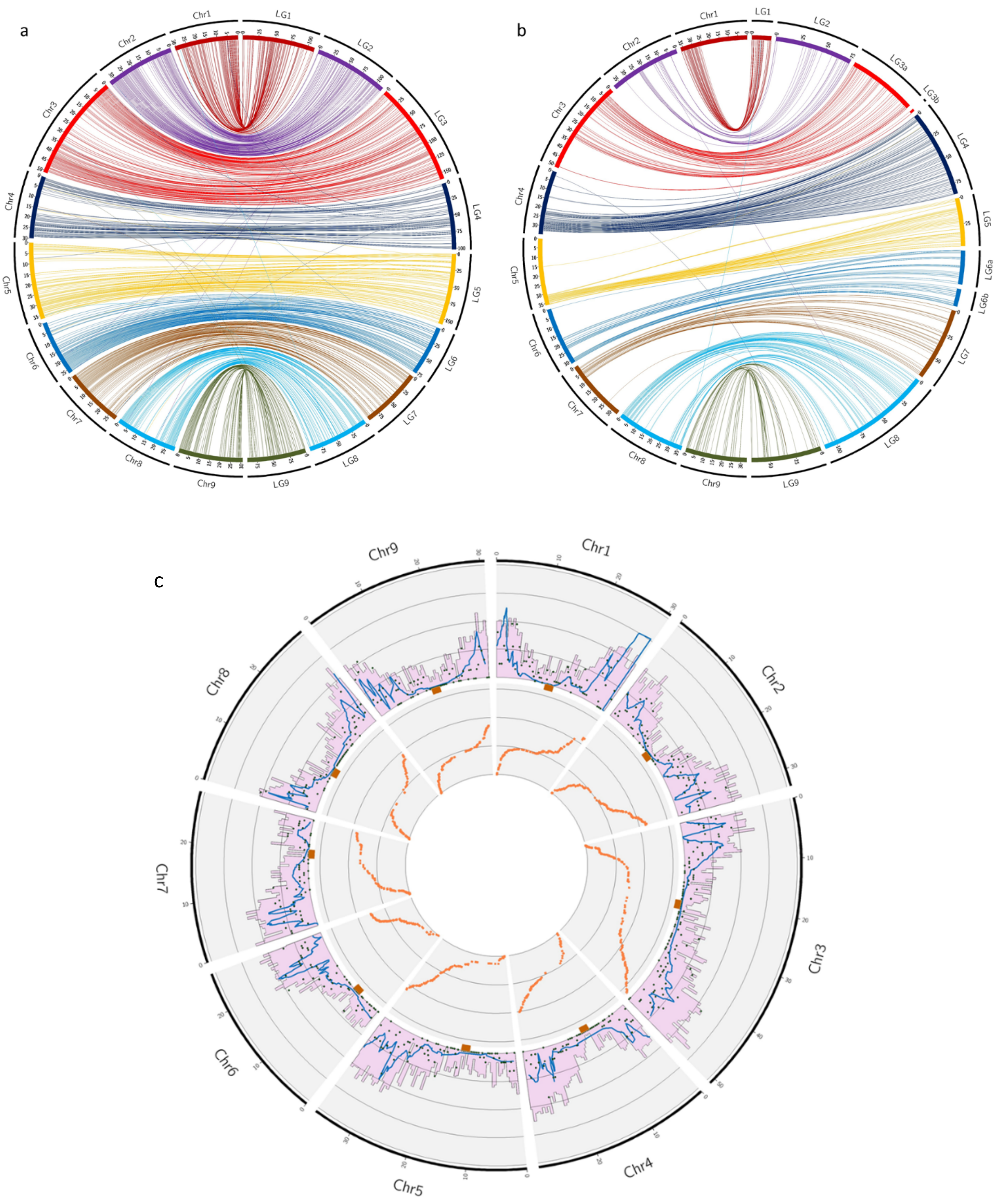

**Figure 2:** Anchorage of genetic maps on genome assemblies. a, b: Link between physical assemblies in pseudochromosomes (Chr) and genetic maps (LG)

a: Cleopatra mandarin; b: Corsican citron. c: Gene sequence density, recombination landscape and estimation of centromere location in CITRE genome. External Outer ring: Pink histogram: gene density (scale 0-100%), blue line local recombination (scale 0-30 cM/Mb; black dot: number of markers (scale: 0-5/100kb); central ring: estimated location of centromere; inner ring: Marey map (x: axis physical position; y axis genetical position (scale 0-160 cM).

**Table 5:** CITRE final assembly in pseudochomosomes and relation with Cleopatra mandarin genetic map

| Chr | Size (bp) | Nb scaffolds | Synt. G.M. | Col. G.M. |
|---|---|---|---|---|
| CITRE_001 | 30415407 | 1 | 98.58% | 0.999 |
| CITRE_002 | 32931837 | 1 | 99.55% | 0.999 |
| CITRE_003 | 50999016 | 1 | 98.10% | 0.968 |
| CITRE_004 | 29834578 | 2 | 96.40% | 0.934 |
| CITRE_005 | 36671209 | 5 | 98.64% | 0.923 |
| CITRE_006 | 26655275 | 1 | 98.19% | 0.928 |
| CITRE_007 | 28520055 | 1 | 98.68% | 0.951 |
| CITRE_008 | 29540655 | 1 | 97.10% | 0.953 |
| CITRE_009 | 30765278 | 1 | 100.00% | 0.956 |
| Total Chr | 296333310 | 14 | 98.46% | 0.957 |
| ChrUn | 35814676 | 296 | | |
| Total assembly | 332147986 | 310 | | |

Synt. G.M: synteny of the CITRE assembly with the Cleopatra mandarin genetic map
Col. G.M.: collinearity of the CITRE assembly with the Cleopatra mandarin genetic map

**Table 6:** CITME final assembly in pseudochomosomes and relation with Corsican citron genetic map

| Chr | Size (bp) | Nb scaffolds | Synt. G.M. | Col. G.M. |
|---|---|---|---|---|
| CITME_001 | 36435954 | 1 | 100.00% | 0.996 |
| CITME_002 | 38834663 | 2 | 97.22% | 0.987 |
| CITME_003 | 51661475 | 1 | 99.12% | 0.999 |
| CITME_004 | 34893391 | 2 | 99.25% | 0.861 |
| CITME_005 | 36212829 | 1 | 98.89% | 0.990 |
| CITME_006 | 31903664 | 2 | 100.00% | 0.827 |
| CITME_007 | 33922902 | 2 | 98.08% | 0.977 |
| CITME_008 | 37166674 | 1 | 99.09% | 0.902 |
| CITME_009 | 33362141 | 1 | 100.00% | 0.986 |
| Total Chr | 334393693 | 13 | 99.19% | 0.935 |
| ChrUn | 11994265 | 128 | | |
| Total assembly | 346387958 | 141 | | |

Synt. G.M: synteny of the CITME assembly with the Corsican citron genetic map
Col. G.M.: collinearity of the CITME assembly with the Corsican citron genetic map

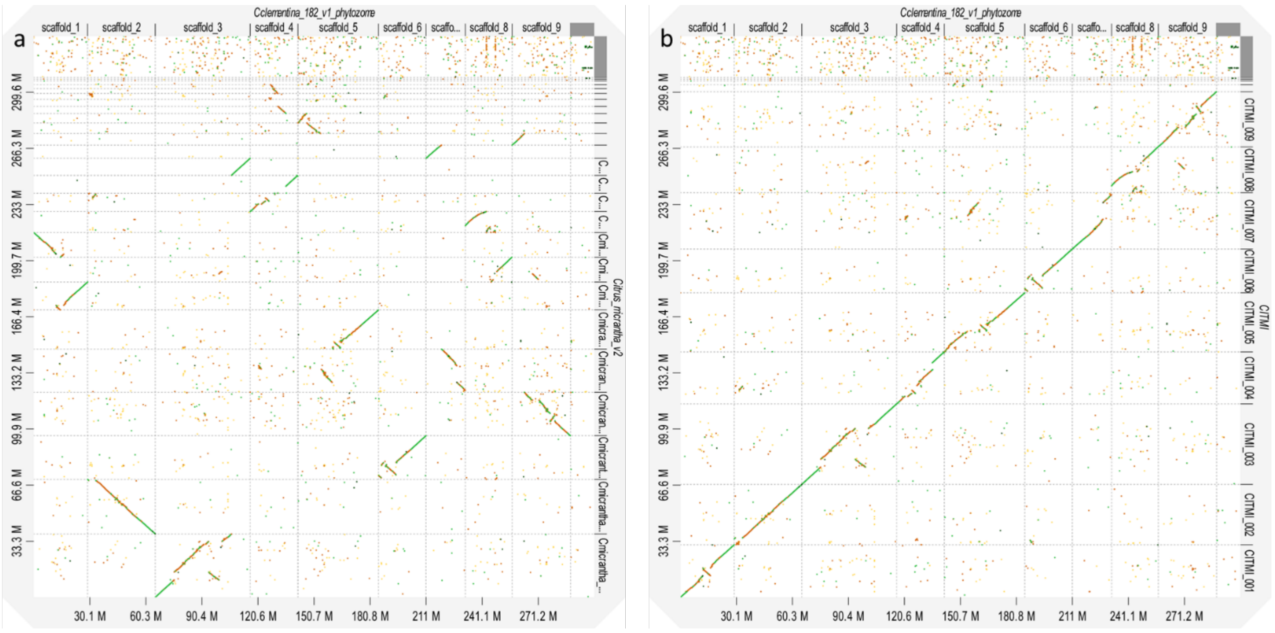

**Figure 3:** Dot plot of CITMI assemblies on the *C. clementina* V1.0 genome

a: CITMI scaffold assembly; b: CITMI pseudochromosome assembly

**Table 7:** CITMI final assembly in pseudochromosomes

| Chr | Size | Nb scaffolds |
|---|---|---|
| CITMI_001 | 31197090 | 2 |
| CITMI_002 | 35784675 | 2 |
| CITMI_003 | 47784602 | 2 |
| CITMI_004 | 30809907 | 5 |
| CITMI_005 | 35102979 | 3 |
| CITMI_006 | 25961129 | 1 |
| CITMI_007 | 33415567 | 2 |
| CITMI_008 | 27176488 | 2 |
| CITMI_009 | 32675320 | 2 |
| Total Chr | 299907757 | 21 |
| ChrUn | 32990560 | 293 |
| Total assembly | 332898317 | 314 |

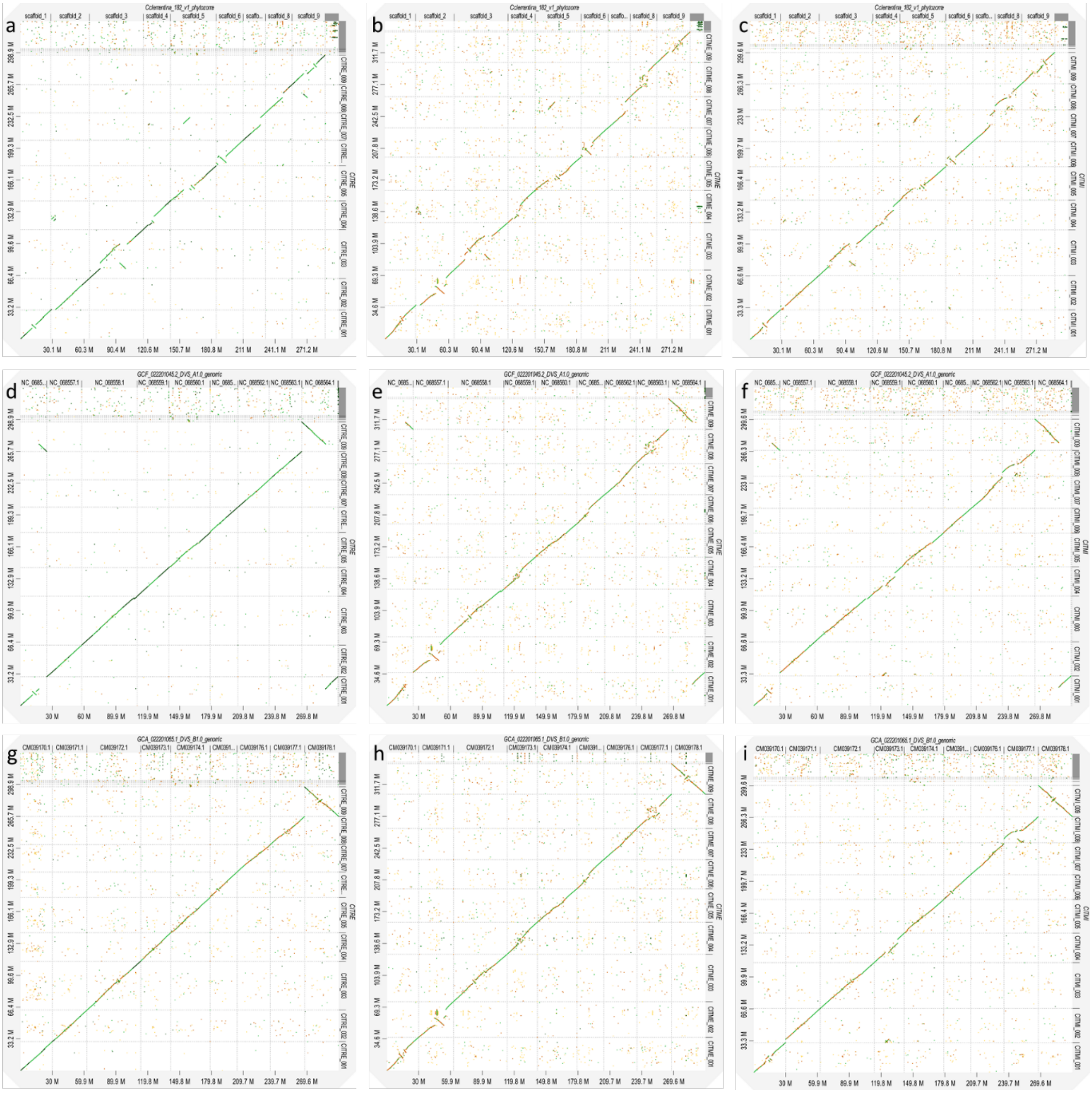

**Figure 4a**: Dot plots of the new assemblies with published genomes

CITRE (a,d,g), CITME (b,e,h) and CITMI (c,f,i) assemblies on *C. clementina* V1.0 genome (a,b,c), Sweet orange mandarin haplotype Citrus sinensis cv. Valencia DVS_A genome v1.0 (d,e,f) and Sweet orange pummelo Citrus sinensis cv. Valencia DVS_B genome v1.0haplotype (g,h,i) target genomes. The considered clementine and sweet orange genomes were published respectively by (Wu et al., 2014) and (Wu et al., 2022).

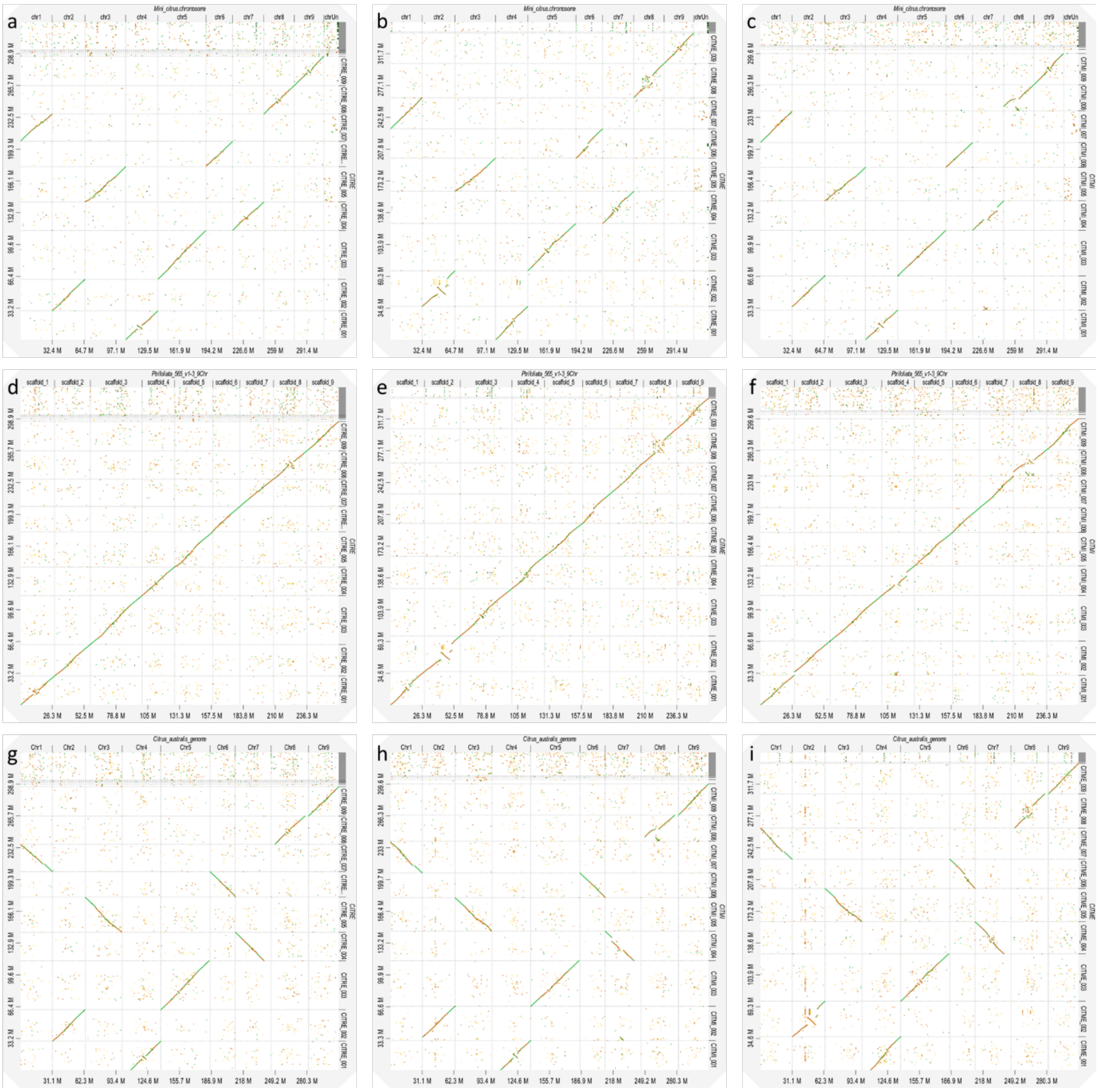

**Figure 4b:** Dot plots of the new assemblies with published genomes

CITRE (a,d,g), CITME (b,e,h) and CITMI (c,f,i) assemblies on *C. hindsii* (a,b,c), *C. trifoliata* (d,e,f) and *C. australis* (g,h,i) target genomes. The C. hindsii, C. trifoliata and C. australis genomes were published respectively by (Wang et al., 2022), (Peng et al., 2020), (Nakandala et al., 2023).

#### 3) Chloroplast assembly

**Table 8**: Characteristics of the three chloroplast *de novo* assemblies

| | CITMI | CITME | CITRE |
|---|---|---|---|
| Genome size | 159936 | 160016 | 160601 |
| %GC | 38 | 38 | 38 |
| LSC | 87191 | 87467 | 87801 |
| IR | 26991 | 29991 | 27008 |
| SSC | 18763 | 18567 | 18784 |
| Total nb genes | 95 | 91 | 97 |
| rRNAs | 10 | 10 | 10 |
| tRNAS | 37 | 37 | 37 |

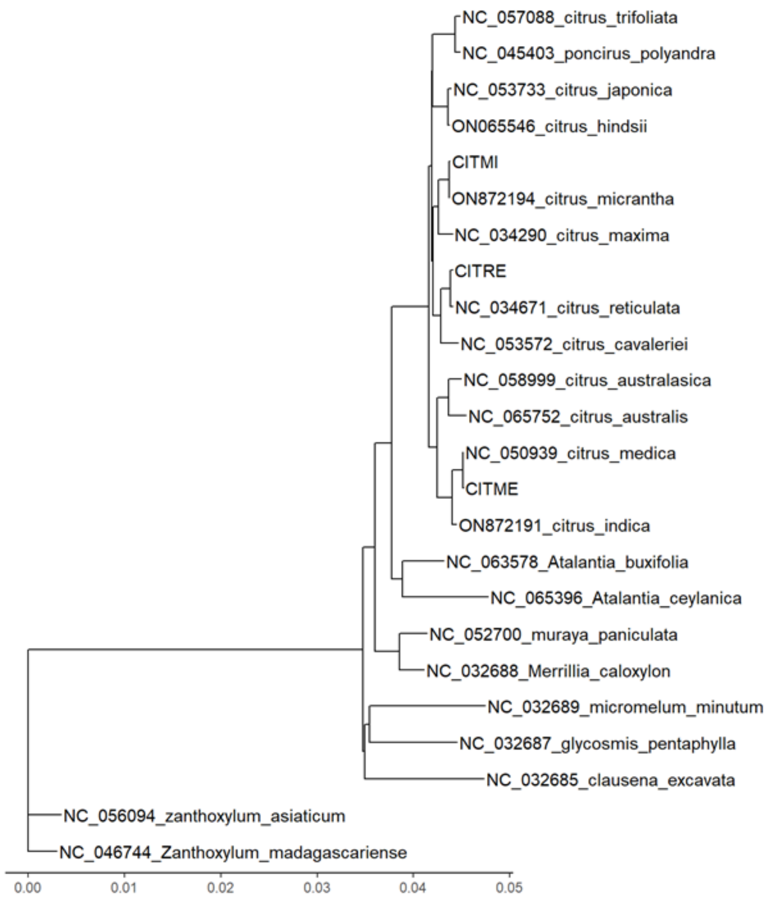

**Figure 5:** Phylogenic situation of the 3 chloroplast genome assemblies
Maximum likelihood analysis performed with phyML 3.0 (Guindon et al., 2010)

#### 4) Gene annotation

**Table 9:** Statistics for RNA-Seq
coverage are relative to the transcriptome size calculated for the corresponding ancestor genome assembly

| | | Number of Reads | Yield (Mbases) | Mean Quality score | percent Base_30 | Coverage | Alignment rate (%) | Number of transcripts |
|---|---|---|---|---|---|---|---|---|
| Micrantha | Mature_leaves | 59996009 | 17999 | 38.76 | 93.39 | 518.8 | 91.7 | 36,802 |
| | Young_leaves | 88062134 | 26418 | 39.04 | 94.43 | 761.5 | 94.24 | 36,536 |
| | Flowers | 42512999 | 12754 | 39.1 | 94.625 | 367.6 | 94.21 | 34,801 |
| | Roots | 67143229 | 20143 | 38.98 | 94.21 | 580.6 | 91.65 | 35,587 |
| Corsican Citron | Mature_leaves | 40547915 | 12165 | 38.85 | 93.71 | 370.4 | 94.65 | 33,053 |
| | Young_leaves | 73183886 | 21956 | 39.01 | 94.31 | 668.5 | 94.91 | 33,279 |
| | Flowers | 66708254 | 20012 | 39.02 | 94.37 | 609.3 | 93.83 | 36,648 |
| | Roots | 62756791 | 18827 | 38.98 | 94.2 | 573.3 | 92.49 | 33,229 |
| Cleopatra Mandarin | Mature_leaves | 58203297 | 17461 | 38.97 | 94.13 | 500.2 | 92.94 | 35,330 |
| | Young_leaves | 78467866 | 23540 | 39.03 | 94.37 | 674.4 | 94.28 | 31,921 |
| | Flowers | 79946794 | 23984 | 39.01 | 94.3 | 687.1 | 94.25 | 31,611 |
| | Roots | 50931681 | 15280 | 38.98 | 94.19 | 437.7 | 89.83 | 36.757 |
| Chandler Pummelo | Mature_leaves | 38609053 | 11583 | 39.16 | 94.79 | 336.6 | 92.44 | 33,570 |
| | Young_leaves | 71257033 | 21377 | 39 | 94.26 | 621.2 | 93.43 | 40,028 |
| | Flowers | 92066979 | 27620 | 39.05 | 94.45 | 802.6 | 94.89 | 38,157 |
| | Roots | 54709016 | 16413 | 39.03 | 94.39 | 476.9 | 92.83 | 36,267 |

**Table 10:** Number of genes and percentage per database with at least one associated term
SwissProt and TrEMBL for putative function inference; KEGG for metabolic pathways inference; InterPro for protein signature provided by several different databases.

| | CITMI | | CITME | | CITRE | | CITMA | |
|---|---|---|---|---|---|---|---|---|
| **Database** | **Nb** | **%** | **Nb** | **%** | **Nb** | **%** | **Nb** | **%** |
| SwissProt | 10983 | 37.54 | 10690 | 38.06 | 10517 | 35.68 | 11555 | 38.39 |
| TrEMBL | 18273 | 62.45 | 17400 | 61.94 | 17960 | 60.93 | 18546 | 61.61 |
| KEGG | 10281 | 35.14 | 10010 | 35.64 | 10407 | 35.31 | 11238 | 37.33 |
| InterPro | 24246 | 82.87 | 25541 | 90.93 | 26696 | 90.57 | 27600 | 91.69 |
| Pfam | 16016 | 54.74 | 15578 | 55.46 | 16187 | 54.91 | 17128 | 56.90 |
| SUPERFAMILY | 12422 | 42.46 | 12038 | 42.86 | 12578 | 42.67 | 13111 | 43.56 |
| Gene3D | 12367 | 42.27 | 15452 | 55.01 | 16290 | 55.26 | 17196 | 57.13 |
| PIRSF | 799 | 2.73 | 787 | 2.80 | 780 | 2.65 | 816 | 2.71 |
| SMART | 3830 | 13.09 | 3647 | 12.98 | 3902 | 13.24 | 3903 | 12.97 |
| ProSiteProfiles | 5853 | 20.00 | 5575 | 19.85 | 5990 | 20.32 | 5948 | 19.76 |
| SFLD | 146 | 0.50 | 140 | 0.50 | 151 | 0.51 | 174 | 0.58 |
| PANTHER | 4979 | 17.02 | 4853 | 17.28 | 5074 | 17.21 | 5116 | 17.00 |

| | | | | | | | | |
|---|---|---|---|---|---|---|---|---|
| PRINTS | 1835 | 6.27 | 1893 | 6.74 | 1859 | 6.31 | 2085 | 6.93 |
| CDD | 1997 | 6.83 | 2009 | 7.15 | 2082 | 7.06 | 2143 | 7.12 |
| ProSitePatterns | 4638 | 15.85 | 4541 | 16.17 | 4633 | 15.72 | 5161 | 17.15 |
| TIGRFAM | 1385 | 4.73 | 1367 | 4.87 | 1427 | 4.84 | 1841 | 6.12 |
| GO | 17425 | 59.56 | 22836 | 81.30 | 23585 | 80.01 | 24824 | 82.47 |
| **Total** | **29258** | **100%** | **28090** | **100%** | **29477** | **100%** | **30101** | **100%** |

**Table 11:** Numbers of annotated genes by chromosome of the three *de novo* assembly and *C. maxima* assembly and gene completeness

| Chr | CITMI | CITME | CITRE | CITMA (*) |
|---|---|---|---|---|
| 1 | 2783 | 2846 | 2831 | 2401 (Chr7Clem) |
| 2 | 3513 | 3396 | 3357 | 4741 (Chr2Clem) |
| 3 | 4852 | 4730 | 4847 | 4395 (Chr5Clem) |
| 4 | 2654 | 2935 | 2793 | 2937 (Chr1Clem) |
| 5 | 3222 | 3203 | 3169 | 2816 (Chr3Clem) |
| 6 | 2434 | 2529 | 2405 | 2359 (Chr6Clem) |
| 7 | 2975 | 2819 | 2687 | 2517 (Chr4Clem) |
| 8 | 2389 | 2622 | 2527 | 2218 (Chr8Clem) |
| 9 | 2599 | 2505 | 2544 | 2749 (Chr9Clem) |
| **Total Chr** | **27421** | **27585** | **27160** | **27133** |
| ChrUn | 1837 | 505 | 2317 | 2968 |
| **Total assembly** | **29258** | **28090** | **29477** | **30101** |
| **BUSCOs (eudicots_odb10)** | | | | |
| Complete | 2202 (94.7%) | 2204 (94.8%) | 2171 (93.3%) | 2167 (93.1%) |
| Single | 2144 (92.2%) | 2163 (93.0%) | 2089 (89.8%) | 2124 (91.3%) |
| Duplicated | 58 (2.5%) | 41 (1.8%) | 82 (3.5%) | 43 (1.8%) |
| Fragmented | 83 (3.6%) | 69 (3.0%) | 105 (4.5%) | 73 (3.1%) |
| Missing | 41 (1.7%) | 53 (2.2%) | 50 (2.2%) | 86 (3.8%) |
| **Total** | **2326** | **2326** | **2326** | **2326** |

#### 5) Transposable element and DNA-satellite sequences

**Table 12:** Estimated quantities of repeated sequences in the four studied *Citrus* genomes

| | | CITMI | | CITME | | CITRE | | CITMA | |
|---|---|---|---|---|---|---|---|---|---|
| | | % genome coverage | Number of full copies | % genome coverage | Number of full copies | % genome coverage | Number of full copies | % genome coverage | Number of full copies |
| From Class I | | | | | | | | | |
| LTR Copia (RLC) | *Tork* | 6.70% | 762 | 6.60% | 876 | 6.40% | 702 | 6.30% | 660 |
| | *Sire* | 3.00% | 287 | 5.20% | 857 | 3.30% | 324 | 3.10% | 290 |
| | *Retrofit* | 3.10% | 614 | 3.40% | 801 | 3.30% | 626 | 3.10% | 604 |
| | *Oryco* | 0.70% | 127 | 0.70% | 122 | 0.70% | 94 | 0.70% | 79 |
| | Unknown | 0.30% | 0 | 0.10% | 0 | 0.20% | 0 | 0.10% | 0 |
| LTR Gypsy (RLG) | *Athila* | 10.40% | 910 | 11.90% | 981 | 9.60% | 837 | 9.80% | 770 |
| | *Tat* | 2.30% | 137 | 3.10% | 338 | 2.90% | 310 | 2.20% | 235 |
| | *Crm* | 1.30% | 108 | 1.20% | 134 | 1.20% | 107 | 1.50% | 136 |
| | *Reina* | 0.90% | 133 | 0.90% | 142 | 1.00% | 142 | 0.90% | 128 |
| | *Del* | 0.80% | 93 | 0.70% | 81 | 1.00% | 80 | 0.80% | 53 |
| | *Galadriel* | 0.30% | 67 | 0.40% | 97 | 0.40% | 108 | 0.40% | 94 |
| | Unknown | 0.03% | 0 | 0.04% | 0 | 0.04% | 0 | 0.04% | 0 |
| LINE (RIX) | | 2.10% | 423 | 2.10% | 437 | 2.30% | 401 | 2.20% | 469 |
| SINE (RSX) | | 0.20% | 204 | 0.20% | 194 | 0.20% | 232 | 0.20% | 461 |
| Unclassified Class I | | 0.02% | 0 | 0.01% | 0 | 0.02% | 0 | 0.01% | 0 |
| From Class II | | | | | | | | | |
| TIR elements | *Mutator* (DTM) | 2.70% | 602 | 2.80% | 990 | 2.80% | 593 | 2.60% | 524 |
| | *hAT* (DTA) | 2.60% | 777 | 2.70% | 787 | 2.60% | 741 | 2.60% | 846 |
| | *CACTA* (DTC) | 1.40% | 121 | 1.30% | 125 | 1.40% | 146 | 1.40% | 188 |
| | *Pif-Harbinger* (DTH) | 0.60% | 123 | 0.60% | 114 | 0.60% | 102 | 0.70% | 158 |
| | *Tc1-Mariner* (DTT) | 0.30% | 52 | 0.30% | 38 | 0.30% | 46 | 0.30% | 53 |
| | Unknown (DTX) | 0.01% | 0 | 0.01% | 0 | 0.01% | 0 | 0.01% | 0 |
| MITE | Unknown (DXX-MITE) | 1.50% | 4693 | 1.30% | 3633 | 1.50% | 4334 | 1.50% | 4506 |
| | *hAT* (DTA-MITE) | 0.10% | 125 | 0.20% | 156 | 0.20% | 115 | 0.10% | 126 |
| | *Mutator* (DTM-MITE) | 0.20% | 1465 | 0.20% | 1459 | 0.20% | 1559 | 0.20% | 1653 |
| | *Pif-Harbinger* (DTH-MITE) | 0.02% | 19 | 0.02% | 11 | 0.02% | 22 | 0.02% | 22 |
| Helitron (DHX) | | 0.50% | 76 | 0.50% | 45 | 0.30% | 51 | 0.40% | 49 |
| Unclassified_TE | | 0.70% | 0 | 0.80% | 0 | 0.70% | 0 | 0.80% | 0 |
| DUF_TE (Domains of unknown function in Pfam) | | 0.20% | 0 | 0.20% | 0 | 0.30% | 0 | 0.30% | 0 |
| TOTAL TE | | 42.80% | 11918 | 47.30% | 12418 | 43.60% | 11672 | 42.50% | 12104 |
| DNA-satellite | | 0.50% | 6732 | 1.60% | 24442 | 2.00% | 31488 | 1.00% | 14664 |
| SSR | | 4.50% | | 3.80% | | 4.70% | | 4.70% | |
| TOTAL repeat sequence | | 47.80% | 18650 | 52.70% | 36860 | 50.30% | 43160 | 48.20% | 26768 |

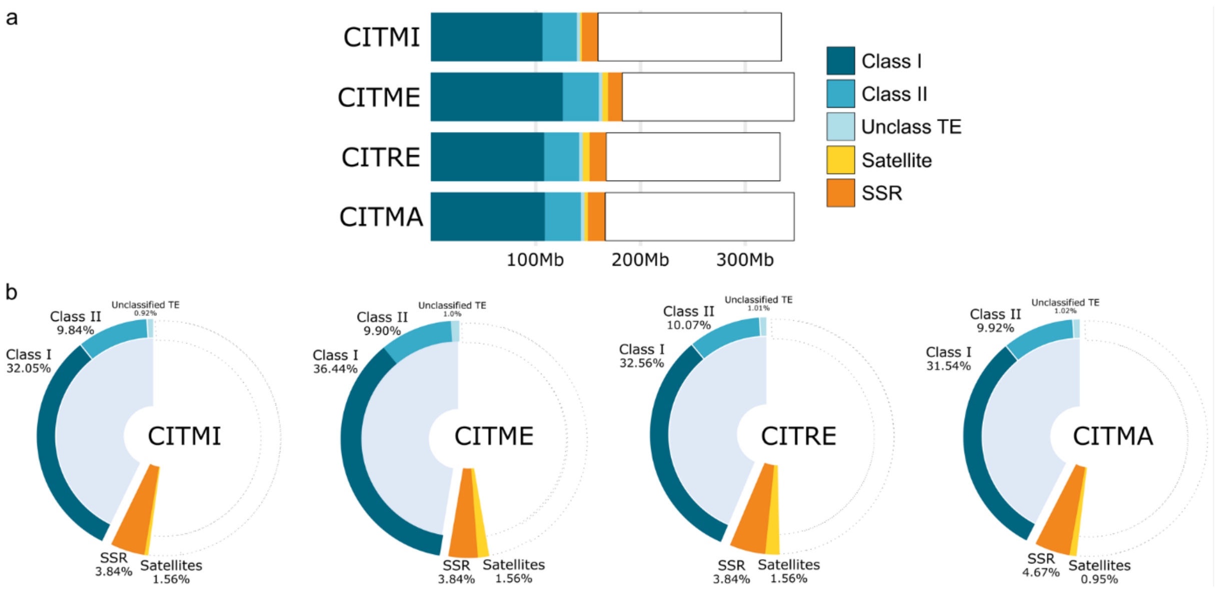

**Figure 6:** Estimated quantities and nature of repeated sequences annotated in the four studied *Citrus* genomes

A: Absolute quantities of repeats in MegaBase. B: Relative quantities of repeats in genomes

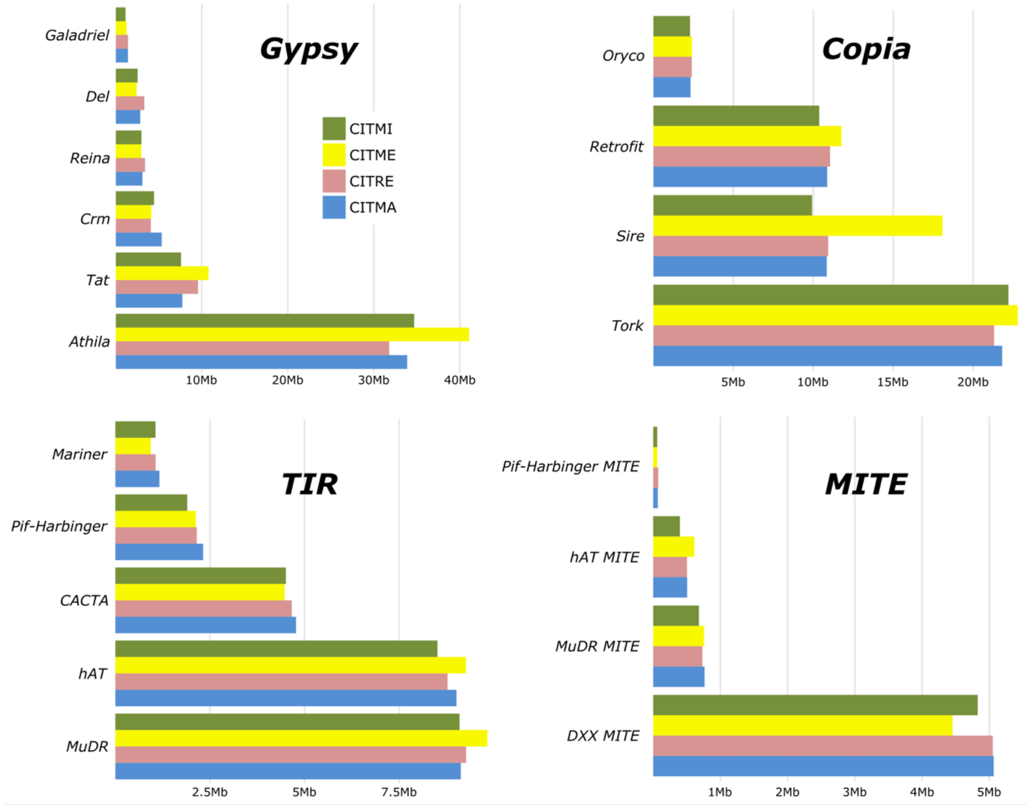

**Figure 7**: Diversity of TE lineages identified in the four *Citrus* genomes

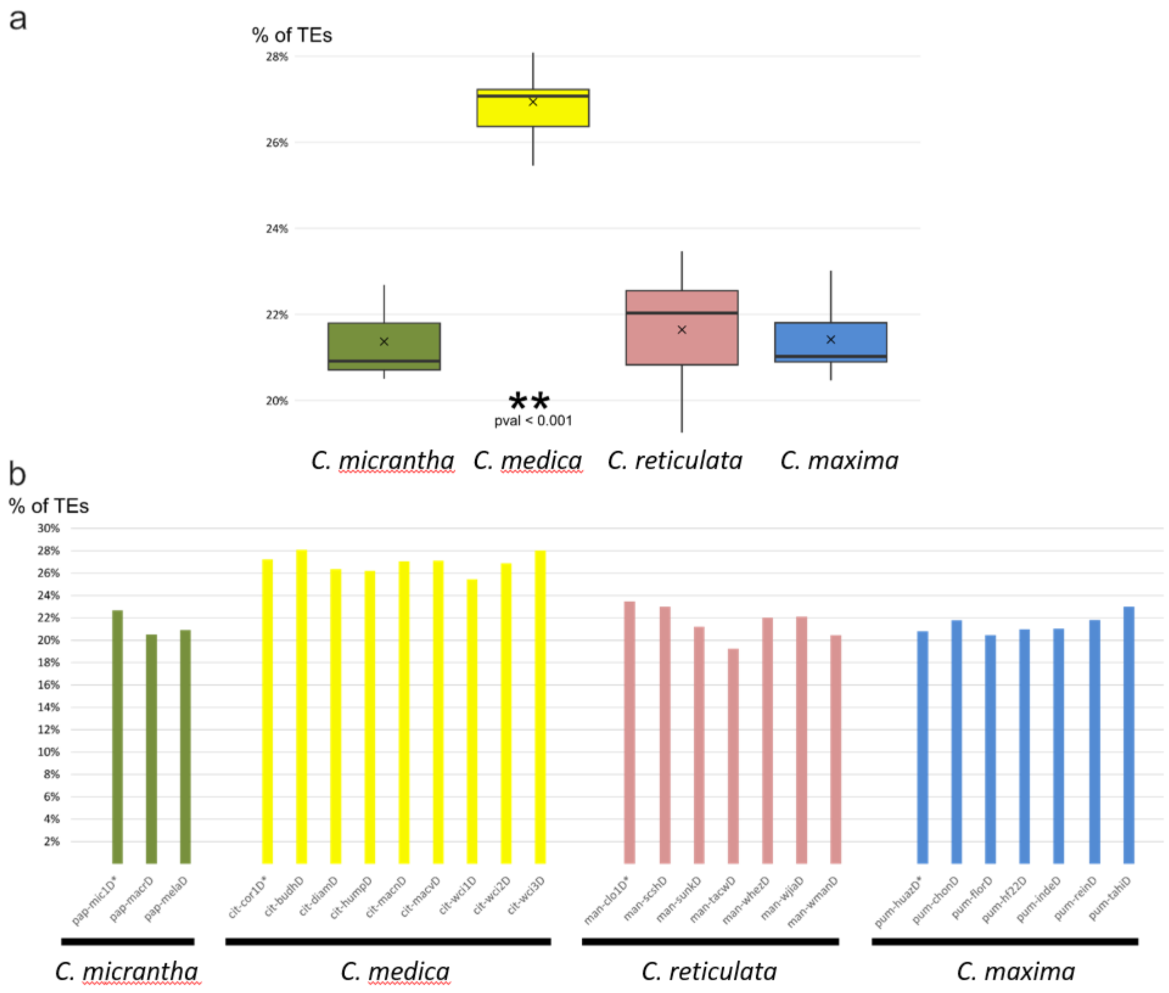

**Figure 8:** Estimated quantities of transposable elements in varieties from the 4 studied Citrus species

a. Boxplot of estimated quantities. The amount of TEs identified in *C. medica* varieties was significantly higher than in varieties of the three other ancestors. Crosses indicate the statistic mean. b. Barplot showing the variation in TE quantities identified in the varieties of each ancestor taxon.

#### 6) PRR analysis

**Table 13:** Quantity of PRR proteins in CITME, CITMI, CITMA and CITRE genomes.

| | CN | CNL | MLO | N | NL | RLK | RLKGNK2 | RLP | RPW8NL | T | TN | TNL | UNKNOWN | TOTAL |
|---|---|---|---|---|---|---|---|---|---|---|---|---|---|---|
| CITMI | 20 | 182 | 17 | 52 | 100 | 97 | 35 | 17 | 6 | 35 | 26 | 73 | 1239 | **1899** |
| CITME | 37 | 139 | 18 | 54 | 83 | 88 | 36 | 25 | 14 | 33 | 15 | 55 | 1030 | **1627** |
| CITRE | 37 | 250 | 17 | 44 | 147 | 102 | 30 | 26 | 12 | 19 | 11 | 56 | 1263 | **2014** |
| CITMA | 41 | 227 | 21 | 48 | 104 | 95 | 32 | 18 | 7 | 14 | 5 | 19 | 1220 | **1851** |

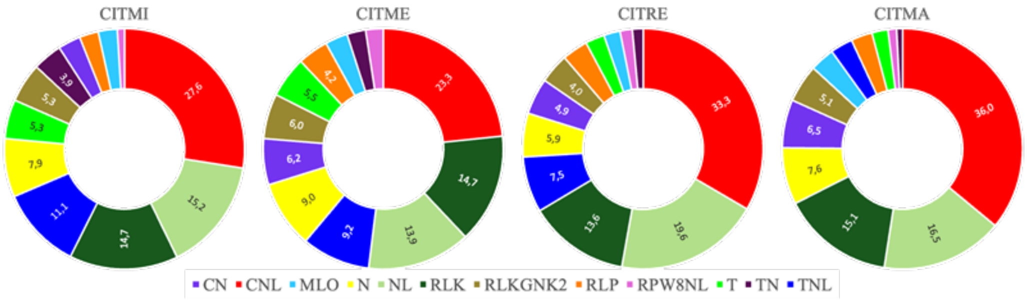

**Figure 9:** Distribution in percent of PRR proteins into the 12 classes as defined by Santana Silva and Micheli (2020).

**Table 14:** Distribution of the PRR of the 12 classes as defined by Santana Silva and Micheli (2020) among the nine chromosomes of each of the four genome assemblies.

| | 1 | 2 | 3 | 4 | 5 | 6 | 7 | 8 | 9 |
|---|---|---|---|---|---|---|---|---|---|
| CITMI | 80 (13.51%) | 66 (11.15%) | 154 (26.01%) | 11 (1.86%) | 77 (13.01%) | 12 (2.03%) | 146 (24.66%) | 19 (3.21%) | 27 (4.56%) |
| CITME | 69 (11.62%) | 52 (8.75%) | 173 (29.12%) | 15 (2.53%) | 109 (18.35%) | 11 (1.85%) | 114 (19.19%) | 35 (5.89%) | 16 (2.69%) |
| CITRE | 55 (9.70%) | 169 (8.64%) | 47 (27.51%) | 18 (2.65%) | 94 (21.69%) | 20 (3.70%) | 137 (19.58%) | 12 (4.41%) | 36 (2.12%) |
| CITMA | 55 (9.35%) | 49 (28.74%) | 156 (7.99%) | 15 (3.06%) | 123 (15.99%) | 21 (3.40%) | 111 (23.30%) | 25 (2.04%) | 12 (6.12%) |

**Table 15.** Clustering of PRR genes (including the UNKNOWN class) of the CITME, CITMI, CITMA and CITRE genomes.

| | | | | | | | | | | | |
|---|---|---|---|---|---|---|---|---|---|---|---|
| | **CITMI** | | | | | | | | | | |
| | chr1 | chr2 | chr3 | chr4 | chr5 | chr6 | chr7 | chr8 | chr9 | chrUN | Total-UN |
| Total amount of genes | 174 | 241 | 339 | 82 | 201 | 86 | 365 | 81 | 118 | 212 | 1687 |
| Number of clusters | 31 | 40 | 58 | 18 | 43 | 15 | 55 | 16 | 26 | | 302 |
| Number of genes in clusters | 151 | 211 | 300 | 60 | 172 | 62 | 348 | 61 | 100 | - | 1465 |
| % clustered genes | 86.8 | 87.6 | 88.5 | 73.2 | 85.6 | 72.1 | 95.3 | 75.3 | 84.7 | **-** | **86.8** |
| | **CITME** | | | | | | | | | | |
| | chr1 | chr2 | chr3 | chr4 | chr5 | chr6 | chr7 | chr8 | chr9 | chrUN | Total-UN |
| Total amount of genes | 187 | 176 | 332 | 85 | 232 | 89 | 311 | 112 | 95 | 8 | 1619 |
| Number of clusters | 35 | 33 | 59 | 21 | 52 | 16 | 60 | 19 | 19 | | 314 |
| Number of genes in clusters | 163 | 139 | 298 | 63 | 218 | 63 | 288 | 88 | 71 | - | 1391 |
| % clustered genes | 87.2 | 79.0 | 89.8 | 74.1 | 94.0 | 70.8 | 92.6 | 78.6 | 74.7 | **-** | **85.9** |
| | **CITRE** | | | | | | | | | | |
| | chr1 | chr2 | chr3 | chr4 | chr5 | chr6 | chr7 | chr8 | chr9 | chrUN | Total-UN |
| Total amount of genes | 173 | 171 | 342 | 92 | 251 | 97 | 256 | 96 | 114 | 422 | 1592 |
| Number of clusters | 29 | 32 | 59 | 20 | 50 | 16 | 46 | 20 | 26 | | 298 |
| Number of genes in clusters | 146 | 137 | 296 | 71 | 233 | 71 | 230 | 78 | 99 | - | 1361 |
| % clustered genes | 84.4 | 80.1 | 86.5 | 77.2 | 92.8 | 73.2 | 89.8 | 81.3 | 86.8 | **-** | **85.5** |
| | **CITMA** | | | | | | | | | | |
| | chr1 | chr2 | chr3 | chr4 | chr5 | chr6 | chr7 | chr8 | chr9 | chrUN | Total-UN |
| | CITMA4 | CITMA2 | CITMA5 | CITMA7 | CITMA3 | CITMA6 | CITMA1 | CITMA8 | CITMA9 | | |
| Total amount of genes | 150 | 406 | 180 | 102 | 224 | 94 | 349 | 115 | 133 | 98 | 1753 |
| Number of clusters | 32 | 72 | 31 | 20 | 46 | 23 | 58 | 23 | 27 | | 332 |
| Number of genes in clusters | 127 | 368 | 126 | 83 | 202 | 77 | 329 | 107 | 109 | - | 1528 |
| % clustered genes | 84.7 | 90.6 | 70.0 | 81.4 | 90.2 | 81.9 | 94.3 | 93.0 | 82.0 | **-** | **87.2** |

#### **Table 16**: Duplications of PRR genes

| CITMI | | CITME | | CITRE | | CITMA | |
|---|---|---|---|---|---|---|---|
| | | | | CN | | CN | |
| | | | | CITRE_005t013410 | CITRE_005t014620 | CITMA_003t008040 | CITMA_003t010190 |
| CNL | | CNL | | CNL | | CNL | |
| CITMI_003t026020 | CITMI_003t026910 | CITME_007t015440 | CITME_007t016690 | CITRE_007t015570 | CITRE_007t016250 | CITMA_001t017800 | CITMA_001t018310 |
| CITMI_007t016080 | CITMI_007t016470 | CITME_007t015870 | CITME_007t016470 | | | CITMA_001t017720 | CITMA_001t018270 |
| CITMI_007t014660 | CITMI_007t015030 | | | | | CITMA_001t018430 | CITMA_001t020050 |
| CITMI_007t014520 | CITMI_007t014910 | | | | | CITMA_001t017740 | CITMA_001t018310 |
| CITMI_007t016310 | CITMI_007t016320 | | | | | CITMA_002t016260 | CITMA_002t016720 |
| CITMI_007t016160 | CITMI_007t016470 | | | | | CITMA_001t016150 | CITMA_001t018230 |
| | | | | | | CITMA_001t010610 | CITMA_003t005850 |
| | | | | | | CITMA_001t010610 | CITMA_003t003680 |
| | | | | | | CITMA_001t010610 | CITMA_003t003830 |
| | | | | | | CITMA_001t018410 | CITMA_001t020150 |
| | | | | | | CITMA_001t016050 | CITMA_001t018410 |
| | | | | | | CITMA_001t017850 | CITMA_001t018410 |
| | | | | | | CITMA_001t017850 | CITMA_001t018270 |
| | | | | | | CITMA_001t017820 | CITMA_001t020010 |
| | | | | | | CITMA_003t003680 | CITMA_003t005850 |
| | | | | | | CITMA_003t003830 | CITMA_003t005850 |
| | | | | | | CITMA_001t010450 | CITMA_003t003820 |
| | | | | | | CITMA_003t008180 | CITMA_003t010020 |
| | | | | | | CITMA_003t008130 | CITMA_003t009890 |
| | | | | | | CITMA_003t008130 | CITMA_003t010270 |
| | | | | | | CITMA_001t010540 | CITMA_003t005790 |
| | | | | | | CITMA_001t010770 | CITMA_003t005650 |
| | | | | | | CITMA_003t003820 | CITMA_003t005650 |
| | | | | | | CITMA_002t015580 | CITMA_002t016310 |
| | | | | | | CITMA_001t010770 | CITMA_003t003750 |
| MLO | | MLO | | MLO | | MLO | |
| CITMI_005t031570 | CITMI_007t000180 | CITME_004t005340 | CITME_008t021670 | CITRE_005t031050 | CITRE_007t000180 | CITMA_001t000180 | CITMA_003t027480 |
| CITMI_004t005520 | CITMI_008t019210 | CITME_005t031420 | CITME_007t000170 | CITRE_004t005530 | CITRE_008t020610 | CITMA_007t006180 | CITMA_008t017880 |
| | | | | | | N | |
| | | | | | | CITMA_001t010560 | CITMA_003t003720 |
| | | | | | | NL | |
| | | | | | | CITMA_003t006310 | CITMA_009t011370 |
| | | | | | | CITMA_003t004510 | CITMA_009t011370 |
| | | | | | | CITMA_008t010190 | CITMA_009t016270 |
| | | | | | | CITMA_001t018010 | CITMA_001t018290 |
| | | | | | | CITMA_001t016400 | CITMA_001t017880 |
| RLK | | RLK | | RLK | | RLK | |
| CITMI_002t014690 | CITMI_002t028830 | CITME_001t015860 | CITME_002t027420 | CITRE_001t014640 | CITRE_002t027370 | CITMA_002t014590 | CITMA_002t041020 |
| CITMI_002t002490 | CITMI_004t026270 | CITME_002t001870 | CITME_004t029080 | CITRE_002t001940 | CITRE_004t027690 | CITMA_001t017260 | CITMA_004t002340 |
| CITMI_002t014730 | CITMI_002t028800 | CITME_002t018330 | CITME_002t027490 | CITRE_002t013780 | CITRE_002t027370 | CITMA_002t022850 | CITMA_002t041010 |
| CITMI_001t015610 | CITMI_002t028790 | CITME_001t015860 | CITME_002t018330 | | | CITMA_002t014600 | CITMA_002t041060 |
| CITMI_001t015480 | CITMI_002t028800 | CITME_001t015780 | CITME_002t027430 | | | CITMA_002t022630 | CITMA_002t041020 |
| | | CITME_002t018300 | CITME_002t027430 | | | | |
| RLKGNK2 | | RLKGNK2 | | RLKGNK2 | | RLKGNK2 | |
| CITMI_002t007650 | CITMI_004t021240 | CITME_002t017900 | CITME_002t027030 | CITRE_003t004900 | CITRE_004t022710 | CITMA_002t007100 | CITMA_007t018790 |
| CITMI_003t005000 | CITMI_004t021240 | CITME_003t004990 | CITME_004t023980 | CITRE_002t006900 | CITRE_004t022710 | CITMA_005t005070 | CITMA_007t018790 |
| CITMI_002t007650 | CITMI_003t005000 | CITME_002t006790 | CITME_003t004990 | CITRE_002t014580 | CITRE_002t027020 | CITMA_002t007100 | CITMA_005t005070 |
| | | | | CITRE_002t006900 | CITRE_003t004900 | | |
| T | | T | | T | | T | |
| CITMI_003t024480 | CITMI_003t025170 | CITME_008t011740 | CITME_008t013560 | CITRE_003t023260 | CITRE_003t024290 | CITMA_002t021430 | CITMA_002t041810 |
| CITMI_001t014090 | CITMI_002t029620 | CITME_001t014500 | CITME_002t028380 | CITRE_001t013330 | CITRE_002t028120 | | |
| TNL | | TNL | | TNL | | TNL | |
| CITMI_003t027290 | CITMI_003t028150 | CITME_003t026460 | CITME_003t027310 | CITRE_003t023160 | CITRE_003t024520 | CITMA_002t017110 | CITMA_005t024710 |
| CITMI_003t027280 | CITMI_003t027940 | | | | | CITMA_004t016290 | CITMA_004t016350 |
| | | | | | | CITMA_004t016290 | CITMA_005t019670 |
| RLP | | | | | | | |
| CITMI_007t020880 | CITMI_007t021600 | | | | | | |

#### 7) Variant calling from resequencing data

**Table 17:** Intraspecific and total diversity from WGS resequencing data of 55 accessions

| Reference genomes | | CITMI | CITME | CITRE | CITMA |
|---|---|---|---|---|---|
| Intraspecific polymorphisms | SNP | 6668709 | 3493927 | 7550888 | 6146739 |
| | Indel | 766230 | 430808 | 920361 | 758788 |
| | Total | 7434939 | 3924735 | 8471249 | 6905527 |
| | Het / kb | 5.10 | 0.95 | 4.17 | 3.70 |
| Four ancestral species representatives | SNP | 18966636 | 20187996 | 19805888 | 19575468 |
| | Indel | 2888875 | 3175663 | 3017540 | 2943651 |
| | Total | 21855511 | 23363659 | 22823428 | 22519119 |
| | Het / kb | 4.95 | 4.44 | 5.03 | 4.79 |
| All accessions | SNP | 20072074 | 21504545 | 21317941 | 20770049 |
| | Indel | 3111963 | 3435129 | 3290700 | 3175025 |
| | Total | 23184037 | 24939674 | 24608641 | 23945074 |
| | Het /kb | 6.85 | 6.36 | 6.97 | 6.83 |
| Add mixed | Het / kb | 11.93 | 11.48 | 12.14 | 12.27 |

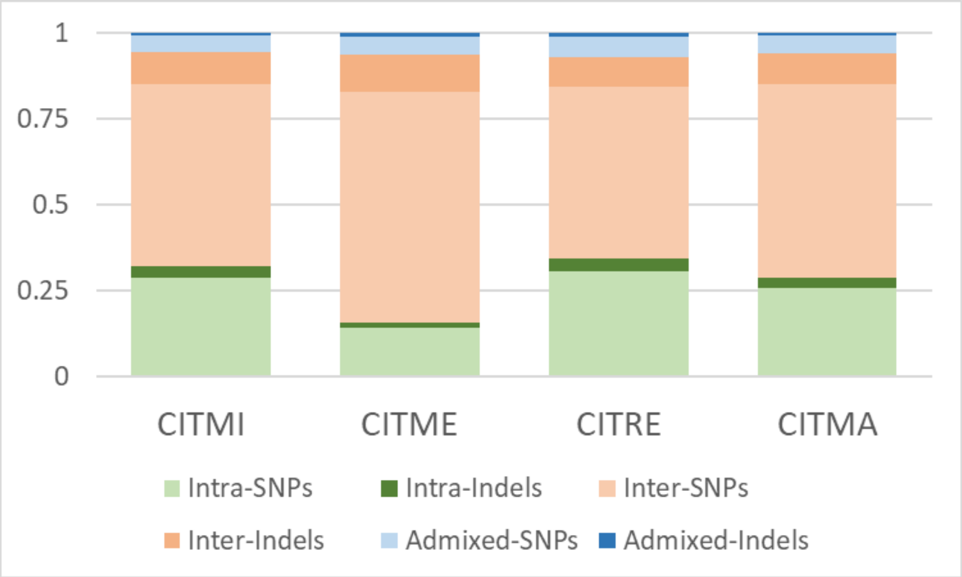

**Figure 10:** Relative contribution of intraspecific, interspecific and admixed accessions to the whole diversity revealed by the variant calling in the four assemblies

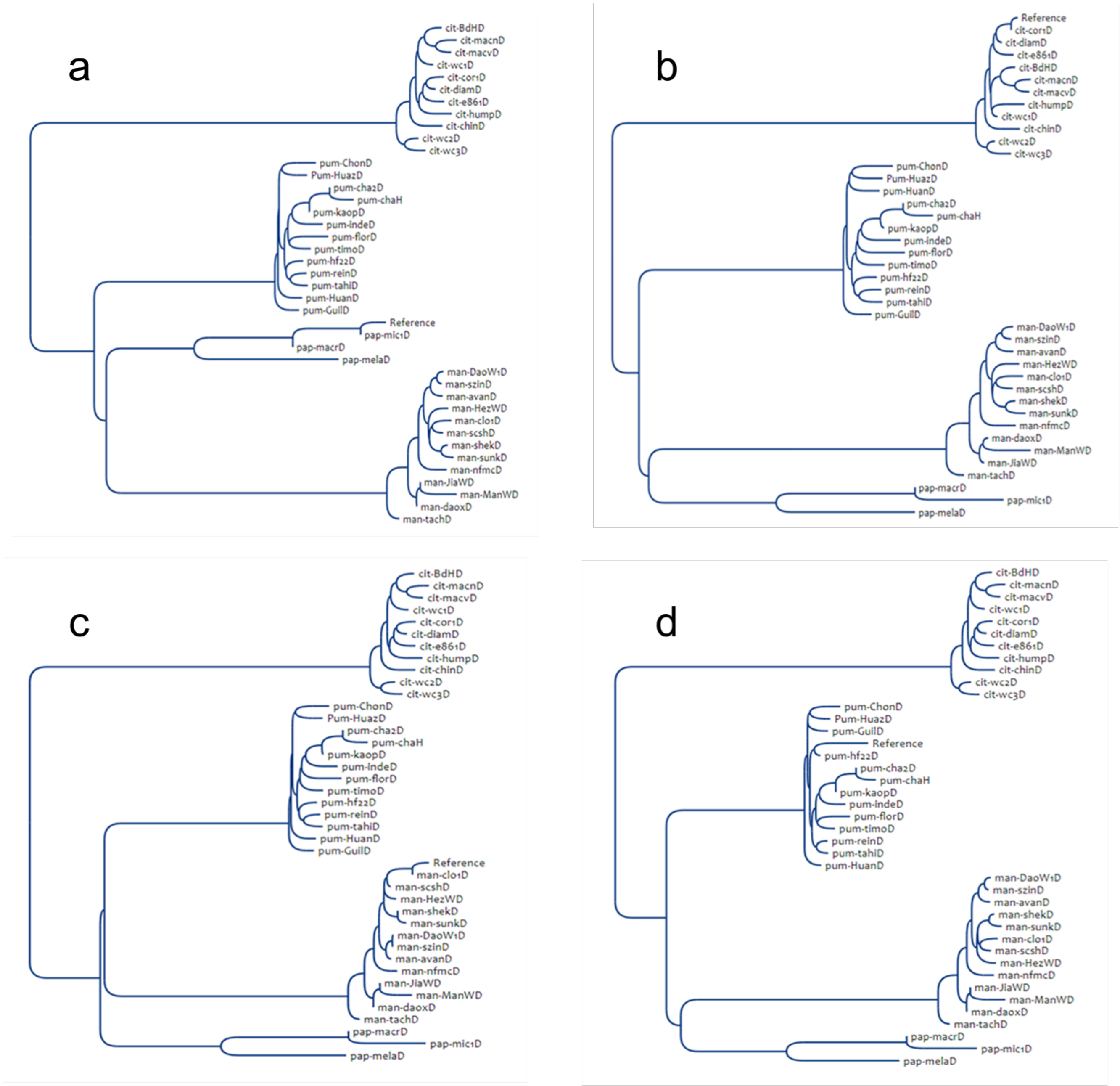

**Figure 11:** Phylogenetic tree based on the minimum-evolution principle (Desper and Gascuel, 2002) of the accession representative of the four ancestors based on diallelic SNPs sampled for at least 1Kb distance between successive markers.

a: CITMI reference genome (228646 SNPs); b: CITME (249231 SNPs); c: CITRE (235801 SNPs); d: CITMA (230638 SNPs).

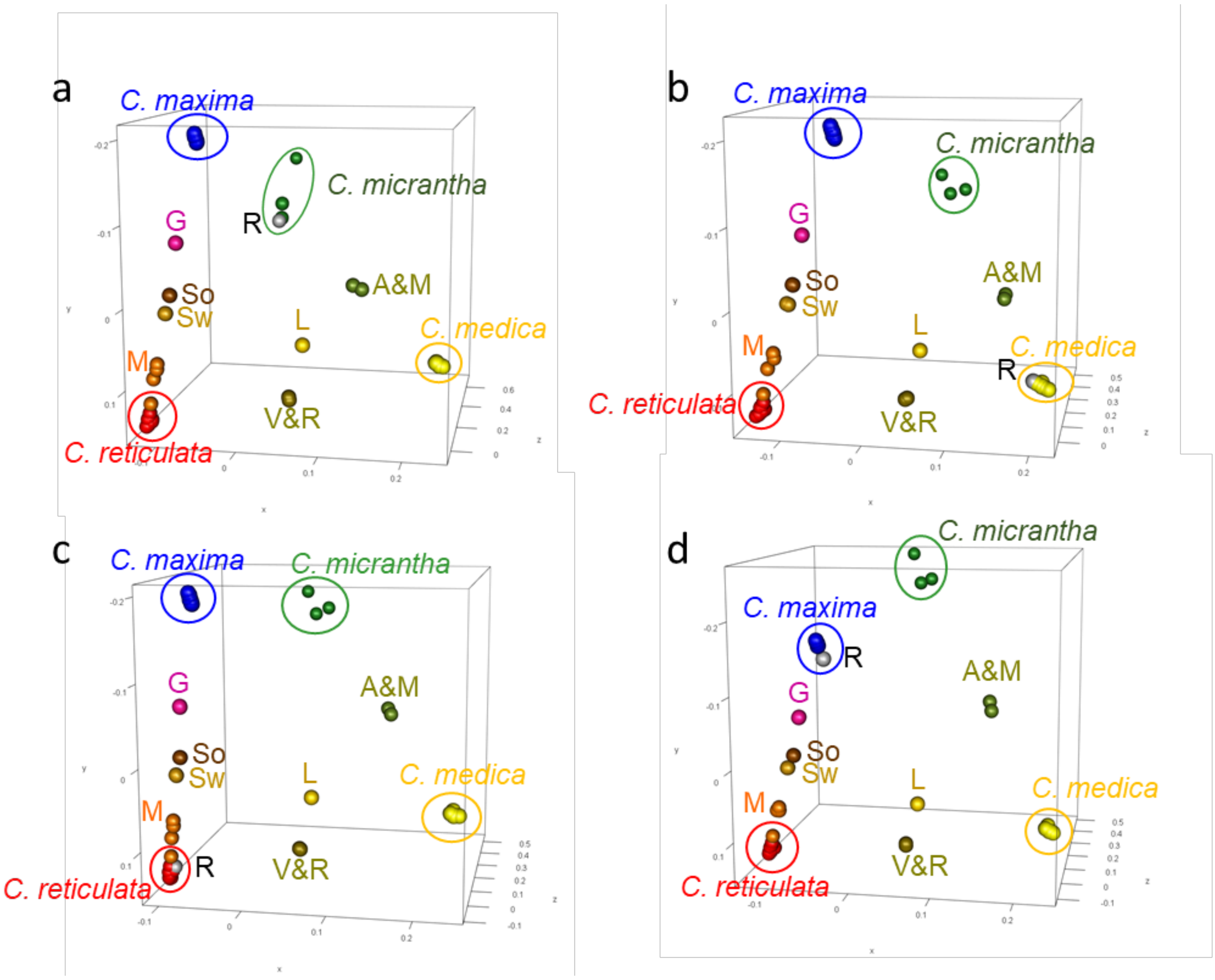

**Figure 12:** Factorial analysis based on diallelic SNPs sampled for at least 1Kb distance between successive markers

a: CITMI reference genome (230601 SNPs); b: CITME (250358 SNPs); c: CITRE (238852 SNPs); d: CITMA (233539 SNPs). R: reference assembly, G: grapefruit, A&M: 'Alemow' and 'Mexican' lime, So: sour orange, Sw: sweet orange, L: 'Eureka' lemon, V&R: 'Volkamer' lemon and 'Rangpur' lime, Man: mandarins and 'Clementine'.

#### 8) Analysis of interspecific mosaic of cultivated Citrus

**Text 1:** detailed method for the analysis of interspecific mosaic structure

The approach we have used is derived from the one described by Ahmed et al. (2019) for GBS data and polyploid genotypes. It is based on the identification of diagnostic diallelic SNP markers that fully differentiate one ancestral species from all others. The frequency of the diagnostic alleles along the genome allow then to identify the contribution of each ancestral species to the analysed accession. Briefly the different steps are as following:

1. *Search of diagnostic SNPs of the four ancestral species*

The identification of the diagnostic markers for each of the four ancestral species was based on GST parameter estimations (Nei, 1973). GST is the coefficient of gene differentiation which measures differentiation among sub-populations. It is equivalent to Wright's FST for biallelic markers and ranges from zero to one for complete differentiation between populations. For each taxon, *GST* estimation was performed considering two sub-populations: (1) the taxon concerned ($T_i$) and (2) a theoretical population of the three other taxa ($T_{-i}$). Analyses were performed from the estimated allele frequency of each taxon considering the same population size for each taxon to estimate the frequency of the two sub-populations ($T_i$ and $T_{-i}$) and the frequency of the whole population (*Tot*):

$$GST = \frac{He_{Tot} - \frac{(He_{Ti} + He_{T-i})}{2}}{He_{Tot}}$$

where *He* is the expected proportion of heterozygous loci per individual under Hardy–Weinberg equilibrium ($He = 1 - \Sigma\, p_i^2$, $p_i$ is the frequency of a given allele in the population or sub-population considered).

Previous studies (Oueslati et al., 2017; Wu et al., 2018) have shown that some varieties of the modern horticultural groups derived from the four ancestral species were not pure representative of theses ancestral species, but have interspecific introgressions in heterozygosity and even sometime in homozygosity. The implementation of the set of DSNPs of ancestral species require therefore the identification of theses introgressed genomic regions and there removing before estimating allele frequencies of the ancestors. The identification of the interspecific introgressed areas for each accession was based on the pattern of two parameters along the genome using consecutive sliding windows of 200kb: (1) the average heterozygosity estimated from diallelic SNPs and (2) the similarity of the accession to the centroid of each of the four horticultural representative groups. Introgressed areas display significant coordinated changes of these patterns according to the level of differentiation between the two taxa involved. To better contrast the pattern discontinuities, we selected at this step SNPs with GST >0.5 for one of the horticultural representative groups of the ancestral taxon. An example of the identification of introgressed region in the Chr8 of CITRE assembly is given for three mandarins (Figure). For Nan Feng Mi Chu mandarin, coordinated variations are observed at the beginning of the chromosome suggesting alternate introgression of C. maxima in heterozygosity and homozygosity. Therefore, the genotyping data in the corresponding region (0 to 6.5 Mb) of this accession were changed on missing data. On the same way, two region of C. maxima introgression were identified for Szinkom mandarin and corresponding genotyping data (0 to 5.8 Mb and 22.1 to 26.8 Mb) suppressed. No evidence of introgression was found for Shekwasha mandarin and its whole genotyping data were used for DSNPs search. Once the interspecific introgressions were removed (considered as missing data) for the 40 representative accessions, the allelic frequencies in the four ancestral taxa and the GST parameter between each ancestral taxon and the three others were estimated again. We then considered SNPs with a GST value =1 as diagnostic markers of the considered taxon. All this process was implemented on SniPlay web tool (https://sniplay.southgreen.fr/cgi-bin/home.cgi) for analysis from the vcf file obtained from the variant calling.

2. *Analysis of the interspecific mosaic structure of cultivated citrus*

The first step aimed to estimate the doses of the ancestral genome fragments along the genome. For each ancestral species, the citrus genome was segmented in windows of 20 consecutive DSNPs and the doses (0, 1 and 2) of the ancestral species considered were estimated for each window by maximum likelihood analysis according to the observed numbers of allele specific to the considered taxa and alternative ones. During this step, the number and position of windows varied between the ancestral species according to the density and positions of the DSNPs of each of the four considered species. To integrate the information obtained for the four ancestral species, we sub-divided the genome into successive fragments of 100 kb. For each ancestor and for each genomic fragment, the corresponding window of 20 DSNPs was identified and the ancestral dose of this window was attributed to the genomic fragment. A non-phased representation of karyotypes with two chromosomes was then generated from the ancestral doses of each genome fragment. For a given genome fragment, if the sum of the allelic doses of the different ancestors was over 2, the phylogenomic origin of the fragment was considered as undefined. When phased haplotypes were known for the parental genomes, we proposed manually phased karyotypes for the concerned accession, assuming the lower number of recombination events as the best model. Graphical display were generated directly by the SniPlay web tool (https://sniplay.southgreen.fr/cgi-bin/home.cgi) or with GEMO for manually phased representation (https://gemo.southgreen.fr/).

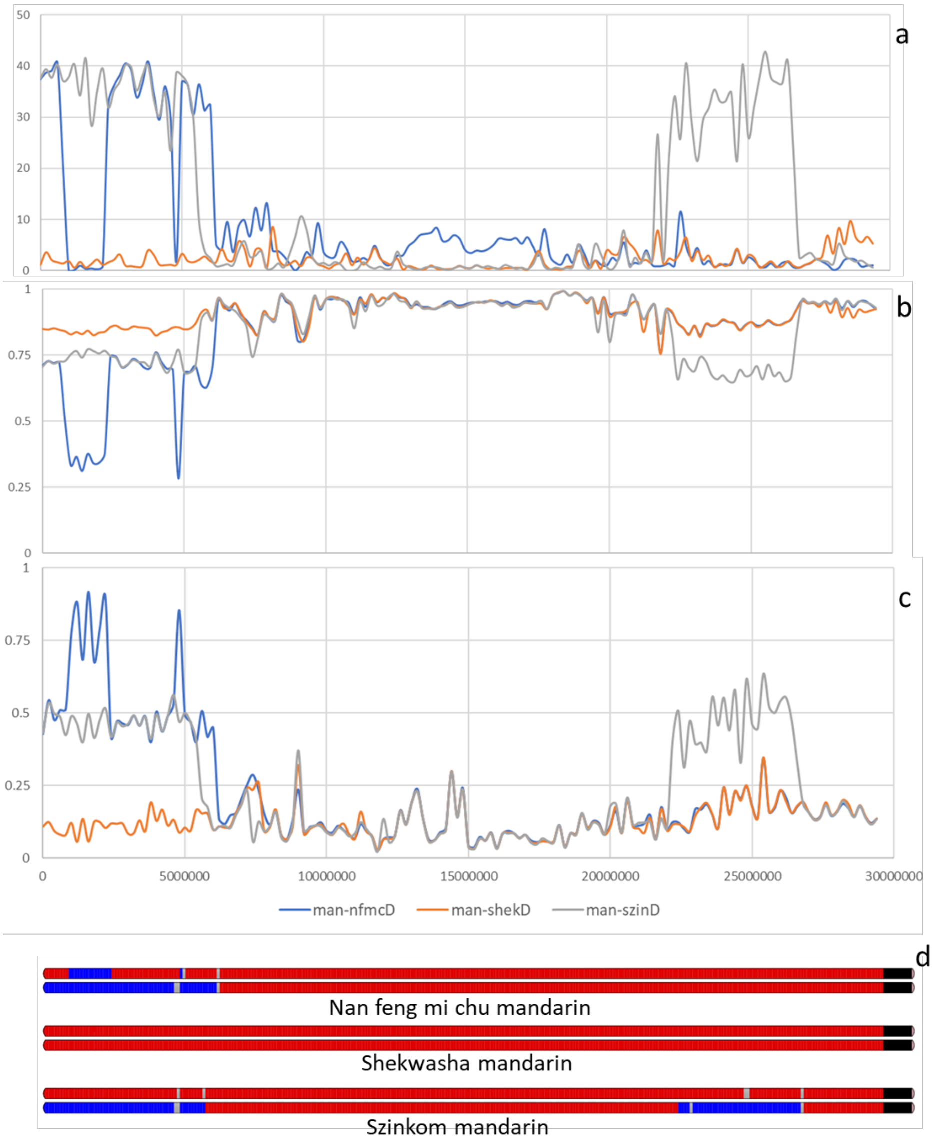

**Figure 13:** Illustration of the identification of introgressed regions in the Chr 8 of three accessions representative of C. reticulata using sliding windows of 200kb

a: % of heterozygosity for diallelic SNP with GST value > 0.5 for one of the four ancestral species; b: similarity with the centroid of *C. reticulata* estimated from diallelic SNPs with GST value > 0.5 for *C. reticulata*; similarity with the centroid of *C. maxima* estimated from diallelic SNPs with GST value > 0.5 for *C. reticulata* or *C. maxima*.

**Table 18:** Numbers of dSNPs of the four ancestral species on the chromosome of the four genome assemblies

| CITMI | CITMI_001 | CITMI_002 | CITMI_003 | CITMI_004 | CITMI_005 | CITMI_006 | CITMI_007 | CITMI_008 | CITMI_009 | Total |
|---|---|---|---|---|---|---|---|---|---|---|
| *C. medica* | 122691 | 136494 | 171377 | 109472 | 98915 | 105536 | 102051 | 98151 | 103008 | 1047695 |
| *C. reticulata* | 71676 | 74001 | 101779 | 59730 | 51455 | 60499 | 54923 | 56053 | 52052 | 582168 |
| *C. maxima* | 41826 | 42078 | 52279 | 33841 | 25939 | 34620 | 30512 | 28146 | 24747 | 313988 |
| *C. micrantha* | 24126 | 30386 | 40122 | 25123 | 20628 | 24898 | 20764 | 22140 | 18615 | 226802 |
| | 260319 | 282959 | 365557 | 228166 | 196937 | 225553 | 208250 | 204490 | 198422 | **2170653** |

| CITME | CITME_001 | CITME_002 | CITME_003 | CITME_004 | CITME_005 | CITME_006 | CITME_007 | CITME_008 | CITME_009 | Total |
|---|---|---|---|---|---|---|---|---|---|---|
| C. medica | 124455 | 122334 | 170083 | 120776 | 103340 | 110646 | 100446 | 103638 | 105183 | 1060901 |
| C. reticulata | 72528 | 69625 | 102190 | 64752 | 52838 | 64194 | 55754 | 58679 | 53519 | 594079 |
| C. maxima | 43376 | 40223 | 53690 | 37584 | 28054 | 36405 | 31445 | 30256 | 25483 | 326516 |
| C. micrantha | 32935 | 37956 | 55287 | 36191 | 29192 | 35609 | 28563 | 34359 | 26851 | 316943 |
| | 273294 | 270138 | 381250 | 259303 | 213424 | 246854 | 216208 | 226932 | 211036 | **2298439** |

| CITRE | CITRE_001 | CITRE_002 | CITRE_003 | CITRE_004 | CITRE_005 | CITRE_006 | CITRE_007 | CITRE_008 | CITRE_009 | Total |
|---|---|---|---|---|---|---|---|---|---|---|
| *C. medica* | 122872 | 128177 | 177656 | 124665 | 105833 | 110067 | 105191 | 101996 | 107463 | 1083920 |
| *C. reticulata* | 67550 | 65247 | 95451 | 61486 | 51442 | 57863 | 52118 | 54079 | 50554 | 555790 |
| *C. maxima* | 41615 | 40516 | 53367 | 37680 | 28059 | 35669 | 31452 | 29252 | 26289 | 323899 |
| *C. micrantha* | 31685 | 38105 | 53942 | 35189 | 28567 | 33810 | 27590 | 32613 | 26372 | 307873 |
| | 263722 | 272045 | 380416 | 259020 | 213901 | 237409 | 216351 | 217940 | 210678 | **2271482** |

| CITMA | Chr1Clem | Chr2Clem | Chr3Clem | Chr4Clem | Chr5Clem | Chr6Clem | Chr7Clem | Chr8Clem | Chr9Clem | |
|---|---|---|---|---|---|---|---|---|---|---|
| | CITMA_004 | CITMA_002 | CITMA_005 | CITMA_007 | CITMA_003 | CITMA_006 | CITMA_001 | CITMA_008 | CITMA_009 | Total |
| *C. medica* | 107467 | 186565 | 190118 | 85803 | 97268 | 100896 | 112755 | 66905 | 112821 | 1060598 |
| *C. reticulata* | 64305 | 94008 | 110898 | 51756 | 51431 | 59408 | 61242 | 39546 | 57364 | 589958 |
| *C. maxima* | 31730 | 48128 | 53233 | 24823 | 23444 | 27791 | 30995 | 16877 | 25194 | 282215 |
| *C. micrantha* | 28871 | 53904 | 54992 | 25048 | 25543 | 27851 | 30576 | 19400 | 28498 | 294683 |
| | 232373 | 382605 | 409241 | 187430 | 197686 | 215946 | 235568 | 142728 | 223877 | **2227454** |

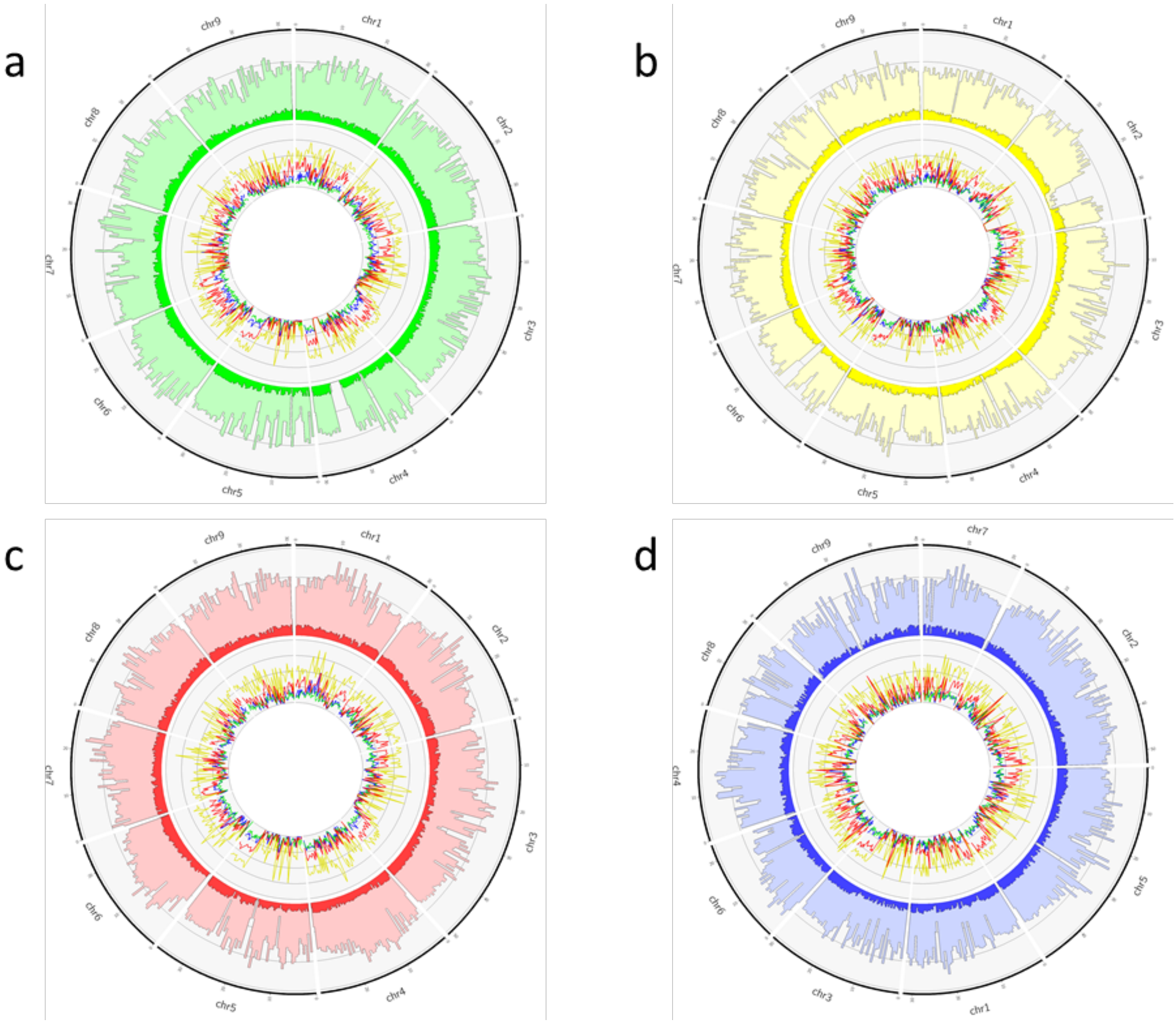

**Figure 14:** Distribution of SNPs, Indels and DSNPs along the four genome assemblies

External ring: distribution of SNPs (light color) and Indels (dark color), scale: 0-120 /kb; Internal ring: distribution dSNPs of the four ancestral species along the genome assemblies, green: *C. micrantha;* yellow: *C. medica;* red: *C. reticulata*; blue: *C. maxima*; scale 0-8 DSNP / kb). a: CITMI reference genome, b: CITME; c: CITRE; d: CITMA. The numbering of the CITMA chromosomes is the one of Wang et al (2017) publication but they are ordered according to the other three genomes.

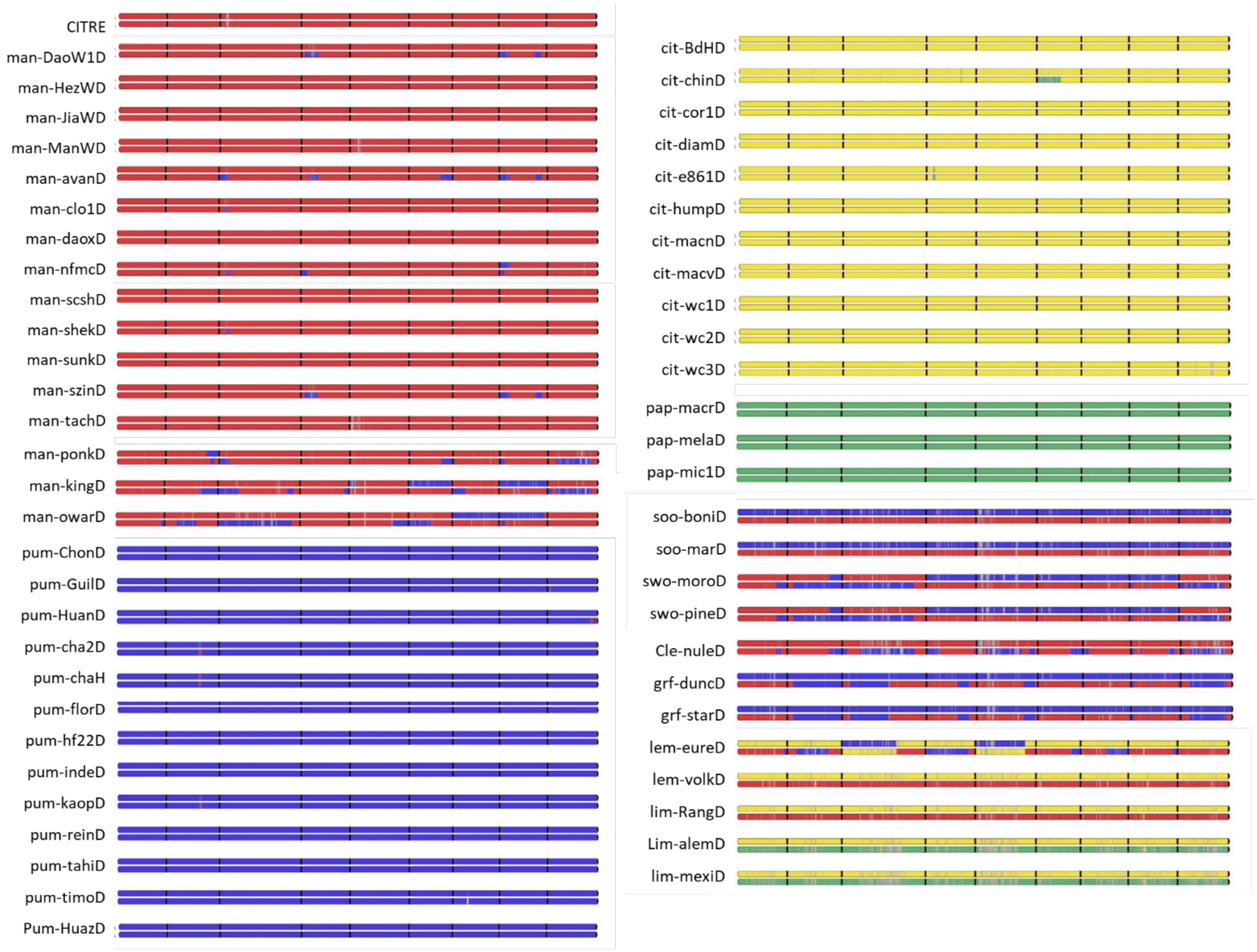

**Figure 15**: Interspecific mosaic of 55 citrus cultivars according CITRE genome assembly
red: *C. reticulata*; blue: *C. maxima*; yellow: *C. medica*; green: *C. micrantha*; grey: undetermined

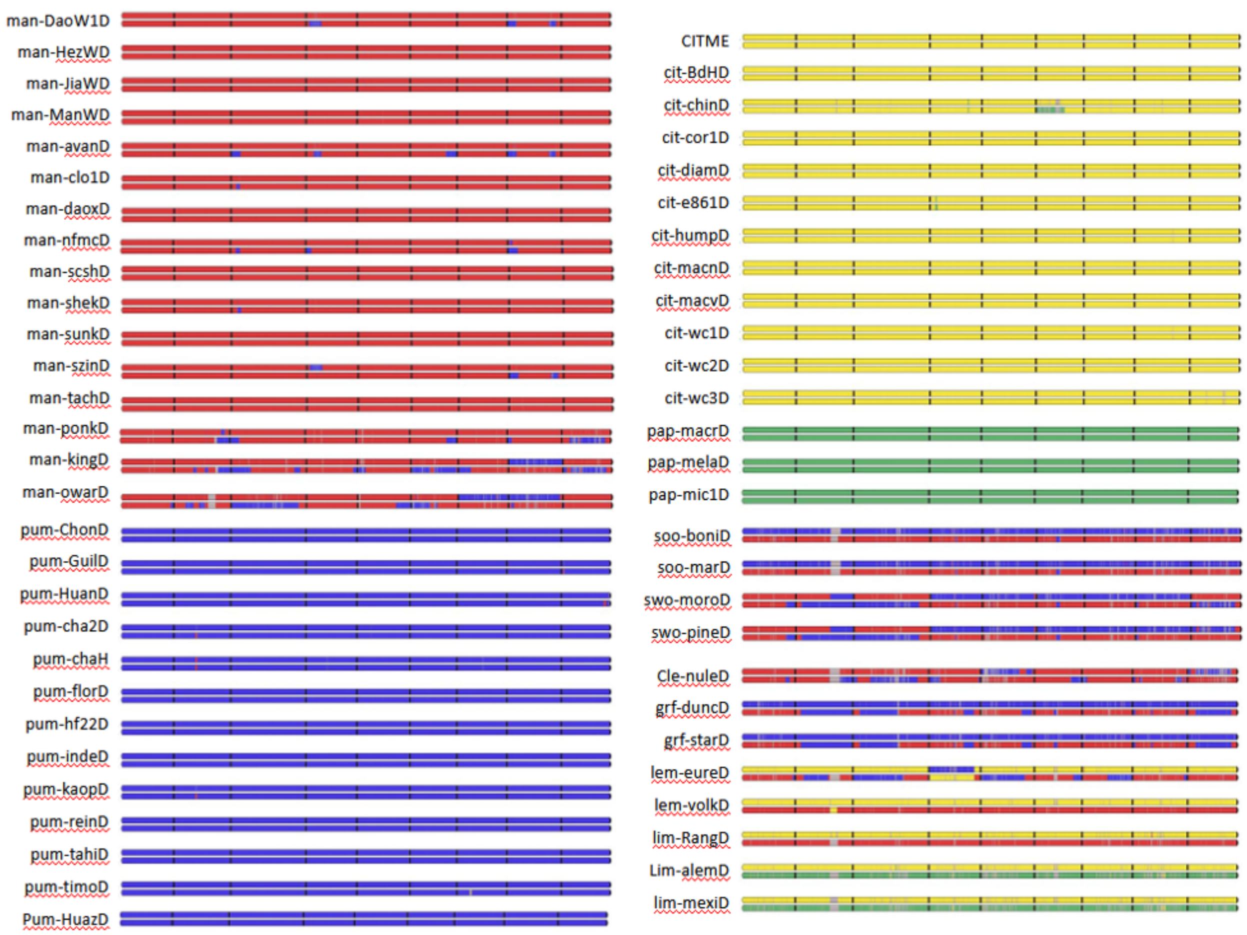

**Figure 16**: Interspecific mosaic of 55 citrus cultivars according CITME genome assembly
red: *C. reticulata*; blue: *C. maxima*; yellow: *C. medica*; green: *C. micrantha*; grey: undetermined

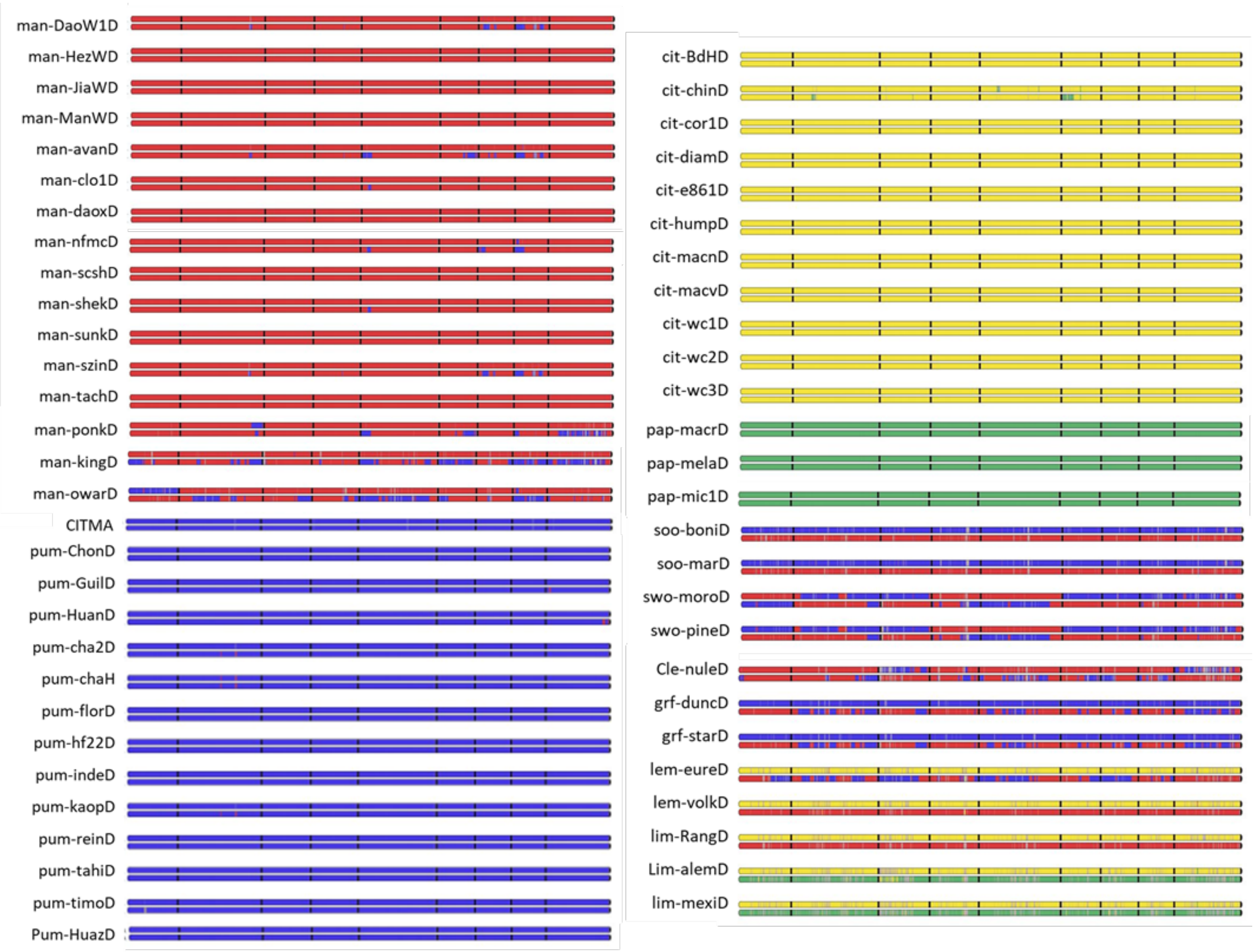

**Figure 17:** Interspecific mosaic of 55 citrus cultivars according CITMA genome assembly
red: *C. reticulata*; blue: *C. maxima*; yellow: *C. medica*; green: *C. micrantha*; grey: undetermined.
The numbering of the CITMA chromosomes is the one of Wang et al (2017).

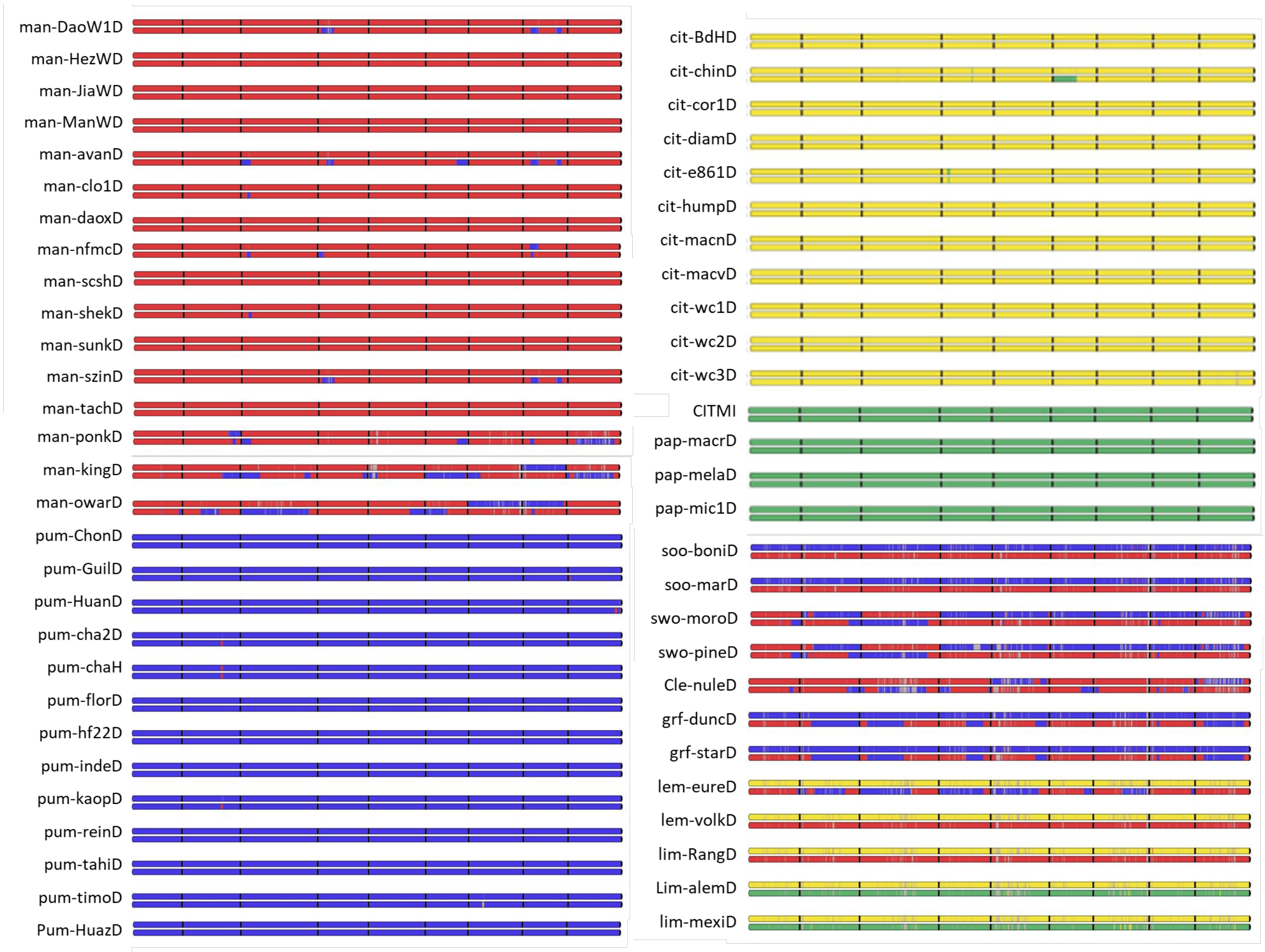

**Figure 18:** Interspecific mosaic of 55 citrus cultivars according CITMI genome assembly
red: *C. reticulata*; blue: *C. maxima*; yellow: *C. medica*; green: *C. micrantha*; grey: undetermined

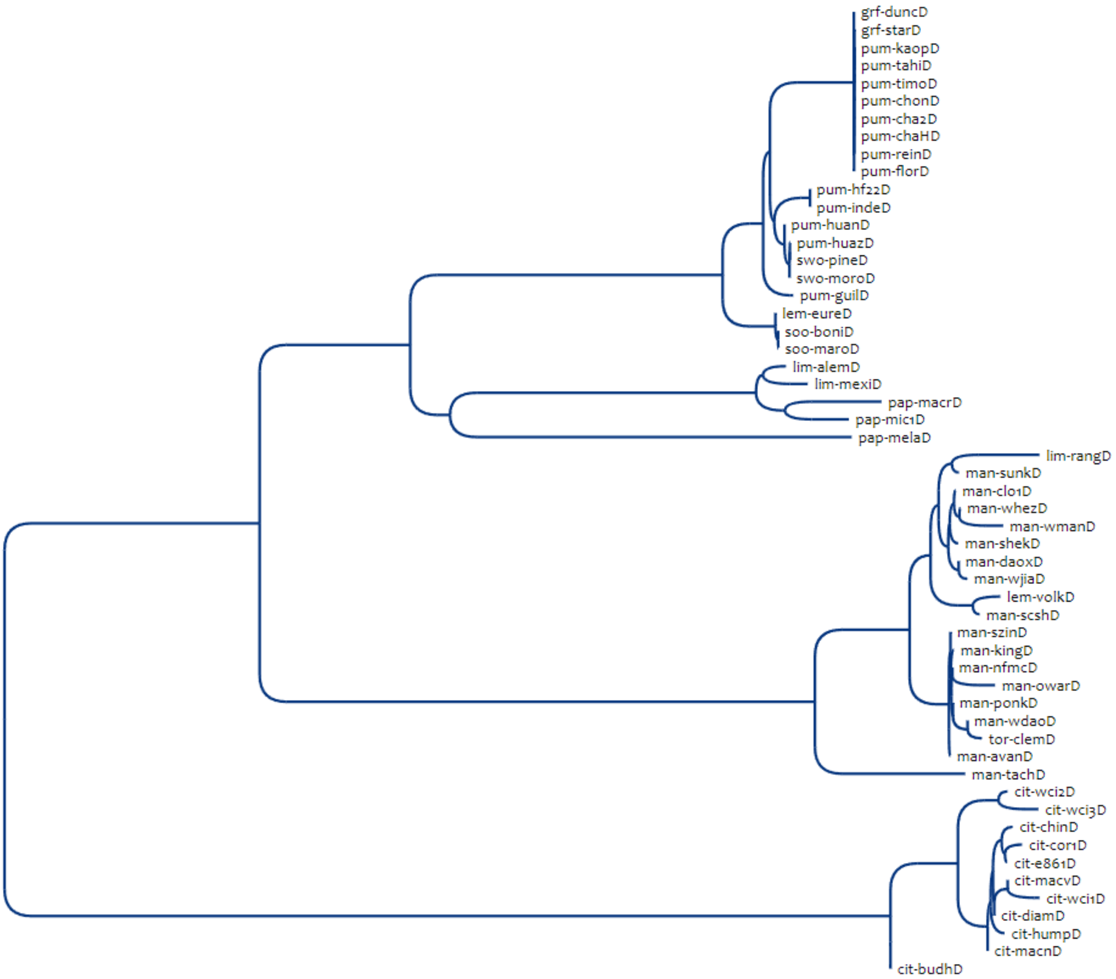

**Figure 19:** Chloroplastic phylogeny based on the minimum-evolution principle (Desper and Gascuel, 2002) from 1319 SNPs

Variant calling in *C. sinensis* chloroplast reference genome (Bausher et al., 2006)

#### 9) PAV analysis of the genes on the four genome assemblies

**Table 19:** Synthesis of PAV data on the four genome assemblies

| | | CITMI | CITME | CITRE | CITMA |
|---|---|---|---|---|---|
| Intraspecific diversity | Total Nb genes | 29258 | 28090 | 29477 | 30101 |
| | Core genes | 24847 | 25861 | 24548 | 24847 |
| | Dispensable genes | 3568 | 1953 | 4577 | 4691 |
| | Fully absent | 843 | 276 | 352 | 563 |
| | Specific to reference | 1112 | 8 | 138 | NA |
| Interspecific diversity | Core genes | 19718 | 20145 | 19920 | 20247 |
| | Diagnostic *C. reticulata* | 178 | 183 | 497 | 191 |
| | Diagnostic *C. medica* | 522 | 990 | 435 | 527 |
| | Diagnostic *C. maxima* | 119 | 117 | 92 | 295 |
| | Diagnostic *C. micrantha* | 210 | 148 | 134 | 135 |

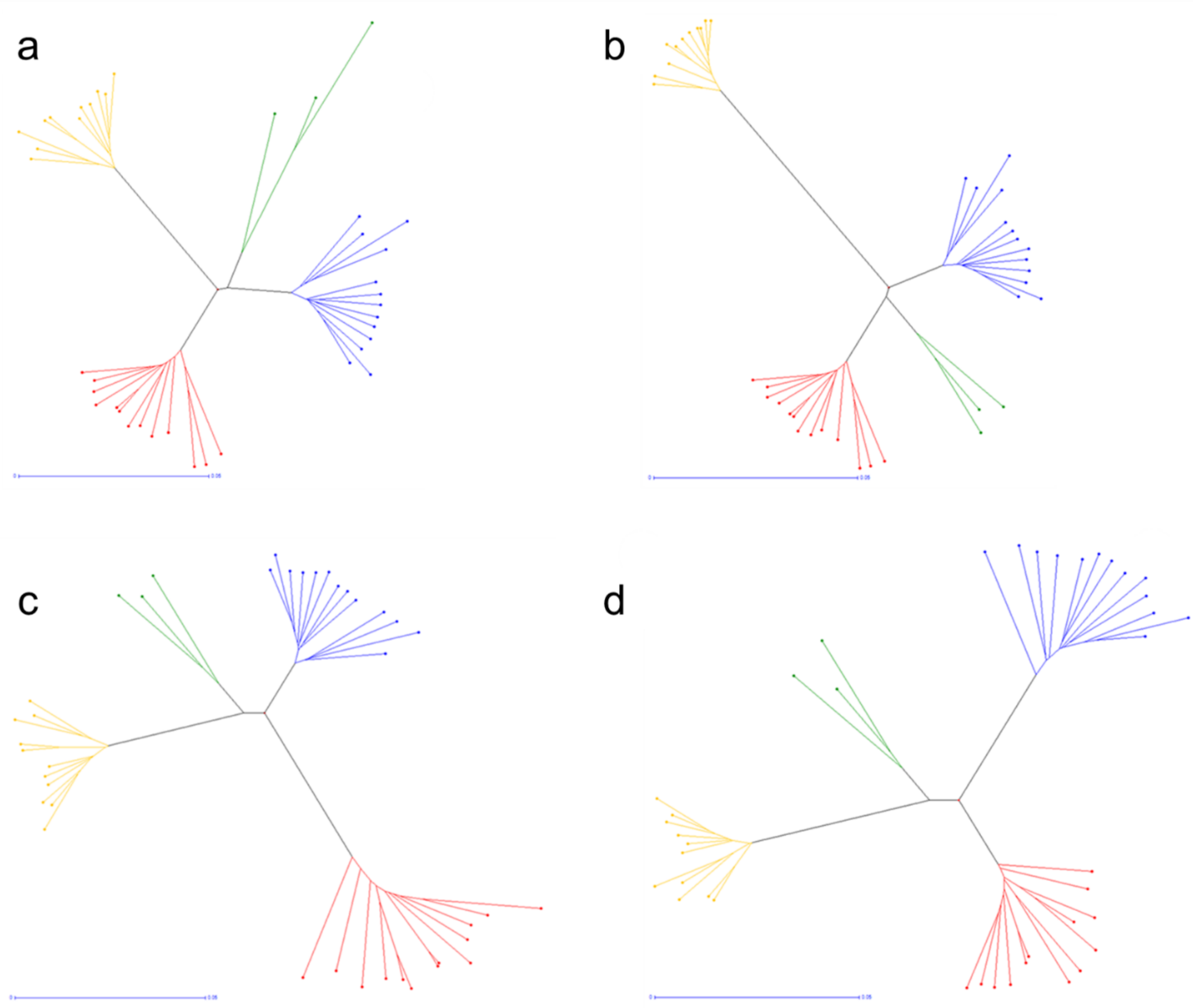

**Figure 20:** Neighbor joining trees of the accession representative of the four ancestors based on PAV.
a: CITMI reference genome (29258 genes); b: CITME (28090 genes); c: CITRE (29477 genes); d: CITMA (30101 genes). Red: mandarins, green: papedas, yellow: citrons; blue: pummelos.

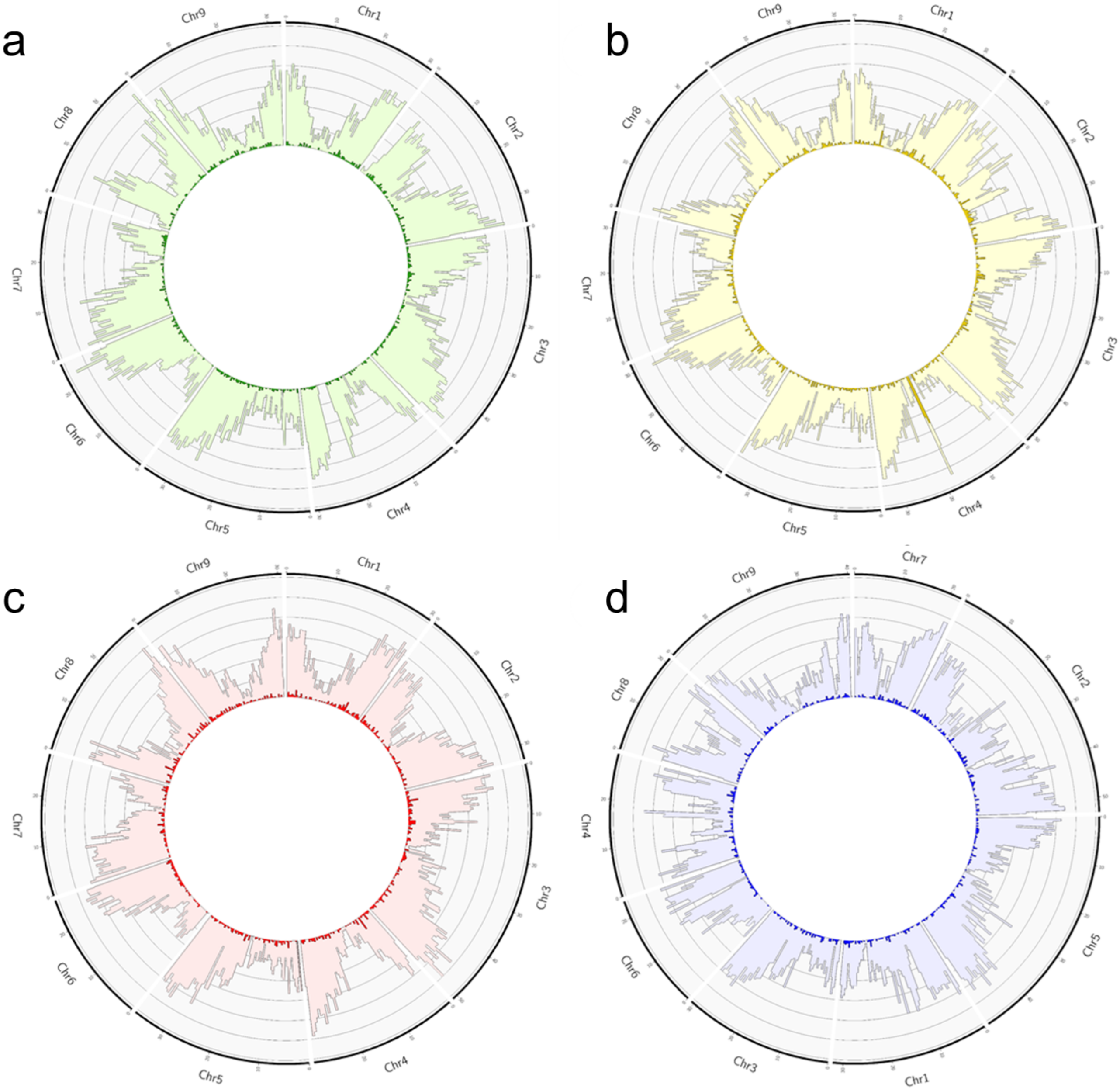

**Figure 21:** Distribution of the number of genes (soft color) and diagnostic genes (dark color) in the four genomes.

a: CITMI, b: CITME; c: CITRE; d: CITMA. Fixed windows of 500kb (scale 0-120 genes / 500kb). The numbering of the CITMA chromosomes is the one of Wang et al (2017) publication but for the figure they are ordered according to the other three genomes.

#### 10) Chloroplast insertion in CITME

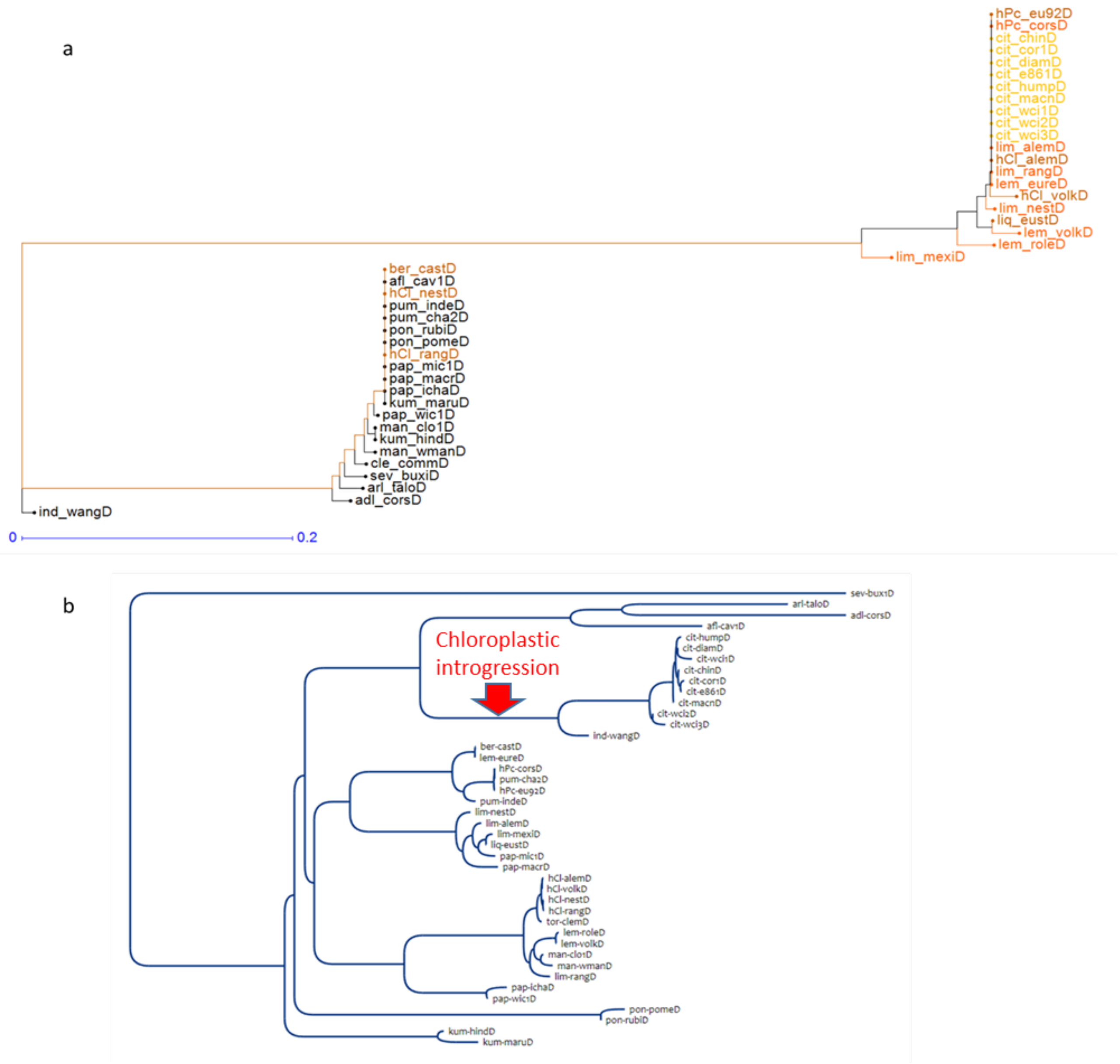

**Figure 22:** Classification of a citrus panel including additional *Citrus* species and hybrids of first and second generation of *C. medica* as male parent.

a: NJ tree based on PAV analysis for 97 CITME dPAV genes of the 17-18 Mb region of chromosome 4 Yellow: citron accessions, orange first generation hybrid of citrons; brown second generation hybrids of citrons; b: chloroplast phylogenetic tree based on the minimum-evolution principle (Desper and Gascuel, 2002) from 2255 SNPs (variant calling in *C. sinensis* chloroplast reference genome, Bausher et al., 2006).

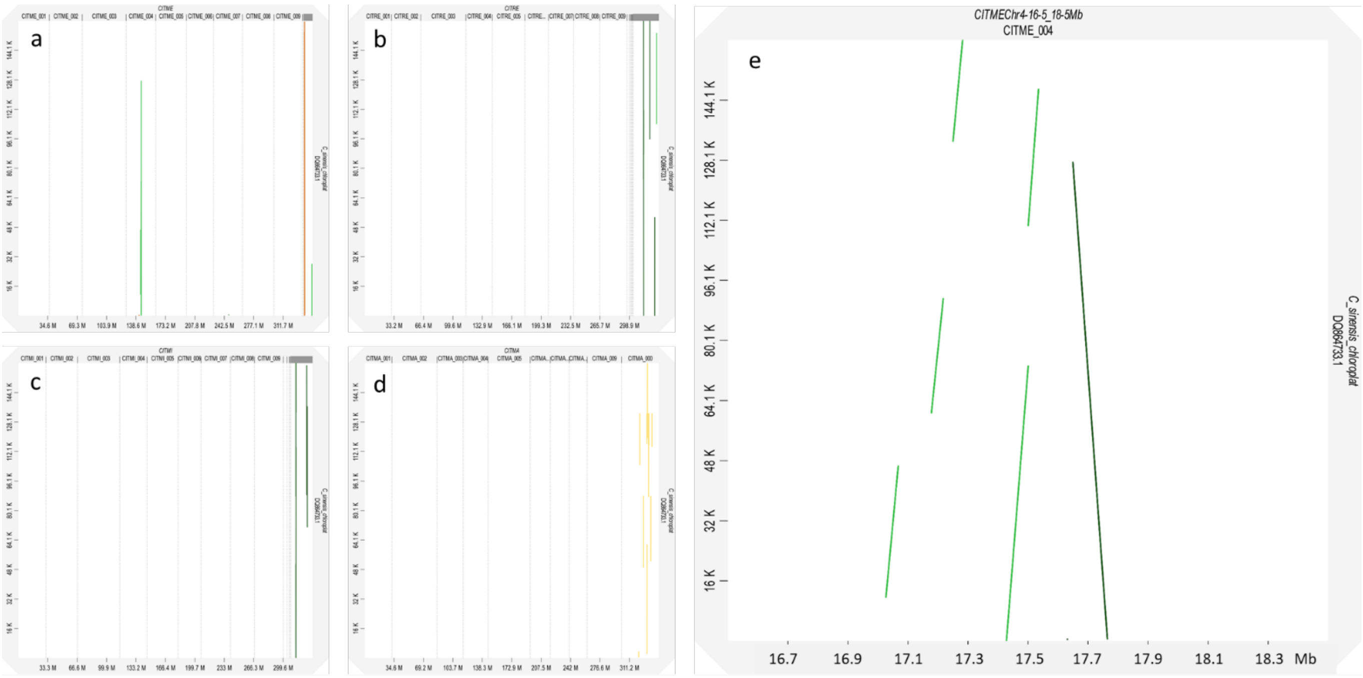

**Figure 23:** Dot plot analysis of the reference chloroplast genome from *C. sinensis* and CITME

(a), CITRE (b), CITMI (c) and CITMA (d) as genome target; d: dotplot of the chloroplast genome with the 16.5-18.5 Mb region of CITME Chr4 as target.

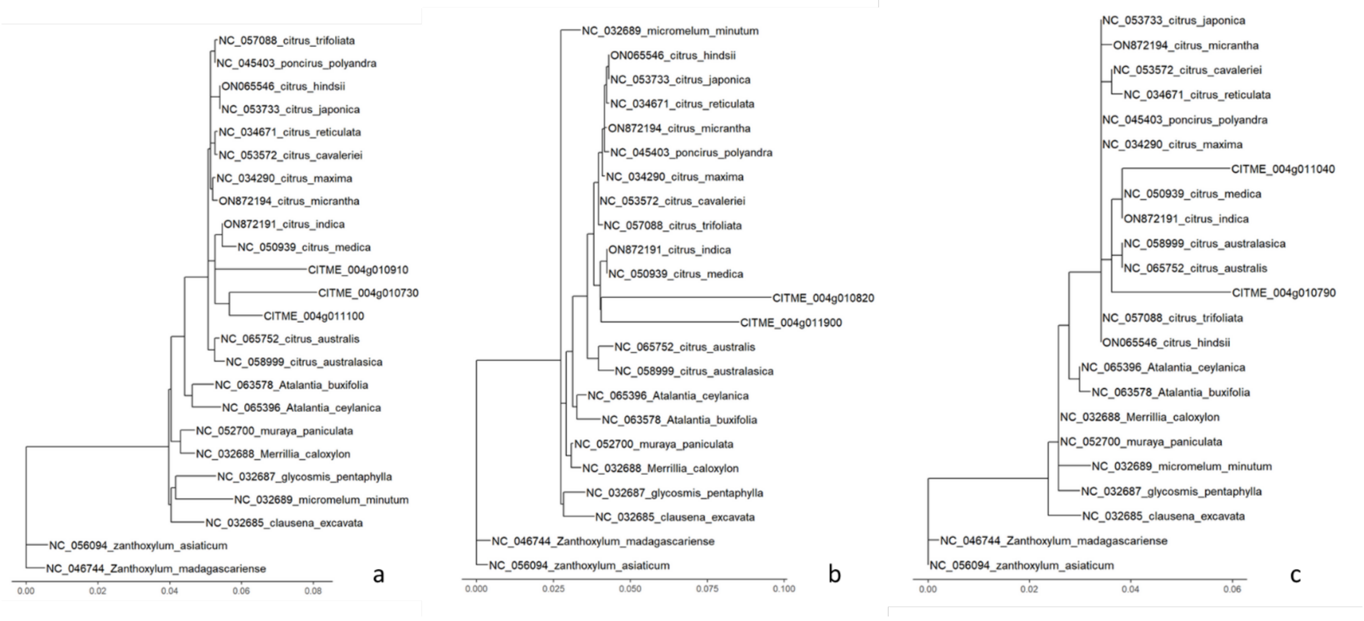

**Figure 24:** Phylogeny of several genes of the 17-18 Mb region of CITME Chr 4 orthologues of chloroplast genes and published chloroplast sequences.

a: MatK; b: rbcL; c: ndh J
Maximum likelihood analysis performed with phyML 3.0; (Guindon et al., 2010)

```
                           ....|....| ....|....| ....|....| ....|....| ....|....|
                                  10         20         30         40         50
CITME_matK          153+   MEEFQVVLEL DRSQQHDFLY PLLFREYIYV LAHDHGLNSS MMSLEGGFYD
CITME_004g010730      +    MEDFQVVLEL DRSQQHDFLY PLLFREYIYV LPHDHGLNSS MMSLEGGFYD
CITME_004g010910      +    XXXXXXXXXX XXXXXXXXXX XXXXXXXXXX XXXXXXXXXX MMSLEGGFYD
CITME_004g011100      +    XXXXXXXXXX XXXXXXXXXX XXXXXXXXXX XXXXXXXXXX XXXXXXXXXX

                           ....|....| ....|....| ....|....| ....|....| ....|....|
                                  60         70         80         90        100
CITME_matK          153+   NKSSSLSVKR LITRMYQRIN LSIAANDSNQ NPIFGHNNKL YSQIISEVFA
CITME_004g010730      +    NKSSSISVKR LITRMHQRIN LSIAANYSNQ NPIFGHNNKL YSQIISEVFV
CITME_004g010910      +    NKSSSLSVKW LITRMYQWIN LSIAANDSNQ SPIFGHNNKL YSQIISEVFA
CITME_004g011100      +    XXXXXXXXXX XXXXXXXXXX XXXXXXXXXX XXXXXXXXXX XXXXXXXXXX

                           ....|....| ....|....| ....|....| ....|....| ....|....|
                                 110        120        130        140        150
CITME_matK          153+   AGVEIPFSLR LVAFLEGKEI EKAPNFQSIH SIFPFFEDKL SHLNYVLDVR
CITME_004g010730      +    AVVEIPFSLR LVAFLEGIEI ESSPNFQSIH SIFPFFEDKL SHLNYVLDVR
CITME_004g010910      +    AVVEILFSLR LVAFLEGKEI EKSPNFQSIH SIFPFFEDKL SHLNYVLDVR
CITME_004g011100      +    XXXXXXXXXX XXXXXXXXXX XXXXXXXXXX XXXXXXXXXX XXXXXXXXXX

                           ....|....| ....|....| ....|....| ....|....| ....|....|
                                 160        170        180        190        200
CITME_matK          153+   IPYPICPEIL VQTLREWVKD ASSLHLLRFF LHEYFNSNSL ITPKNSISGF
CITME_004g010730      +    IPYPICPKIL VQTLREWVKD ASSLHLLRFF LHEYFNSNNL ITPKNSISGF
CITME_004g010910      +    MPYPICLEIL VQTLREWVKD ASSLHLLRFF LHKYFNSNSL ITPKNSISGF
CITME_004g011100      +    XXXXXXXXXX XXXXXXXXXX XXXXXXXXXX XXXXXXXXXX XXXXXXXXXX

                           ....|....| ....|....| ....|....| ....|....| ....|....|
                                 210        220        230        240        250
CITME_matK          153+   LKSNPRLLLF LYNSHVYEYE SILLFLCNQS SHLQSTSFRV LVERTYFYGK
CITME_004g010730      +    LKSNPRLLLF VYNSHVYEYE SILLFLCNQS SHLQSTSFRV LVERTYFYGK
CITME_004g010910      +    LKSNPRLLLF LYNSHVYEYE SILLFLCNQS SHLQSTSFRV LIERTYFYGK
CITME_004g011100      +    XXXXXXXXXX XXXXXXXXXX XXXXXXXXXX XXXXXXXXXX XXXXXXXXXX

                           ....|....| ....|....| ....|....| ....|....| ....|....|
                                 260        270        280        290        300
CITME_matK          153+   VEHLVEVFAT DFQDILGLVK DPFMHYVRYQ GKSILASKDT PLLMNKWKYY
CITME_004g010730      +    VEHLIEVFTK DFQDILGLVK DPFMHYVRYQ GKSILASKDM PLLMNKWKYY
CITME_004g010910      +    VEHLVEVFAR XXXXXXXXXX XXXXXXXXXX XXXXXXXXXX XXXXXXXXXX
CITME_004g011100      +    XXXXXXXXXX XXXXXXXXXX XXXMHYVRYQ GKSILASKDM PLLMNKWKYY

                           ....|....| ....|....| ....|....| ....|....| ....|....|
                                 310        320        330        340        350
CITME_matK          153+   LVGLWQWHFH ASSQPGRVQL NHLYLGKYAI NFLGYLSGVR LNSLLVRSQM
CITME_004g010730      +    LVGLWKWHFH ASSQPGRVQL NHLYLGMYAT NFLGYLSGVR LNSLLVRSQM
CITME_004g010910      +    XXXXXXXXXX XXXXXXXVQL NHLYLGKYAI NFLGYLFGMR LNSLLVLSQM
CITME_004g011100      +    LVGLWQWYFH ASSQPRRVQL NHLYLGKYAI NFLGYLSGVR LNSLLVRSQM

                           ....|....| ....|....| ....|....| ....|....| ....|....|
                                 360        370        380        390        400
CITME_matK          153+   LENSFLIDNS MKKVDTTVPI IHLIGSLTKA RFCNALGHPI SKSTWSDFSD
CITME_004g010730      +    LENSFIIDNS MKKVDTTIPI IHLIGSLTKA RFCNALGHPI SKSTWADFSD
CITME_004g010910      +    LENSFLIDNS MKKVDTTIPI IHLIGSLTKA RFCNALGHPI SKSTWADFSD
CITME_004g011100      +    LENSFIIDNS MKKVDTTVPI IHLIGSLTKA RFCNALGHPI SKSTWADFSD

                           ....|....| ....|....| ....|....| ....|....| ....|....|
                                 410        420        430        440        450
CITME_matK          153+   SHLIDRFVRI CRNLSHYYSG SSKKKSLYRV KYILRLSCVK SLVRKHKSTV
CITME_004g010730      +    SHLINRFVRI CRNLSHYYNK SLKKKXLYRV .......... ..........
CITME_004g010910      +    SHLIDRFVRI CRNLSHYYSG SSKKKSLYRV KYIFRLFCVK TLVREHKSIV
CITME_004g011100      +    SHLIDRFVRI CRNLSHYYSG SSKKXXXYRV KFILRLSCVK SLVRKHKTXX

                           ....|....| ....|....| ....|....| ....|....| ....|....|
                                 460        470        480        490        500
CITME_matK          153+   RAFLKRLGSE LLEEFLMEEE HVLALLFPGA SSTSRRFYLY RGRIWYLDIF
CITME_004g010730      +    .......... .......... .......... .......... ..........
CITME_004g010910      +    RAFLKRLGSE LLEEFLTEEE HVFAFLFPGA SSTSCRFYLY RGRIWYLDIF
CITME_004g011100      +    XXXXXXXXXX XXXXXXXXXX XXXXXXXXXX XXXSRRFYLY RGQIWYLDIF

                           ....|....|
                                 510
CITME_matK          153+   CINDLVNYQ*
CITME_004g010730      +    ..........
CITME_004g010910      +    CINDLVNYQ*
CITME_004g011100      +    CINDLVNYQ*
```

**Figure 25:** Alignment of nuclear sequence of nuclear and chloroplastic ortologues of MatK gene

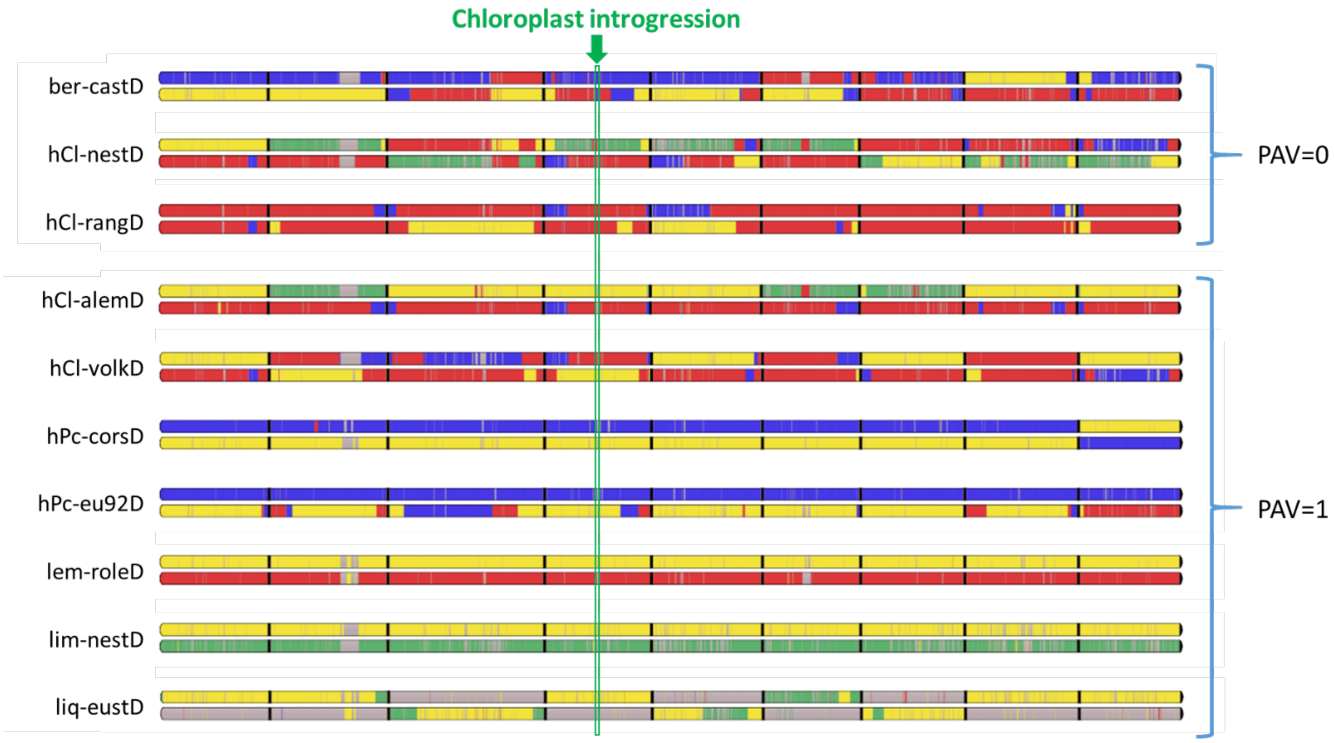

**Figure 26:** Relation between the contribution of C. medica nuclear genome in the 17-18 Mb region of Chr 4 and the presence of the supposed introgressed chloroplast genes in additional hybrids of first and second generation with *C. medica* male parent.

red: *C. reticulata*; blue: *C. maxima*; yellow: *C. medica*; green: *C. micrantha*; grey: undetermined; for Eustis limequat (liq-eustD) the grey area corresponds to *C. japonica* genome.

#### 11) Pangene results

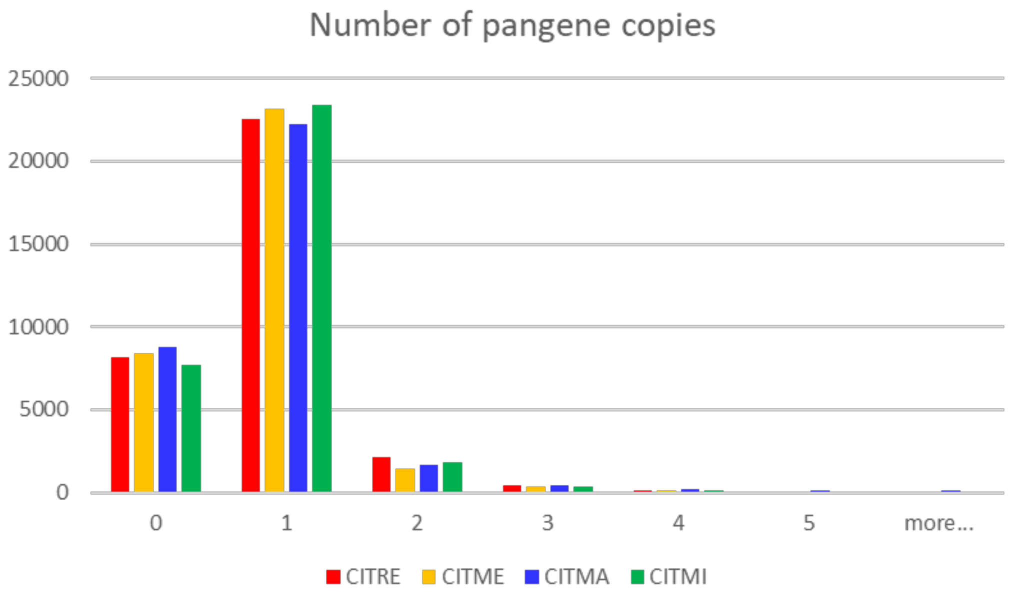

**Figure 27:** Number of pangene copies in the four ancestral genome assemblies

**Table 21**: Constitution of the specific and *Citrus* pangenome

| | *C. micrantha* | *C. medica* | *C. reticulata* | *C. maxima* | Super-pangenome |
|---|---|---|---|---|---|
| Core | 29156 | 28689 | 28843 | 28736 | 25291 |
| Soft-Core | 1521 | 522 | 723 | 809 | 2431 |
| Dispensable | | 1269 | 1930 | 2122 | 5171 |
| Private | 858 | 216 | 209 | 182 | 69 |
| Absent | 1427 | 2266 | 1257 | 1113 | 0 |
| Total pangenes | 31535 | 30696 | 31705 | 31849 | 32962 |

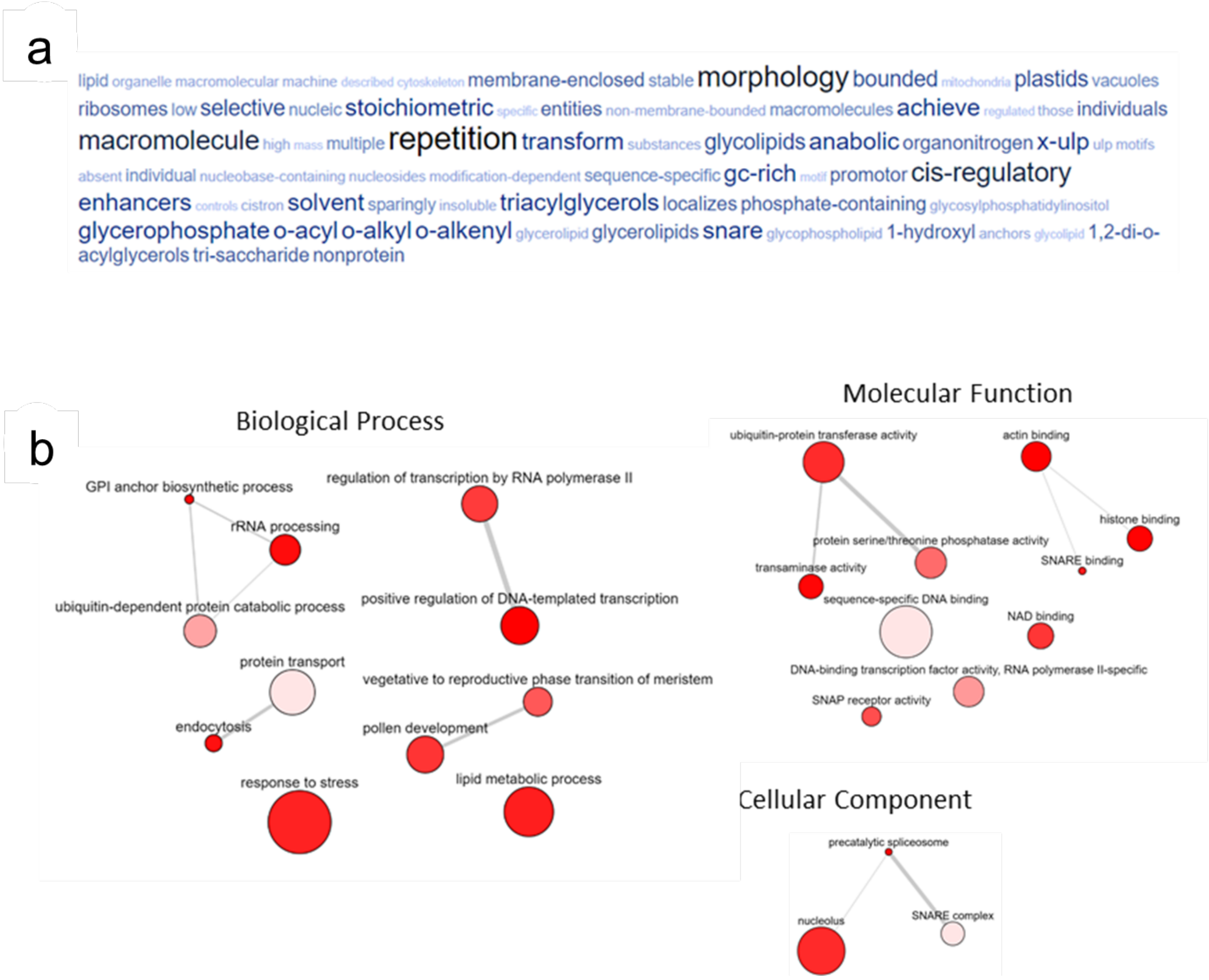

**Figure 28:** Core genome functional enrichment analysis

a) Functional enrichment analysis of the core genome. Graphical representation of enriched biological process (GOs). Size of the words and color contrast are proportional to their representativeness in the gene pool. b) Integrative map, generated using REVIGO, visually represents the enriched Gene Ontology (GO) terms identified in the citrus core genome. Each point on the map corresponds to a GO term, with the distance between points indicating the semantic similarity between the terms. Clusters of related GO terms are color-coded to reflect their association with specific biological processes, molecular functions, or cellular components. Larger points represent GO terms with greater statistical significance, indicating a higher level of enrichment.

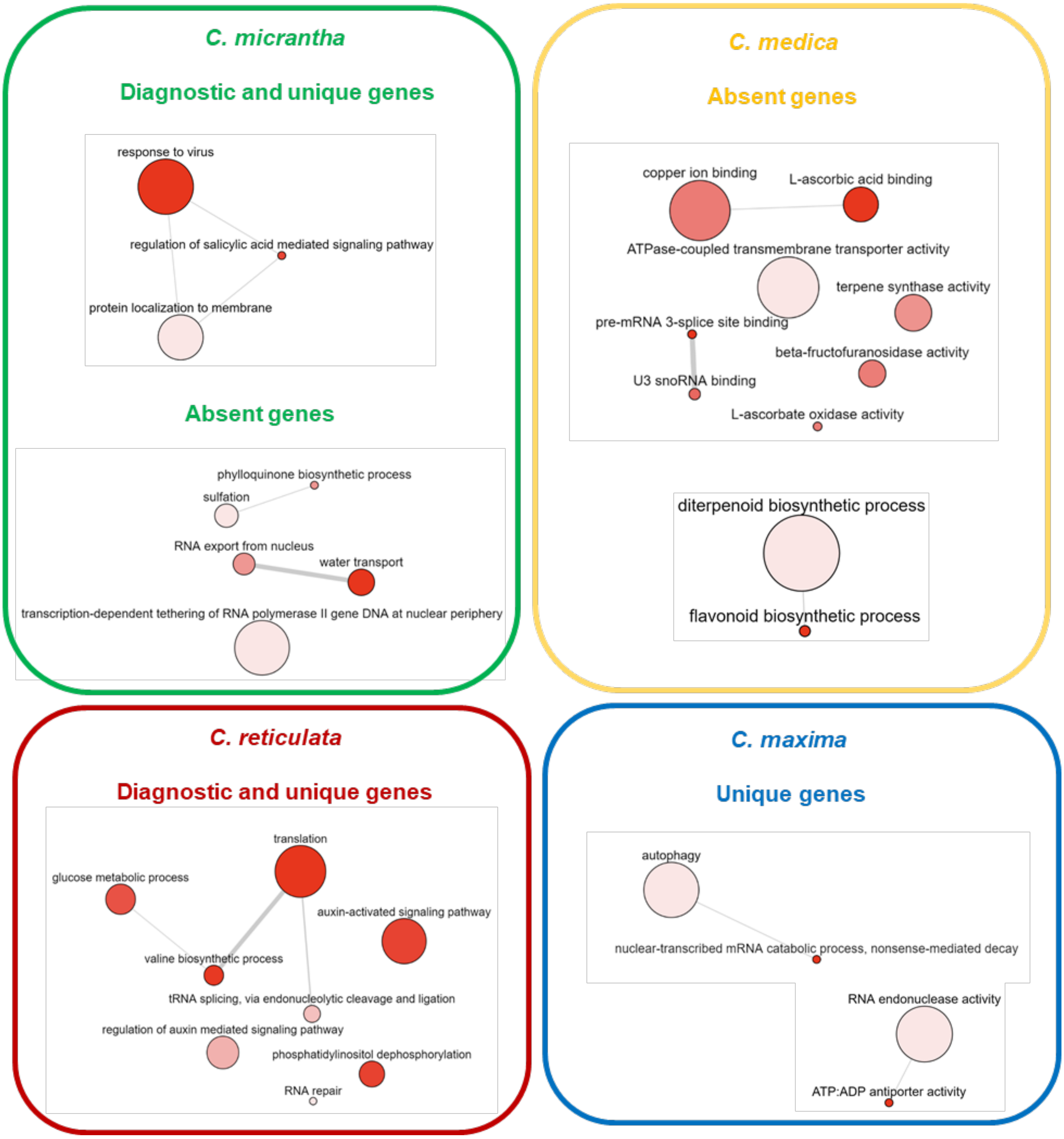

**Figure 29:** Integrative REVIGO Maps of Enriched GO Terms for the four Citrus Ancestral Citrus Species.

Integrative maps generated using REVIGO, summarizing the enriched Gene Ontology (GO) terms for the unique and diagnostic genes and/or absent genes of each citrus species within the super-pangenome: *C. reticulata*, *C. maxima*, *C. medica*, and *C. micrantha*. Each panel represents a species-specific integrative map, where individual points correspond to enriched GO terms, and their placement reflects semantic similarities. Larger points indicate higher statistical significance of enrichment. These maps provide a comparative overview of the distinct biological processes and molecular functions enriched in each species, illustrating their unique adaptive traits and differences in functional specialization. The maps collectively underscore the evolutionary diversification within the citrus pangenome, while also highlighting the shared core processes across the species.
